# Genomic basis of circannual rhythm in the european corn borer moth

**DOI:** 10.1101/633362

**Authors:** Genevieve M. Kozak, Crista B. Wadsworth, Shoshanna C. Kahne, Steven M. Bogdanowicz, Richard G. Harrison, Brad S. Coates, Erik B. Dopman

## Abstract

Genetic variation in life-history timing allows populations to synchronize with seasonal cycles but little is known about the molecular mechanisms that produce differences in circannual rhythm in nature. Changes in diapause timing in the European corn borer moth (*Ostrinia nubilalis*) have facilitated rapid response to shifts in winter length encountered during range expansion and from climate change, with some populations emerging from diapause earlier to produce an additional generation per year. We identify genomic variation associated with changes in the time spent in winter diapause and show evidence that the circadian clock genes period (*per*) and pigment dispersing factor receptor (*Pdfr*) interact to underlie this adaptive polymorphism in circannual rhythm. *Per* and *Pdfr* are located within two epistatic QTL, strongly differ in allele frequency among individuals that pupate earlier or later, have the highest linkage disequilibrium among gene pairs in the QTL regions despite separation by > 4 megabases, and possess amino-acid changes likely to affect function. One *per* mutation in linkage disequilibrium with *Pdfr* creates a novel putative clock-cycle binding site found exclusively in populations that pupate later. We find associated changes in free-running daily circadian rhythm, with longer daily rhythms in individuals that end diapause early. These results support a modular connection between circadian and circannual timers and provide testable hypotheses about the physiological role of the circadian clock in seasonal synchrony. Winter length is expected to continually shorten from climate warming and we predict these gene candidates will be targets of selection for future adaptation and population persistence.

## INTRODUCTION

Many species display tremendous flexibility in the annual timing of physiological, morphological, and behavioral transitions that enable survival in seasonal environments. The capacity to adjust the timing of circannual rhythms and track local seasonal cycles can facilitate expansion into new geographic areas with different seasonal environments (1, 2). Dramatic shifts in seasonal timing seen across plants and animals in recent decades (3) may also enable adaptation (4) and persistence (5, 6) during rapid, anthropogenic alterations of the environment. When shifts in seasonal timing additionally change the number of generations per year (7-9), populations may grow faster and even tolerate a faster rate of sustained environmental change (Figure S1) (10). Nevertheless, if environments change too rapidly, timing mismatches with seasonality can occur (11, 12), resulting in loss of fitness and population decline (5, 6, 12). The capacity to adjust seasonal timing and track changes in seasonal cycles, as well as our ability to evaluate risks to biodiversity, depends in part on the proximate causes of variation in circannual rhythm (10, 13). However, relatively little is known about the molecular basis of this diversity (11, 14, 15).

Insects are highly variable in the seasonal timing of transitions into and out of diapause, a stress-tolerant physiological state that enables coping with seasonal challenges (1, 16-19). For temperate species, which generally use seasonal changes in photoperiod and temperature to synchronize diapause with winter stress, the timing of diapause transitions in spring and autumn varies widely within and among species (15, 20) and therefore provides an excellent opportunity for analysis of the genetic control of circannual timing. We used natural variation in timing of spring transitions from larval diapause to active development of the European corn borer moth (*Ostrinia nubilalis*) to understand the genetic basis of this seasonal adaptation. In corn borers, the duration of developmental arrest in the spring (i.e., diapause termination timing, (18)) generally tracks winter length. Shifts in diapause timing have rapidly evolved across latitudinal gradients in winter after range expansion from Europe to North America in ∼1910 (21, 22). The length of winter decreases with decreasing latitude in North America. Consequently, the optimal time to exit diapause is advanced to earlier in the year and the number of days required to end larval diapause evolved into a positive correlation with latitude (ranging from 17.5 to 49 days across 9.28°N latitude; Figure S2) (22). Changes in diapause timing similarly tracks shorter winters associated with climate warming, with populations at the same latitude showing a ∼50% average reduction in time needed to transition out of diapause since the 1950s in some locations (22, 23). Although broad-scale changes in climate may be leading to directional change in diapause timing across space and over time, polymorphism is common in natural populations (24, 25) and may be maintained (26) because timing shifts alleviate competition for limiting resources (such as host plants) or may be favored by year-to-year fluctuations in seasons (27). In the mid-Atlantic region of the United States, earlier springtime pupation (∼20 days) reduces generation time, thereby enabling production of two generations per year (bivoltine) rather than one generation per year (univoltine; Figure S3) (28, 29). Geographic co-occurrence of earlier-and later-pupating individuals (∼40 days) leads to asynchronous adult mating flights (June versus July) and allochronic reproductive isolation (Figure S4) (29). Thus, in corn borers, range expansion and population growth, enhanced tolerance of environmental change, and speciation may all be byproducts of natural selection on diapause timing during seasonal adaptation (29, 30).

Despite decades of work on *Ostrinia* (25, 26, 31-33), no causative loci for natural variation in diapause termination timing have been identified definitively. We have therefore taken an unbiased, whole-genome approach to identify loci underlying this variation. Previous work found sex-linked inheritance (31, 32) and evidence for a single quantitative trait locus (QTL) on the Z (sex) chromosome (25). However, a putative inversion encompassing 39% of the Z chromosome was discovered and the QTL was locked into a non-recombining region along with hundreds of genes, many of which were differentially expressed during diapause break (33-35). Subsequent work demonstrated that the recombination suppressor is polymorphic in field populations (35) and we therefore performed QTL mapping in pedigrees with putatively collinear Z chromosomes. An advantage of this approach is that it will exclusively identify the genetic architecture of diapause timing, but a challenge is that individual genes or mutations associated with trait differences cannot be easily identified. Therefore, we obtained the higher resolution needed using population genomic sequencing data derived from phenotyped, field-caught moths. As population genomic analysis will identify mutations within individual genes underlying changes in measured traits, as well as loci controlling unmeasured traits subject to correlated selection in nature (such as temperature tolerance), we view the approaches as parts of a complementary, two-pronged forward-genetics strategy to characterize the genetics of natural variation in circannual rhythm.

## RESULTS

### QTLs for diapause timing

Our first forward genetic approach used QTL mapping of diapause termination timing. In corn borers, the main environmental trigger to end diapause is photoperiod (36, 37). The time required to end diapause and return to active development was quantified as the time to pupation under diapause-breaking photoperiod and temperature (referred to as post-diapause development (PDD) time). Variation in PDD time was measured in backcross pedigrees of females from a two-generation (bivoltine), early-emerging, short-PDD population collected in East Aurora, NY in 2011 (EA, PDD time < 19 days) and males from a one-generation (univoltine), later-emerging, long-PDD laboratory colony originally derived from Bouckville, NY in 2004 (BV, PDD time ≥ 39 days) (25, 28, 34, 35, 38). Diapause in 5^th^ instar backcross larvae was induced by a winter-like short-day 12 hour (h) photoperiod. Subsequently, PDD time was measured as the number of calendar days required for diapausing larvae to pupate after transfer to a summer-like long-day (16 h) photoperiod. PDD time varied from 11 to 63 days. Using 167 autosomal and 18 Z-linked molecular markers, we found that only the Z chromosome was significantly associated with PDD time (*N* = 67 offspring, LOD = 9.67, P < 0.001).

We refined the Z chromosome QTL map by including 226 offspring and 35 markers which resulted in the prediction of two adjacent, interacting QTL (Figure 1a). Model fit was improved by inclusion of QTL1 (*F*_2_ = 22.89, *P* < 0.001), QTL2 (*F*_2_ = 9.09, *P* < 0.001), and their interaction (*F*_1_ = 7.80, *P* = 0.005; Table S1; Figure 1c,e), indicating that natural variation for diapause termination time in the analyzed populations is regulated by at least two genes. Together these QTL and their interaction explained 35.3% of phenotypic variance. QTL1 was located at 29.5 cM (*BCI* = 24.5-31 cM) (Figure 1b) and QTL2 was located at 34 cM (*BCI* = 31.5-36 cM) (Figure 1c,d). QTL1 was estimated to be at ∼3.1 Mb in size (on the 21 Mb Z chromosome), containing ∼48 genes with annotations in draft European corn borer moth genome (GenBank BioProject: PRJNA534504; Accession SWFO00000000; Supplemental Material; Table S2). QTL2 was estimated to be ∼3.7 Mb in size with ∼42 annotated genes. The QTL1 allele originating from the parental strain expressing shorter PDD time (EA) was epistatic to allelic changes at QTL2 and masked its effect (two-way ANOVA, *F*_1,218_ = 8.76, *P* = 0.003; Figure S5). In contrast, an allele at QTL1 originating from the parental strain expressing longer PDD time (BV) resulted in an unusually long PDD time when paired with a QTL2 allele from a short-PDD time parent (EA) (mean PDD time ± SD: QTL1_BV_/QTL2_EA_ = 51 ± 11.92, QTL1_BV_/QTL2_BV_ mean PDD time = 39.52 ± 8.05).

**Figure 1.**
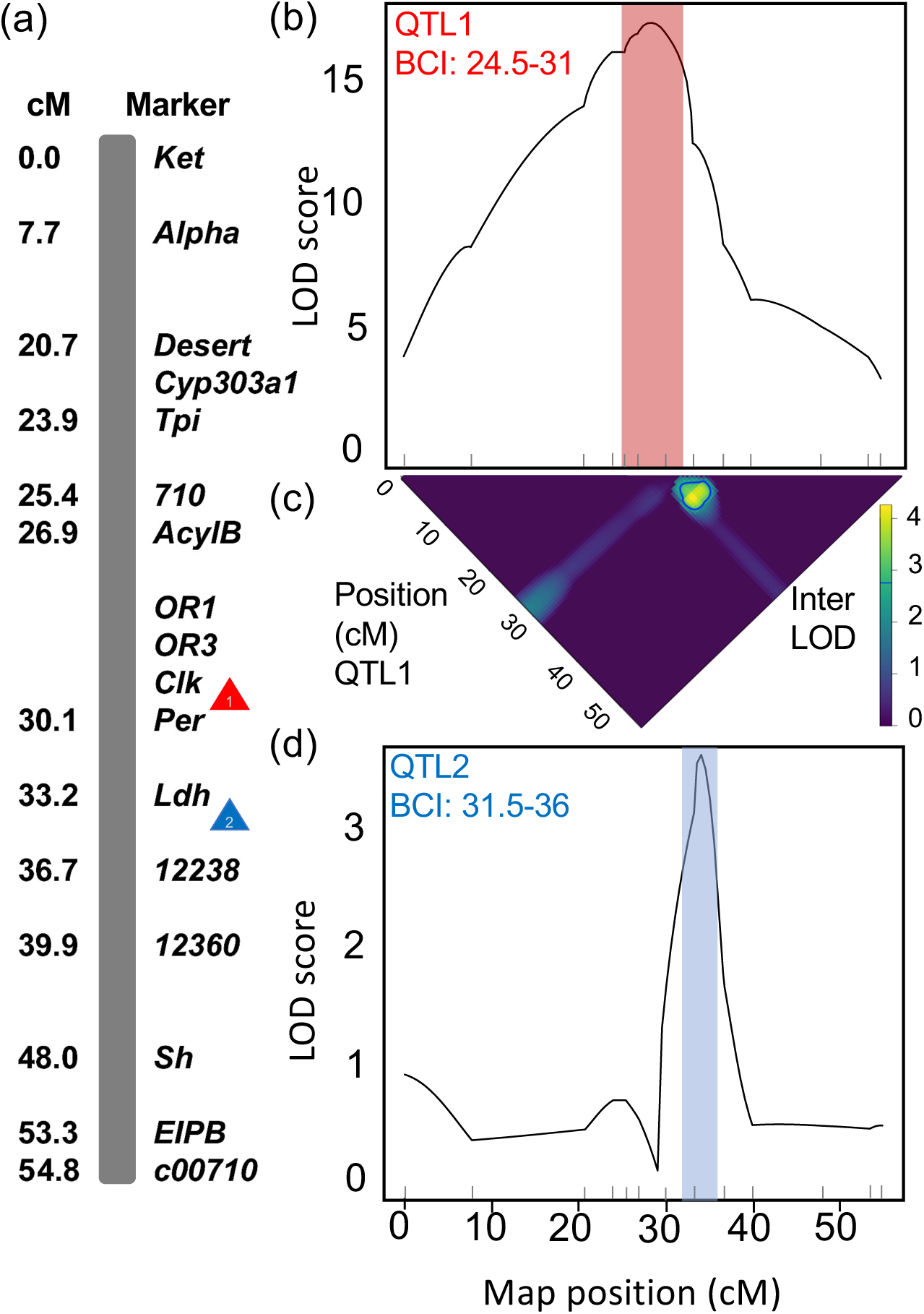
QTL for PDD time on the Z chromosome. a) Z chromosome consensus linkage map, adapted from Kozak et al. (2017) (35); b) plot of QTL1 estimated using scanone, with Bayesian credible interval (BCI) shaded; c) plot of the LOD score for a model with an epistatic interaction compared to a model including only a single QTL, blue line indicates the 1.5 LOD interval contour and the location of QTL2; d) plot of QTL2 estimated using scanone on individuals with slow allele at QTL1 (BCI shaded). N = 226 offspring for a-c; N = 101 for d.

### Genetic variation in natural populations

We used genome sequencing of wild populations as a second forward genetic approach to identify segregating chromosomal regions affecting diapause timing. We tested for associations with PDD time in pooled sequencing (pool-seq) data derived from field collections of 5 populations (Table S3). In addition to a pool of a pool of individuals from the EA population (*N* = 34; mean PDD time = 9.8 ± 1.2 days) and a separate field collected pool from the BV population (*N* = 20; mean PDD time = 42.9 ± 7.9 days), we sequenced pools from Penn Yan, NY (PY, *N* = 26; univoltine, long; mean PDD time = 52.9 days ± 5.3), Geneva, NY (GEN, *N* = 25; univoltine, long; mean PDD time = 45.3 days ± 5.4), and Landisville, PA (LA, *N* = 39; bivoltine, short; PDD time < 19 days). Paired-end 150 bp Illumina reads were aligned to the draft European corn borer moth genome ordered and oriented into 31 chromosomes (61% of 454.7 Mb assigned locations genome-wide; 20.9 Mb of the Z chromosome ordered; Table S4). Average coverage was 25X (range = 12-40X).

Overall genetic differentiation was low among populations as estimated from mean pairwise *F*_*ST*_ across 1 kb windows (mean autosomal *F*_*ST*_ = 0.05; Z chromosome *F*_*ST*_ = 0.06) with the highest values of *F*_*ST*_ between short and long populations observed on the Z chromosome in the QTL regions (Figure 2, Figure S6). To identify gene regions associated with PDD time while directly accounting for population demography we used a Bayesian framework. BayPASS 2.1 (39) was used to estimate a covariance matrix that represents an approximation of the unknown demographic history and test associations of single nucleotide polymorphisms (SNPs) with PDD time while accounting for any covariance. BayPASS was run separately for Z chromosome and autosomal loci, as our expectations about the demographic history and number of haploid chromosomes in each pool differed between sex chromosomes and autosomes (see methods). Significantly associated alleles were defined as containing SNPs with a Bayes Factor (BF) > 20 deciban units (*dB*), correlation coefficient (*r*) ≥ 0.5, and strength of association (β) with a posterior distribution that had a probability < 0.01% of β = 0 (“empirical Bayesian P-value” *eBP*_*is*_ > 2) (39, 40). The autosomal analysis identified 7 SNPs in predicted genes with BF > 20 *dB*, but all of these had *eBP*_*is*_ < 0.5. On the Z chromosome, 16 of 8,435 SNPs in predicted genes had BF > 20 *dB* and showed a strong association with PDD time (*r* ≥ 0.57). However, only four Z-linked SNPs (0.05%) in predicted genes had *eBP*_*is*_ > 2. Congruent with our mapping results, all SNPs with *eBP*_*is*_ > 2 fell inside the two interacting QTL regions (Table 1a; Figure 2c) and three were within two genes known to interact in the same pathway— the circadian clock genes period (*per*) and pigment-dispersing factor receptor (*Pdfr*). One SNP was within QTL1 in *per* (Figure 3a; Table S5), a core circadian clock gene (41), and two SNPs within QTL2 were in *Pdfr*, the gene encoding the receptor for the main circadian neurotransmitter PDF (42, 43). The remaining SNP with *eBP*_*is*_ > 2 was within terribly reduced optic lobes (*trol*) (Figure 3b) which encodes an extracellular matrix protein and is not known to interact with *per* (44). Two additional intergenic SNPs with *eBP*_*is*_ > 2 were between *Pdfr* and *trol* (Figure 3b). No additional outlier loci were detected when we analyzed all genome scaffolds (including those lacking an assigned chromosomal location) as if they were on the Z chromosome, indicating that *per* and *Pdfr* have the strongest association with PDD time across the entire sequenced genome (Figure S7-S8). *Per* and *Pdfr* also displayed extreme values of *F*_*ST*_ (> 0.5) and significance (q < 10^−10^) in Cochran-Mantel-Haenszel (CMH) outlier tests (45) (see supplemental results; Table S6; Figure S9).

**Table 1.**
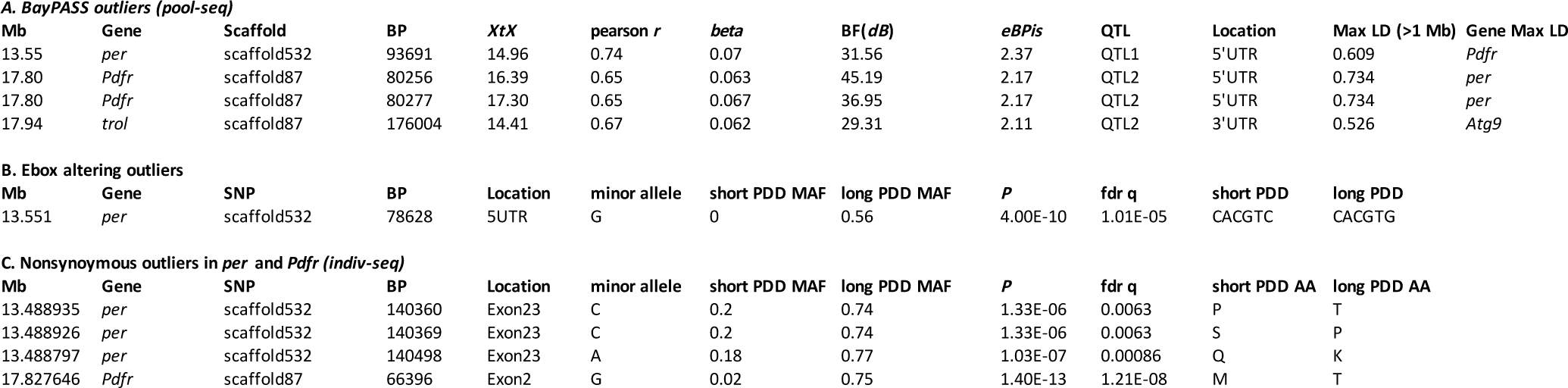
Locations of outlier SNPs in PDD QTL. a) SNPs identified by BayPASS in pool-seq data including the demographic corrected measure of population differentiation (*XtX*), and measures of association with PDD time: pearson *r, beta*, Bayes Factor (BF, measured in *dB*), *eBP*_*is*_, and maximum linkage disequilibrium with other SNPs under the QTL. Outliers in the individual resequencing data case/control analysis (*P* values and FDR corrected q-values shown) with long PDD allele (minor allele) and minor allele frequency (MAF) for short and long PDD samples for b) E-box altering SNP and c) amino-acid (AA) altering SNPs. All case-control results listed in Table S9-S10. Position in base pairs (BP) on the scaffold shown.

**Figure 2.**
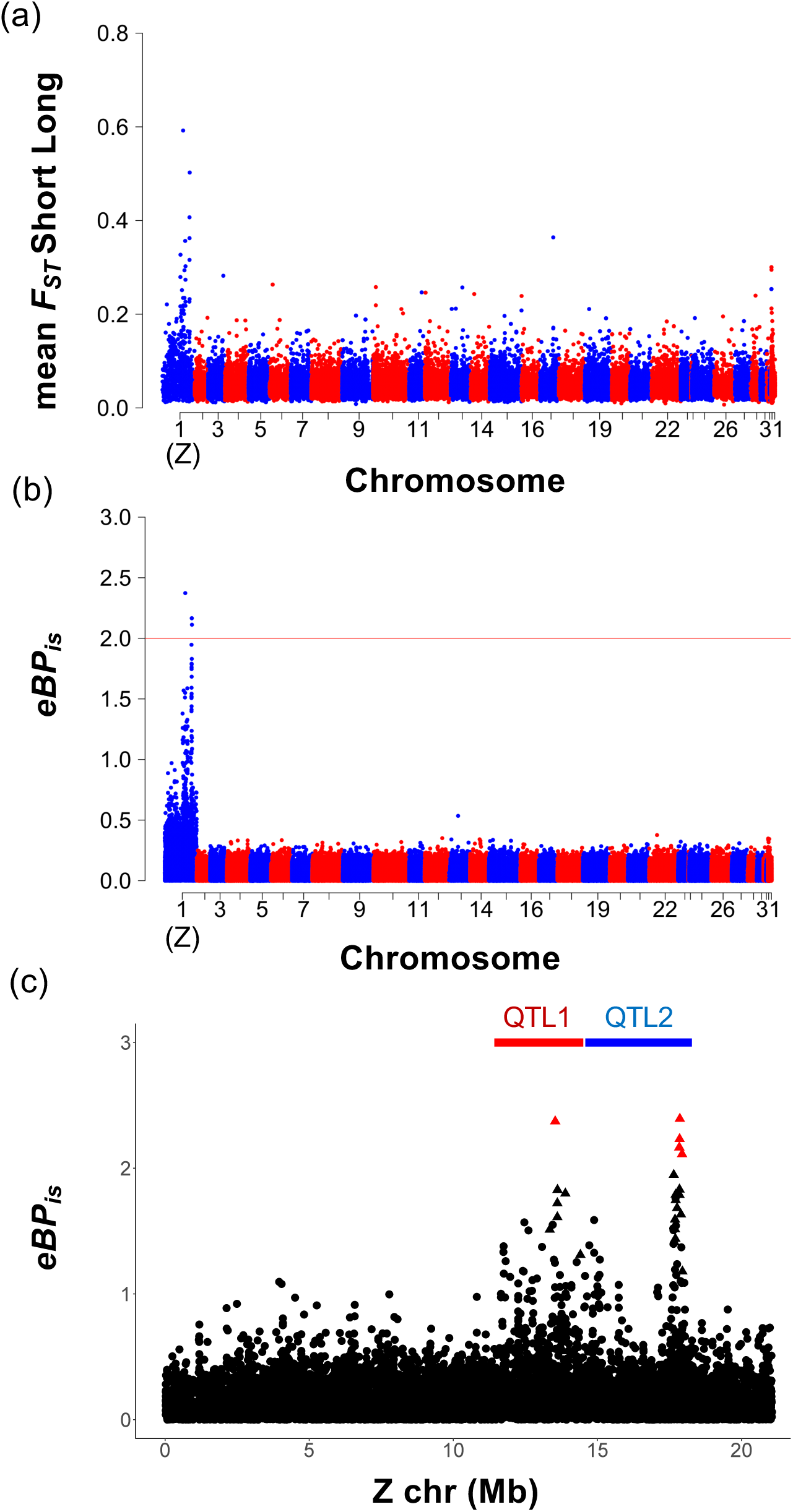
SNP association with PDD time in natural populations. a) Genome-wide plot of mean *F*_*ST*_ between short and long populations gene regions for chromosomes 1(Z)-31, with unscaffolded chromosomes assigned to chromosome 32; N = 57,842 1 kb windows. (b) Genome-wide plot of BayPASS empirical Bayesian P-values (*eBP*_*is*_) for PDD association (N = 293,590 SNPs in CDS); *eBP*_*is*_ > 2 (equivalent to *P*< 0.01) indicated by red line. (c) Plot across the Z chromosome. SNPs with strongest evidence of association denoted by triangles (Bayes Factor > 20 *dB*) and labeled in red (*eBP*_*is*_ > 2); no evidence denoted by black circles (BF < 20 *dB*); location of QTL1 and QTL2 BCI shown; N = 8,435 SNPs within genes.

**Figure 3.**
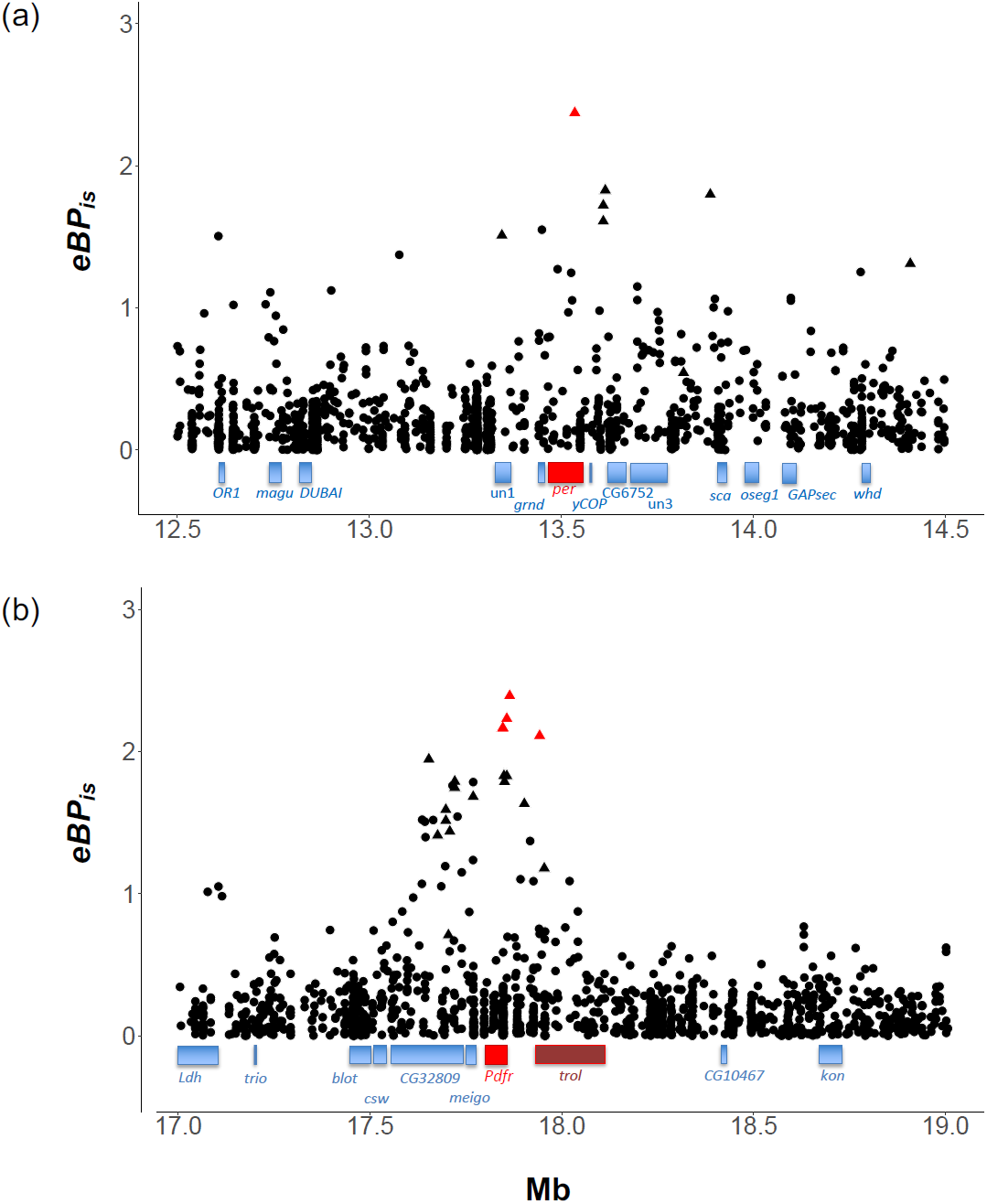
Gene locations relative to peak association with PDD time. a) *eBP*_*is*_ plotted for 2 Mb interval around *per*, showing *per* (red) location and other flanking gene intervals (blue); entire region is within QTL1 BCI. Intergenic SNPs included (N=1,441). b) *eBP*_*is*_ plotted for 2 Mb interval around *Pdfr* (bright red) location, *trol* (dark red) and other gene intervals (blue); all within QTL2 BCI except *kon* and CG10467 (N=1,423 SNPs). *eBP*_*is*_ > 2 labeled in red. BF > 20 *dB* denoted by triangles. Full gene descriptions listed in Table S5.

Linkage mapping indicated two QTL located ∼4.5 cM (∼ 5 Mb) apart contribute to the evolution of PDD time. In wild populations, alleles in QTL1 and QTL2 associated with PDD time should be in high linkage disequilibrium (LD) due to their joint effect on the phenotype. To measure LD, we resequenced the genomes of individual moths from all 5 populations with long (*N* = 18) and short (*N* = 25) PDD times using 150 bp paired-end Illumina sequencing at 14X coverage (mean coverage = 14.22 ± 4.55). We calculated *r*^2^ between SNPs in 627 genes on different genome scaffolds of the Z chromosome ≤ 10 Mb apart for a total of 41,193 biallelic SNPs with MAF ≥ 0.25 (since 18/43 individuals had long PDD, only SNPs with a high minor allele frequency represent potential candidate mutations underlying PDD time). LD was high for genes < 2 Mb apart (maximum LD = 0.97, 99.9% quantile = 0.77), but decayed over larger physical distances (2-10 Mb: maximum LD = 0.77, 99.9% quantile = 0.56).

We identified LD outliers by calculating a 95% confidence interval for the 99.9% quantile of all gene pairs within a 1 Mb window (*N* = 10,000 bootstrap replicates). Among gene pairs located within or between the two QTL regions (≥ 2 Mb apart and ≤ 7 Mb), there were 12 outliers (Figure S10; Table S7). Of these, the most extreme LD outlier was between *per* and *Pdfr* (Figure 4; Figure S11) and specifically, maximum LD occurred between 9 SNPs in the 5’UTR intron of *Pdfr* and 3 SNPs in the 5’UTR intron of *per* (*r*^*2*^ = 0.75). In both genes, introns within the 5’ UTR contain E-box *cis*-regulatory enhancer elements where the circadian transcription factors CLOCK (CLK) and CYCLE (CYC) bind (46, 47). In *Drosophila melanogaster, Pdfr* contains one CLK-CYC binding site in the 5’UTR intron and *per* contains three in the 5’UTR intron and one E-box upstream of the promoter (46-47). The long-PDD corn borer allele for one of the high LD SNPs created a novel E-box element (CACGTG) in the 5’UTR and this allele was completely absent in the short-PDD populations (Table 1b). Additionally, *per* and *Pdfr* were present in other outlier gene pairs (*Pdfr* with the genes *magu* and *CG6752*; *per* with genes flanking *Pdfr*: *trol, meigo*, and *CG32809*), and one pair contained the circadian gene *clk* with 1-Cys peroxiredoxin (*Prx6005*). We also performed LD analysis on the outlier SNPs identified by BayPASS (*eBP*_*is*_ > 2) with all other SNPs (>1 Mb apart; MAF ≥ 0.25) and found the 2 outlier SNPs in *Pdfr* had the highest LD with SNPs in *per* (*r*^*2*^ = 0.73) and the single SNP in *per* had the highest LD with SNPs in *Pdfr* (*r*^*2*^ = 0.61). The SNP in *trol* had the highest LD with autophagy-related 9 (*Atg9*; *r*^*2*^ = 0.53).

**Figure 4.**
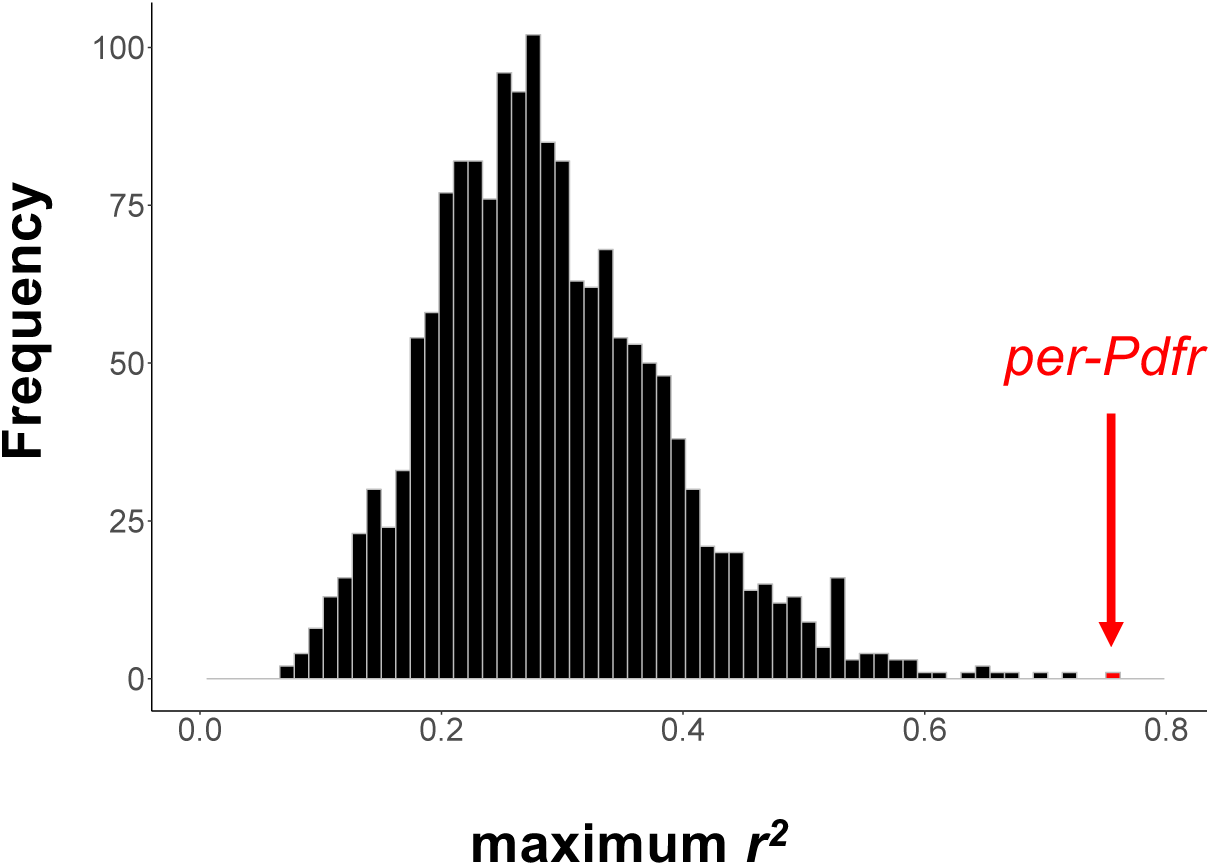
Linkage disequilibrium among genes in QTL1 and QTL2. Histogram of maximum linkage disequilibrium (LD, measured by *r2*). Red bar indicates *per*-*Pdfr*, the genes with the maximum LD observed for any genes in the two intervals over 1 Mb apart (N = 1,678 pairs).

Our genome-wide Bayesian and linkage disequilibrium outlier analyses suggest that *per* and *Pdfr* represent the best gene candidates in QTL1 and QTL2, respectively. We analyzed variation in the resequenced individuals using a case/control association analysis in plink (48) to detect other mutations within *per* and *Pdfr* that might have a phenotypic effect, such as changes in amino acids, changes in splice junctions leading to splice variants, and large structural variants that disrupt exons or *cis*-regulatory regions (other than E-boxes) (Table S9-S10). Using homology with *per* in other insects, we identified protein domains (27 exons) in the corn borer ortholog. Three nonsynonymous SNPs were significantly associated with PDD time and all were located in *per* exon 23 (Table 1c; outlined in red in Figure 5a). Both proline/threonine (P/T) and serine/proline (S/P) polymorphisms are in a 33 amino-acid region of *per* that is deleted in an artificially selected line of flesh fly (*Sacrophaga bullata*) showing enhanced diapause and the S/P polymorphism is 3 residues away from a 9 amino-acid insertion in a selected flesh fly line with decreased diapause (Figure 5a) (49). No consistent associations were found among splice variants, PDD time, and polymorphisms at splice sites (see Supplemental Infromation), nor were any large structural variants detected in within the gene.

**Figure 5.**
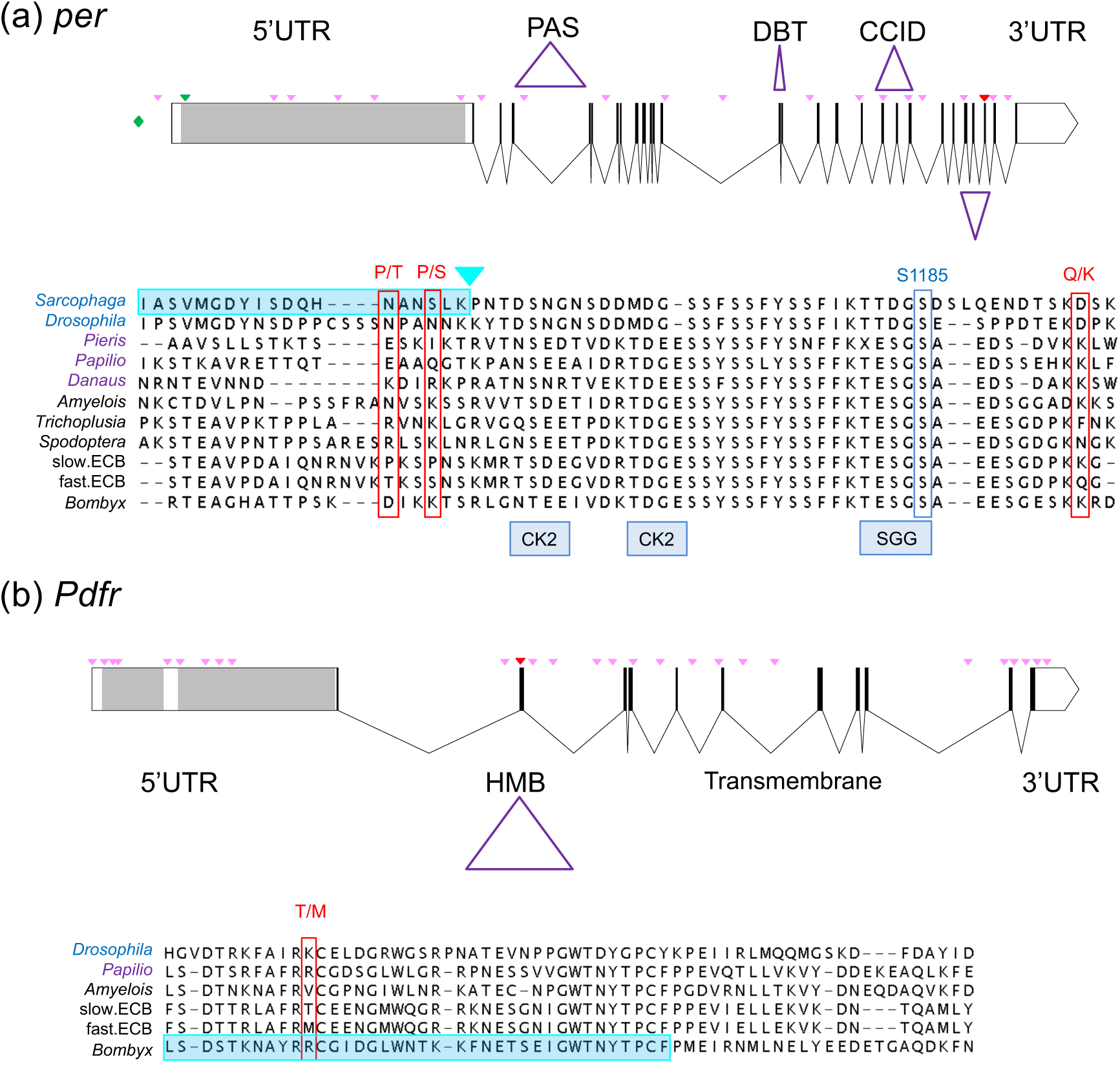
Gene models for candidate PDD time genes and amino acid changes. Candidate gene models including 5’ UTR, 3’ UTR, exons (black bars), with protein domains (purple triangles) labeled. Gray portions of 5’UTR are putative 5’UTR introns (i.e., sequences not present in RNA transcripts). Locations of polymorphisms that showed significant association (q < 0.01) in individual sequencing data indicated by light pink triangles, those that change amino acid sequence denoted by red triangles. Below, amino acid sequence for exons with differences in sequence between ECB short-and long-PDD populations aligned with selected species of flies (blue), butterflies (purple) and moths (black). a) Gene model for *per* in ECB, including the upstream E-box enhancer element (green diamond) and novel E-box in 5’UTR (green triangle). Domains for TIMELESS binding (PAS), DOUBLETIME binding (DBT) and CLOCK-CYCLE inhibitory domain (CCID) indicated. Two amino acid changes (outlined in red) are in the same region which *Sarcophaga* high diapausing mutants have a deletion (shaded in teal) and non-diapausing mutants have an insertion (teal triangle; (49)). The amino acid changes also flank a region containing several predicted casein kinase 2 (CK2) sites in ECB and a conserved serine phosphorylated by SHAGGY identified in *Drosophila* (outlined in blue; (125)); N=78 polymorphisms. b) Gene model for *Pdfr* including PDF hormone binding domain (HMB), and transmembrane domain (7 alpha helices). Amino acid sequence shown for portion of the PDF hormone binding domain region annotated in *Bombyx mori* (outlined in teal) with differences between ECB slow and fast populations outlined in red; N = 166 polymorphisms.

*Pdfr* consisted of 12 exons (Figure 5b). In the predicted extracellular hormone binding domain for PDF (exon 2) there was one nonsynonymous SNP, coding for a methionine in short PDD individuals and a threonine in long PDD individuals (*q* = 1.12 ×10^−8^; Figure 4b; Table 1c). There were no splice junction polymorphisms or variants in *Pdfr*. Although its specific sequence is unknown, an enhancer is located ∼8.5 kb upstream of *Pdfr* in *D. melanogaster* (42). In corn borers, we found a ∼419 kb inversion associated with PDD time (*q* = 6.48 × 10^−9^) with one breakpoint 7.05 kb upstream from *Pdfr* in this putative enhancer region. The second breakpoint was predicted to occur 162 kb after *trol*.

### Circadian activity

Prior work has shown that circadian rhythm of locomotor activity in mice, fruit flies, and humans is the behavioral output of circadian clock genes (50) and that mutations in *per* and *Pdfr* result in altered circadian rhythms in *D. melanogaster* (43, 51). We evaluated evidence for a difference in free-running circadian rhythm under total darkness (DD) between adult moths with short and long PDD times. Male pupae entrained to 16:8 were transferred to DD shortly before eclosion. We found that endogenous period length (τ) differed by approximately 1.3 hours, with short-PDD males showing longer average circadian periods (τ = 22.7 ± 0.2 h, *N* = 24) than long-PDD males (τ = 21.4 ± 0.28 h, *N* = 22) (ANOVA, *F*_1,44_ = 13.79, *P* < 0.001; Figure 6; Supplemental Results).

**Figure 6.**
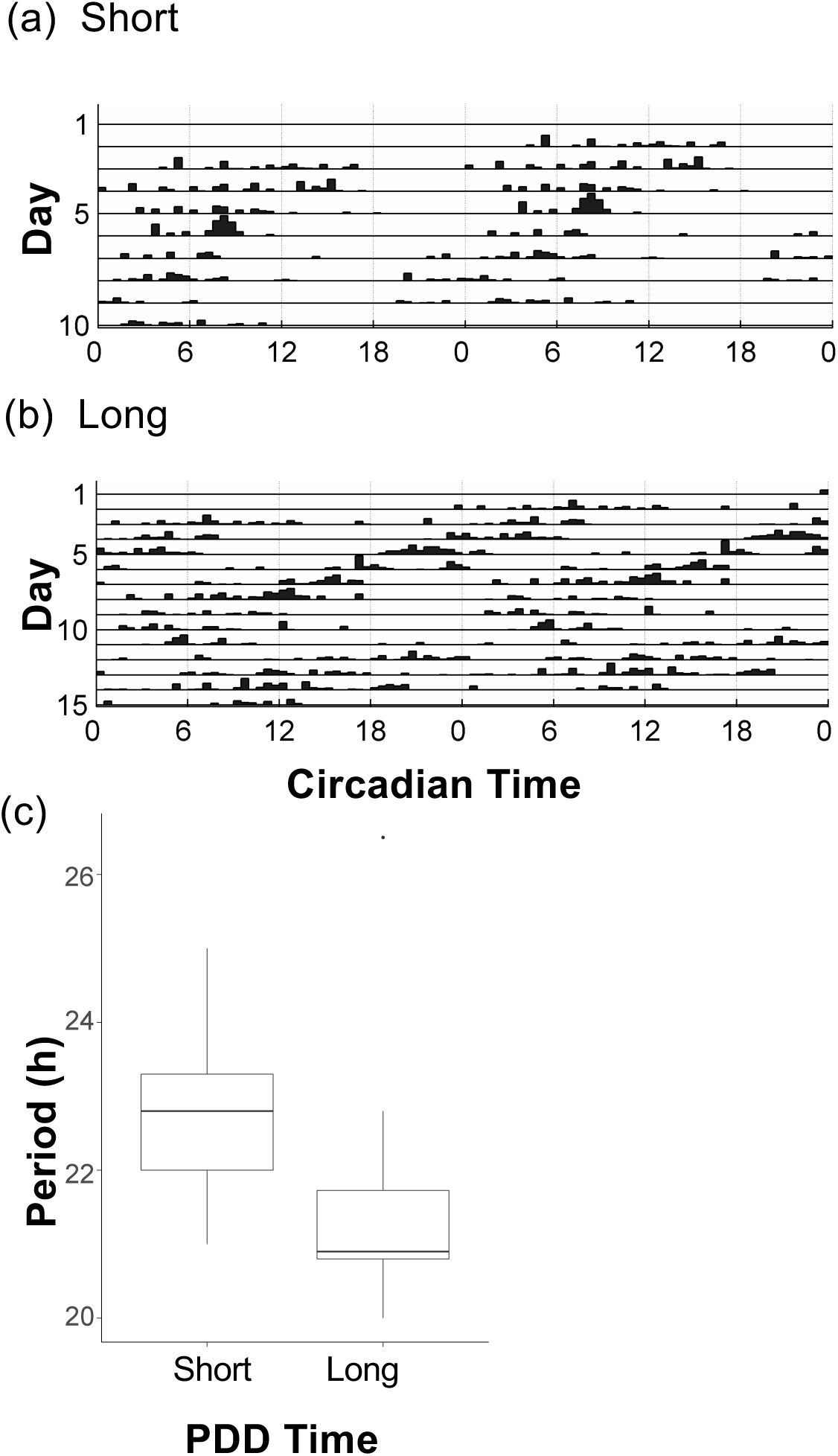
Circadian period for short- and long-PDD insects. Actograms showing the locomotor activity in complete darkness (DD) over 2 day windows (double-plotted) for up to 15 days used to estimate circadian period (τ) for a representative a) short-PDD male, b) long-PDD male. c) Boxplot of length of circadian period (in hours) for males with short and long PDD (median, first and third quartile shown), lines indicate 95% confidence interval. Long-PDD individuals have a significantly shorter period (*P* < 0.001); N = 46 adults.

## DISCUSSION

Genetic changes in circadian clock genes associate with natural variation in the time needed to end winter diapause and return to active springtime development in the European corn borer. Clock genes *per* and *Pdfr* are located within two epistatic QTL, strongly differ in allele frequency among individuals that pupate earlier or later, have the highest linkage disequilibrium among gene pairs in the QTL regions, and possess amino-acid changes that may affect protein function. *Per* alleles containing an additional putative CLK-CYC binding site were also exclusively identified in populations that pupate later. While additional work is needed to understand how identified allelic variants affect gene function and to verify that there are no other genetic polymorphisms contributing to diversity in seasonal timing, our combined results suggest that allelic variation in *per* and *Pdfr* is causal to evolution of diapause timing when confronting rapid environmental changes associated with range expansion (Figure S2) (22) and human-induced climate warming (23).

The presence of epistatic QTL indicates that genes underlying PDD time are likely members of the same genetic pathway. Both *per* and *Pdfr* interact in circadian pacemaker neurons in insect brains, where they synchronize biological activity to daily cycles of night and day (Figure 7) (52-54). In the laboratory, mutations in *per* are known to alter the length of the circadian activity period (51) and null mutants lose rhythm completely in *D. melanogaster* (41). Likewise, *Pdfr* is integral to the function of circadian pacemaker neurons in insect brains, where they receive secreted PDF neuropeptides that coordinate, synchronize, and reset the clock neuron network to new light cycles (55-57). Loss of expression of *Pdf* or *Pdfr* in these neurons can result in shorter circadian activity period (τ), abnormal peaks of circadian activity, and an inability to entrain to longer photoperiods in *D. melanogaster* (55, 58-60). Although robust connections between circadian clock genes and seasonal phenotypes have been discovered in plants (61), evidence in insects has been primarily based on RNAi studies demonstrating that functional clock genes are essential for diapause (62-64). It is less clear whether allelic variation in these genes typically responds to selection in natural populations to drive changes in the seasonal timing of diapause transitions. For example, polymorphism in the timeless gene in *D. melanogaster* influences diapause capacity in the laboratory, but in nature, latitudinal variation of timeless does not match variation in diapause and observed patterns are opposite of those predicted (65). Instead, diapause differences are more strongly associated with non-circadian genes, such as couch potato (66). Similarly, in *Wyeomyia smithii* pitcher plant mosquitos diapause appears to be independent of the circadian pathway, with these traits evolving separately in selection lines (67). Our study provides the first evidence that *per* and *Pdfr*, core components of the molecular clock, are associated with the duration of developmental arrest for insects in the spring. Recent studies in two other insects show that *per* alleles are associated with polymorphism in the timing of autumnal initiation of diapause (critical photoperiod for entrance) (68, 69). Genetic changes at *per* may therefore provide the capacity to adjust diapause transition times across two different seasons, enabling insects to synchronize with both the end and beginning of winter.

**Figure 7.**
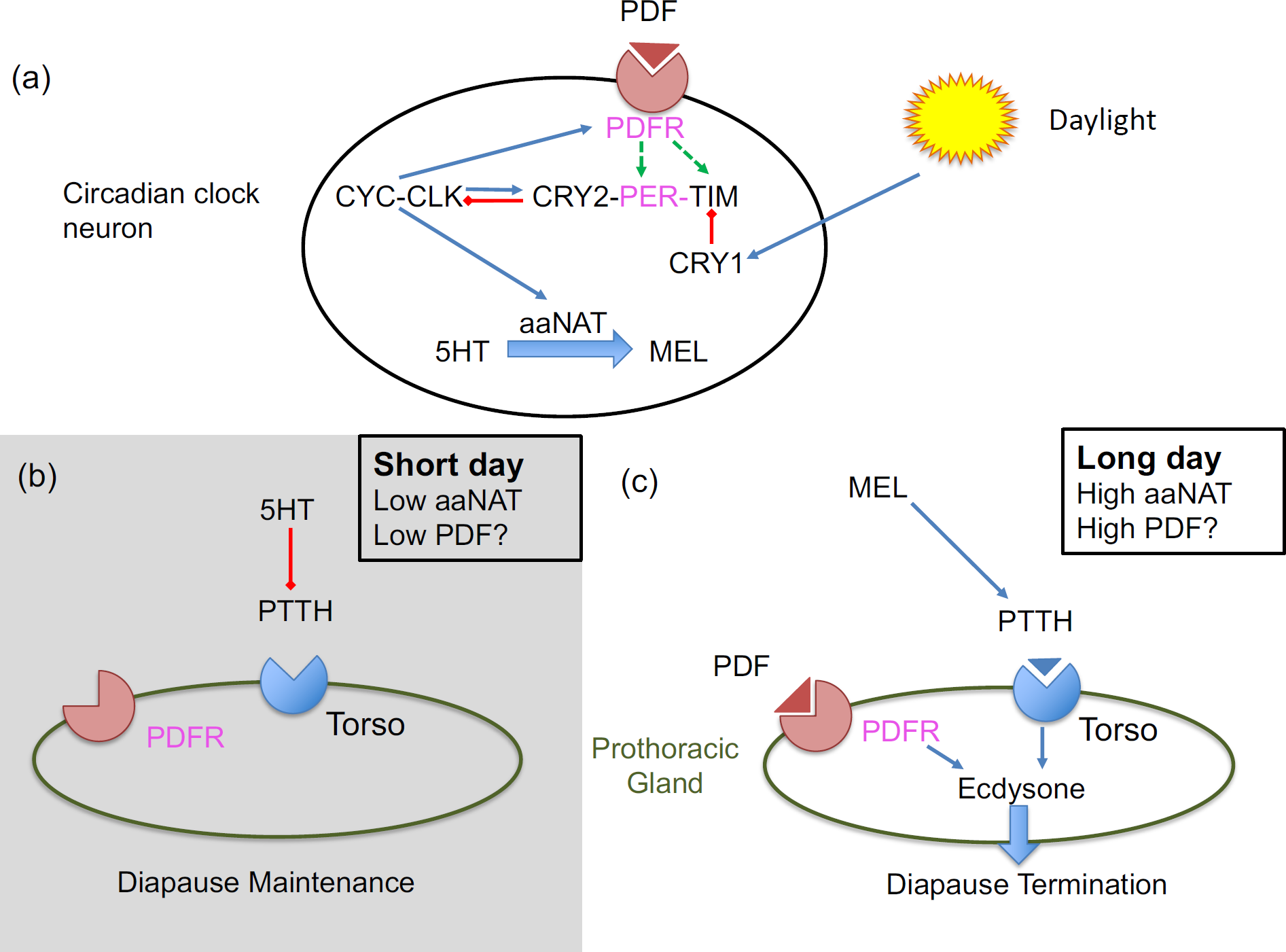
Hypothesized pathway for circadian clock involvement in diapause termination. Regulation of circadian clock genes in clock pacemaker neurons shown (adapted from Lepidopteran clock in (125)). Blue arrows indicate activation, red arrows are suppression, black dashes are heterodimer formation, green dashed arrows are stabilization. Candidate genes shown in pink. The heterodimer formed by Clock (CLK) and Cycle (CYC) upregulate Period (PER), Timeless (TIM), and Pigment dispersing factor receptor (PDFR) (47). When PER and TIM are bound to Cryptochrome2 (CRY2), they migrate into the nucleus and PER-CRY2 repress CLK-CYC (126). Cryptochrome1 (CRY1) degrades TIM in the presence of light. The neurotransmitter pigment dispersing factor (PDF) binds to its receptor (PDFR) and this activation stabilizes both TIM (82) and PER (52). CLK-CYC activates arylalkylamine N-acetyltransferase (aaNAT) which converts Serotonin (5HT) to Melatonin (MEL) (81). b) Under short day conditions, serotonin levels are high, preventing PTTH release and leading to diapause maintenance. c) Under long days, melatonin levels are high and PTTH is released, leading to activation of ecdysone release by the PG and diapause termination. Ecdysone release is also facilitated by activation of PDFR in the PG (84).

In 1936, Bünning hypothesized that mechanisms underlying circadian rhythmicity control circannual rhythmicity (70). Alternatively, the circadian clock and the seasonal timer could act as two modules with largely separate genes, although individual genes may have cross-module effects (71-73).We find that populations of European corn borer moth differing in PDD time also differ in their internal circadian oscillator, such that the population spending more time in diapause (long PDD time) shows an accelerated circadian period (shorter τ). A similar inverse relationship between circadian and circannual rhythm has been found in Scandinavian flies (*Drosophila littoralis*), where shorter circadian periods are associated with earlier diapause initiation (74), and in mustard plants (*Boechera stricta*), where shorter circadian periods are associated with delayed flowering (75). Combined with the fact that multiple interacting circadian clock genes (*per, Pdfr*) are implicated in photoperiodic diapause termination, patterns of circadian activity in the European corn borer moth suggest that allelic variation and interactions between *per* and *Pdfr* might affect seasonal timing by altering circadian clock function (modular pleiotropy), rather than by a direct effect on diapause that is unrelated to the circadian clock (gene pleiotropy). Further evidence for at least partial circadian control of diapause termination timing was found by Beck (76), who tested Bünning’s model in the European corn borer moth using the Nanda-Hamner protocol. He found a circadian resonance cycle (∼24 h peaks) between the period of the diapause inducing photoperiod and PDD time, supporting the hypothesis that diapause timing is mediated or controlled by a circadian based physiological system. Future work will be needed to understand how molecular mechanisms might directly link expression of the daily clock and the seasonal timer in this species.

Physiological experiments suggest several molecular mechanisms by which *per* and *Pdfr* could regulate the neuroendocrine switch underlying the transition from diapause to development. Larval termination of diapause in the European corn borer moth and many other Lepidoptera is triggered by release of the developmental hormone ecdysone from the prothoracic gland (PG) due to stimulation from prothoracicotropic hormone (PTTH) (77-79). Work in the Chinese oak silkmoth (*Antheraea pernyi*) suggests that PTTH release or synthesis is regulated by the circadian clock pathway via the indolamine metabolism pathway. Specifically, a key step may involve the enzyme arylalkylamine N-acetyltransferase (*aaNAT*) and its opposing interaction on levels of melatonin (MEL) and gated PTTH synthesis/release under long-day photoperiod, or on levels of serotonin and PTTH suppression under short-day photoperiod (Figure 7) (80, 81). In *A. pernyi, aaNAT* is synthesized in circadian clock neurons when levels of CLK-CYC are high. PER represses CLK-CYC activity and RNAi against *per* results in increased *aaNAT* transcription, increased MEL protein, and diapause termination (81). In *D. melanogaster*, CLK-CYC binds to *Pdfr*, putatively regulating its expression (47). Activation of PDFR by PDF binding increases protein kinase A (PKA), which stabilizes PER and TIMELESS (TIM), preventing degradation, and increasing circadian period by ∼2 h (52, 82). Thus, *per* and *Pdfr* alleles of the European corn borer moth may function differently under seasonal changes in photoperiod by interacting in pacemaker neurons to alter *aaNAT* production, influencing synthesis/release of PTTH and the timing of diapause termination. Some evidence of differential regulatory control of the circadian clock–indolamine pathways exist between short and long PDD populations, potentially due to changes at *per* and *Pdfr*. We previously found that transcription in adult female heads of *aaNAT*, its putative regulator (*cyc*), and its downstream target (*PTTH*) is lower in strains with longer than shorter PDD times one hour before the light-dark transition under long-day photoperiod (83). If the novel E-box element we identified in *per* from long-PDD individuals leads to increased *per* expression and repression of *cyc*, it could hypothetically lower *aaNAT*, leading to a perception of days as shorter and delaying diapause termination. In addition to interaction of *per* and *Pdfr* in pacemaker neurons, a second route for epistasis and control of termination could occur by the release of ecdysone through an independent *Pdfr* cascade in the PG discovered in the silkmoth *Bombyx mori* (Figure 7) (84). Indeed, knockdowns of *Pdf* are sufficient to induce diapause under long photoperiods in mosquitos (*Culex pipiens*) and ablation of PDF-positive neurons impairs the photoperiodic regulation of diapause in bean bugs (*Riptortus pedestris*) and blow flies (*Protophormia terraenovae*) (63,85,86).

Despite repeated observation of geographic variation in circannual rhythm within and among species, and widespread alterations of seasonal activity in response to climate change and range expansion (3, 15, 20, 30, 87), seasonal timing in nature has rarely been linked to causal mechanisms. This gap in knowledge is alarming given that recent work suggests that roughly half of Lepidopteran species may be currently in decline (88) and accumulating connections between seasonal timing flexibility and population persistence (89). Establishing the genomic determinants of circannual variation is essential for understanding the capacity of species to tolerate rapidly changing environments (encountered through species movement or changes in local climate), as well as to accurately predict their future evolutionary trajectories (across geographic space and through time) (10, 13). We have shown using multiple whole-genome approaches in the European corn borer that evolution to earlier spring termination of diapause and an associated added generation has a relatively simple genetic basis, likely involving two genes that also orchestrate circadian timekeeping. Earlier springtime activity can allow populations to track preferred seasonal environments and to produce more generations per year, both of which improves population tolerance of sustained environmental change in theoretical (10) and empirical studies (87, 89). The duration of insect diapause generally tracks winter length, which will decrease by a month or more over the next century according to most climate change models (90, 91). Therefore, intense selection on alleles at *per* and *Pdfr* in this species is likely to be an important component of continued adaptation, anticipated range expansion (92, 93), and long-term species persistence under rapidly changing seasonal environments. As a major pest of corn and other crops in North America and Europe, the ecological and economic ramifications of these microevolutionary changes will be significant. To understand why certain pests like *Ostrinia* moths have the capacity to become greater threats under projected climates and why certain beneficial species may require enhanced conservation management to prevent extinction, future efforts should be made to more broadly understand the mechanisms underlying circannual rhythm in nature.

## METHODS

### QTL mapping of termination time

Backcross F_2_ female offspring, F_1_ parents and F_0_ grandparents were genotyped for polymorphic SNPs segregating within families using multiplexed PCR amplicons (500 bp amplicons, 384 unique individual barcodes per lane) sequenced on an Illumina MiSeq at the Cornell University Sequencing Facility (primer sequences used are described in Kozak et al. (35)). Additional Z-linked and autosomal markers were genotyped using Sequenom Assays developed for polymorphic SNPs (Sequenom Assay Design Suite 1.0, Sequenom, San Diego, CA, USA) and run at the Iowa State University Center for Plant Genomics (ISU-CPG) as described in Coates et al. (94) and Levy et al. (26). Linkage maps were constructed for each family separately using a maximum recombination frequency of 0.35 and a minimum LOD of 3 in R 3.4 using rQTL and the estimate map function (95-97).

QTL mapping of PDD time was performed using the scanone and scantwo functions with standard interval mapping and an interval of 0.5 cM (similar results were obtained when using extended Haley-Knott regression) for autosomal and Z linked markers in 1 family (F6, 67 offspring) and the Z chromosome only using 5 families (F2,5,6,9,11; 226 offspring total). For 5 families, a consensus Z map was constructed from the individual Z family maps using the LPmerge package in R (root mean squared = 10.89 and standard deviation = 7.31) (98). The 95% Bayesian credible interval (BCI) for the QTL were estimated and significance of QTL determined by F-test comparisons of models with and without QTL and their interaction using fitqtl function. For estimating BCI for the epistatic QTL, mapping was repeated on 101 individuals with the slow genotype at QTL1.

### Population genomic analyses

We sampled from 5 European corn borer (ECB) populations (see Table S3). Individuals were collected from the field as diapausing larvae and PDD time was characterized in the lab. ECB have two known pheromone strains (E and Z) and field caught individuals were classified as Z strain as determined by genotyping at a polymorphic *Taq*1α restriction endonuclease cleavage site in the gene responsible for differences in pheromone components, *pgfar* (99). PDD time was measured as the number of days for diapausing larva to pupate after being placed in 16:8 LD and 26°C. Fast PDD individuals pupate < 19 days after exposure to these conditions while slow PDD individuals pupate after ≥ 39 days (24, 25, 29). For some long PDD individuals, we only had time to eclosion data (when adults emerged from puparium). Mean time ± SD from pupation to eclosion for BV = 9.9 ± 2.9 (N=107) so we conservatively estimated PDD time by subtracting 13 days from time to eclosion. Pennsylvania individuals were collected as adults in pheromone traps (100) and this population has been consistently phenotyped as fast PDD/bivoltine over a 15-year period (101, 102).

For each population, samples were pooled using equal DNA quantities from each individual. DNA was extracted using the Qiagen DNeasy tissue protocol except tissues were not vortexed during isolation to preserve high molecular weight DNA. Pooled libraries were prepared using the Illumina TruSeq protocol (Illumina Inc., San Diego, CA). Libraries were sequenced on an Illumina HiSeq3000 at the Iowa State University DNA Facility using 150 bp paired-end sequencing and 2 libraries run per lane. Genomic reads were trimmed using Trimmomatic v.35 to remove Illumina adapters (TruSeq2 single-end or TruSeq3 paired-end), reads with a phred quality score (*q*) < 15 over a sliding window of 4 and reads < 36 bp long. Trimmed genomic data were aligned to the 454.7 Mb draft ECB genome (GenBank BioProject PRJNA534504; BioSample SAMN11491597; accession SWFO00000000) which consists of 8,843 scaffolds (N50 = 392.5 kb, largest scaffold = 3.32 Mb; BUSCO 3.0.2 (103) score = 93.1% complete from 1066 from the arthropoda_odb9 gene set (Table S2). Prior to alignment repetitive regions were masked by RepeatMasker (using Drosophila melanogaster TE library from repbase; http://www.repeatmasker.org/; accessed March 2017; (104)). Genomic scaffold chromosomal location was determined as described in the supplemental material. Alignment was done using Bowtie2 (105). Due to poor quality of some of the reverse mate libraries, the number of reads aligned was found to be higher when reads were aligned as single-end libraries (forward mate pairs, broken mate pairs). Filtration of low quality alignments and duplicates were performed using picard tools (http://picard.sourceforge.net).

Samtools was used to identify SNPs (106). Scripts from the Popoolation2 package (107,108) were used to filter SNPs (removing SNPs near small indels, and those with rare minor alleles that did not appear twice in each population), calculate allele (read count) frequency of SNPs using a minimum coverage of 14 reads and a maximum coverage of 200, calculate *F*_*ST*_ over non-overlapping 1 kb windows (with >100 bp above minimum coverage in all populations), and perform CMH tests (see Supplemental Material). We used population read counts from Popoolation to test the association of alleles among our 5 populations while controlling for population demography using BayPASS 2.1 (39) and the standard (STD) model with PDD time (slow = 1, fast = −1) and d_0_y_ij_ = 6. Significantly associated alleles were defined as SNPs that had *XtX* above the 0.001% quantile of pseudo-observed data (POD) of simulated “neutral” loci (using simulate.baypass and mean read coverage for each population), BF > 20 *dB* (the difference between a model with and without PDD time included; with BF = 20 indicating “decisive” evidence in support of an association; (109)) and *eBPis* > 2 (which estimates how likely it is that the posterior distribution of β includes zero; equivalent to P < 0.01) (39, 40, 110). BayPASS was run separately for Z chromosome (14,724 SNPs; haploid pool sizes: EA = 50, GEN = 38, LA = 78, PY = 37, BV = 34) and autosomal loci (N = 577,412 SNPs; EA = 68, GEN = 50, LA = 78, PY = 52, BV = 40).

### Individual resequencing

To identify the specific polymorphisms associated with voltinism differences and calculate LD, individual re-sequencing was done for 18 slow PDD individuals (10 GEN, 4 BV, 4 PY), 25 fast PDD individuals (14 EA, 11 LA). Individual libraries were prepared using the Illumina TruSeq protocol and were sequenced on an Illumina NextSeq using 150 bp paired-end sequencing at Cornell University. Trimmed genomic data were analyzed using the GATK best practices pipeline (111-113). Data were aligned to the draft reference ECB genome using BWA (106). Aligned reads were sorted and filtered using picard and samtools to remove duplicates and reads with a mapping quality score (Q) below 20. SNPs and small indels (< 50 bp indels) were called using GATK Haplotype caller (run in joint genotyping mode) after realigning around indels. Variants were filtered using recommended GATK hard filters (113). Larger structural variants (indels > 300 bp and inversions) were called from individual aligned bam files using information from split paired end reads in Delly2 (114).

### Linkage disequilibrium

LD was calculated after the phase of genotypes was imputed using Beagle 5.0 (115). Prior to LD calculation, phased genotypes were filtered to include only those located within genes and MAF ≥ 0.25 and inter-scaffold *r*^2^ was calculated in vcftools (116). We summarized *r*^2^ over genes and performed bootstrapping analyses in R using data.table, plyr, and boot packages (117-119). Plots were constructed using the ggplot2 and qqman packages (120-121).

We ran an association analysis on sequencing data from individual ECB samples. GATK allele calls for SNPs and small indels (< 50 bp) were combined with delly2 variant calls of large indels and inversions using the combine variants function in GATK. We then analyzed the association of these polymorphisms with PDD time in plink 1.9 (48) with PDD time coded as a binary case/control phenotype (1 = fast, 2 = slow) and using a Fisher’s exact test to detect significant differences in allele frequencies. P-values were FDR corrected genome-wide using the fdrtools package in R (122).

### Circadian activity

To measure the endogenous circadian clock, we used laboratory colonies from BV (slow-PDD) and a colony from a fast-PDD population collected near Geneva NY raised in the lab at 16:8 and 26°C (25, 28, 34, 35, 38). After pupation, male pupae were transferred to tubes within activity monitors in free running conditions (total darkness; DD) at 26°C. Activity was measured using a Trikinetics activity monitor (model LAM25, Waltham, MA) from the first day of adult eclosion. 16 individuals of each type were measured in two replicates for a total of 32 individuals per PDD type. Data were analyzed using custom MATLAB toolboxes (123).

## ACKNOWLEDGEMENTS

Gabriel Golczer and Erastus Thuo assisted with DNA isolations. Henry Kunerth and Ben Hamilton assisted with sample acquisition. We thank F. Rob Jackson, Mary Roberts, and Jasper and Rigel Hatch Dopman for help conducting circadian activity experiments. This research was funded by the National Science Foundation (DEB-1257251 to E.B.D.; DEB-1256688 to R.G.H), Tufts University, a Tufts University Faculty Research Award (E.B.D.), and cooperative agreement 58-5030-7-066 between the United States Department of Agriculture, Agricultural Research Service (USDA-ARS), and Tufts University. Funding was also received from USDA-ARS Project CRIS-5030-22000-018-00D, USDA-ARS: Project CRIS-3625-22000-017-00 and the Iowa Agriculture and Home Economics Experiment Station, Ames, IA Project 3543. This article reports the results of research only and any mention of products or services does not constitute an endorsement by USDA-ARS. USDA-ARS is an equal opportunity employer and provider. While performing this research, S.C.K. was partially funded by the Tufts University Summer Scholars program and C.B.W. by a National Science Foundation Graduate Research Fellowship (2011-116050).

## Author contributions

G.M.K. and E.B.D. designed and performed research, analyzed data, and wrote the paper. B.S.C. contributed population genomic and genome sequencing. C.B.W. and S.C.K. performed primer design, DNA isolation, PDD phenotyping. S.M.B. performed primer design, amplicon and individual resequencing library preparation. R.G.H. assisted with research design.

## Data deposition

### Genbank (sequencing data)

ECB Genome: BioProject PRJNA534504; BioSample SAMN11491597; accession SWFO00000000 Pool-seq: BioProject PRJNA540655 (BV,Gen,EA,PY); BioProject PRJNA361472 (LA: SRX249882) Indiv-seq: BioProject PRJNA540833

## REFERENCES

1. Danks HV Insect dormancy: an ecological perspective (Biological Survey of Canada, Ottawa). (1987).

2. Chuine I (2010) Why does phenology drive species distribution? Philos Trans Roy Soc B 365:3149–3160.

3. Walther GR, et al. (2002) Ecological responses to recent climate change. Nature 416:389–395.

4. Bradshaw WE, Holzapfel CM (2008) Genetic response to rapid climate change: it’s seasonal timing that matters. Mol Ecol 17:157–166.

5. Willis CG, Ruhfel B, Primack RB, Miller-Rushing AJ, Davis CC (2008) Phylogenetic patterns of species loss in Thoreau’s woods are driven by climate change. Proc Natl Acad Sci USA 105:17029–17033.

6. Møller AP, Rubolini D, Lehikoinen E (2008) Populations of migratory bird species that did not show a phenological responseto climate change are declining. Proc Natl Acad Sci USA 105: 16195–16200.

7. Roff DA (1980) Optimizing developmental time in a seasonal environment: The “ups and downs” of clinal variation. Oecologia 45:202–208.

8. Roff DA (1983) Phenological adaptation in a seasonal environment: a theoretical perspective. Diapause and Life Cycle Strategies in Insects (Junk, The Hague), pp 253–270.

9. Stearns SC (1992) The evolution of life histories (Oxford University Press).

10. Chevin L-M, Lande R, Mace GM (2010) Adaptation, Plasticity, and Extinction in a Changing Environment: Towards a Predictive Theory. PLoS Biol 8:e1000357.

11. Helm B, et al. (2013) Annual rhythms that underlie phenology: biological time-keeping meets environmental change. Proc R Soc Lond B Biol Sci 280:20130016.

12. Van Dyck H, Bonte D, Puls R, Gotthard K, Maes D (2014) The lost generation hypothesis: could climate change drive ectotherms into a developmental trap? Oikos 124(1):54–61.

13. Hoffmann AA, Sgrò CM (2012) Climate change and evolutionary adaptation. Nature 470:479–485.

14. Visser ME, Caro SP, Van Oers K, Schaper SV, Helm B (2010) Phenology, seasonal timing and circannual rhythms: towards a unified framework. Philos Trans Roy Soc B 365:3113–3127.

15. Denlinger DL, Hahn DA, Merlin C, Holzapfel CM, Bradshaw WE (2017) Keeping time without a spine: what can the insect clock teach us about seasonal adaptation? Philos Trans Roy Soc B 372:20160257–9.

16. Saunders DS (1980) Some effects of constant temperature and photoperiod on the diapause response of the flesh fly, *Sarcophaga argyrostoma*. Physiol Entomol 5:191–198.

17. Tauber MJ, Tauber CA, Masaki S (1986) Seasonal Adaptations of Insects (Oxford University Press).

18. Koštál V (2006) Eco-physiological phases of insect diapause. J Insect Physiol 52:113–127.

19. Bradshaw WE, Holzapfel CM (2010) Light, Time, and the Physiology of Biotic Response to Rapid Climate Change in Animals. Annu Rev Physiol 72:147–166.

20. Danilevsky AS, Goryshin NI, Tyshchenko VP (1970) Biological rhythms in terrestrial arthropods. Annu Rev Entomol 15:201–244.

21. Caffrey DJ, Worthley LH (1927) Progress report on the investigations of the European corn borer. Series: Department bulletin (United States. Dept. of Agriculture); no. 1476.

22. Showers WB (1979) Effect of diapause on the migration of the European corn borer into the southeastern United States. In: Movement of highly mobile insects: concepts and methodology in research; proceedings of a conference (Raleigh, NC, University Graphics, 1979).

23. Derron JO, Goy G, Breitenmoser S (2009) Biological characterisation of the bivoltine race of the European corn borer (*Ostrinia nubilalis*) in the Lake Geneva region. Revue Suisse d’Agriculture 41:179–184.

24. Glover TJ, Knodel JJ, Robbins PS, Eckenrode CJ, Roelofs WL (1991) Gene Flow Among Three Races of European Corn Borers (Lepidoptera: Pyralidae) in New York State. Environ Entomol 20:1356–62.

25. Dopman EB, Perez L, Bogdanowicz SM, Harrison RG (2005) Consequences of reproductive barriers for genealogical discordance in the European corn borer. Proc Natl Acad Sci USA 102:14706–14711.

26. Levy RC, Kozak GM, Wadsworth CB, Coates BS, Dopman EB (2015) Explaining the sawtooth: latitudinal periodicity in a circadian gene correlates with shifts in generation number. J Evol Biol 28:40–53.

27. Istock CA (1981) Natural selection and life history variation: theory plus lessons from a mosquito. Insect Life History Patterns (Springer), pp 113–127.

28. Wadsworth CB, Woods WA Jr, Hahn DA, Dopman EB (2013) One phase of the dormancy developmental pathway is critical for the evolution of insect seasonality. J Evol Biol 26:2359–2368.

29. Dopman EB, Robbins PS, Seaman A (2010) Components of reproductive isolation between North American pheromone strains of the European corn borer. Evolution 64:881–902.

30. Altermatt F (2010) Climatic warming increases voltinism in European butterflies and moths. Proc R Soc Lond B Biol Sci 277:1281–1287.

31. McLeod DGR (1978) Genetics of diapause induction and termination in the European corn borer, *Ostrinia nubilalis* (Lepidoptera: Pyralidae), in Southwestern Ontario. Can Entomol 110:1351–1353.

32. Glover TJ, Robbins PS, Eckenrode CJ, Roelofs WL (1992) Genetic control of voltinism characteristics in European corn borer races assessed with a marker gene. Arch Insect Biochem Physiol 20:107–117.

33. Wadsworth CB, Dopman EB (2015) Transcriptome profiling reveals mechanisms for the evolution of insect seasonality. J Exp Biol 218:3611–3622.

34. Wadsworth CB, Li X, Dopman EB (2015) A recombination suppressor contributes to ecological speciation in *Ostrinia* moths. Heredity:1–8.

35. Kozak GM, et al. (2017) A combination of sexual and ecological divergence contributes to rearrangement spread during initial stages of speciation. Mol Ecol 26:2331–2347.

36. McLeod DGR, Beck SD (1963) Photoperiodic termination of diapause in an insect. Biol Bull 124:84–96.

37. Beck SD, Alexander N (1964) Chemically and photoperiodically induced diapause development in the European corn borer, Ostrinia nubilalis. Biol Bull 126:175–184.

38. Dopman EB, Bogdanowicz SM, Harrison RG (2004) Genetic mapping of sexual isolation between E and Z pheromone strains of the European corn borer (*Ostrinia nubilalis*). Genetics 167:301–309.

39. Gautier M (2015) Genome-wide scan for adaptive divergence and association with population-specific covariates. Genetics 201(4):1555–1579.

40. Bourgeois YX, et al. (2017) A novel locus on chromosome 1 underlies the evolution of a melanic plumage polymorphism in a wild songbird. Royal Society open science 4:160805.

41. Reddy P, et al. (1984) Molecular analysis of the period locus in Drosophila melanogaster and identification of a transcript involved in biological rhythms. Cell 38:701–710.

42. Hyun S, et al. (2005) Drosophila GPCR Han is a receptor for the circadian clock neuropeptide PDF. Neuron 48:267–278.

43. Lear BC, et al. (2005) AG protein-coupled receptor, groom-of-PDF, is required for PDF neuron action in circadian behavior. Neuron 48:221–227.

44. Lindner JR, et al. (2007) The *Drosophila* Perlecan gene *trol* regulates multiple signaling pathways in different developmental contexts. BMC Dev Biol 7:121.

45. Wiberg RAW, Gaggiotti OE, Morrissey MB, Ritchie MG (2017) Identifying consistent allele frequency differences in studies of stratified populations. Methods Ecol Evol 8:1899–1909.

46. Taylor P, Hardin PE (2008) Rhythmic E-box binding by CLK-CYC controls daily cycles in *per* and t*im* transcription and chromatin modifications. Mol Cell Biol 28:4642–4652.

47. Meireles-Filho AC, Bardet AF, Yáñez-Cuna JO, Stampfel G, Stark A (2014) Cis-regulatory requirements for tissue-specific programs of the circadian clock. Curr Biol 24:1–10.

48. Chang CC, et al. (2015) Second-generation PLINK: rising to the challenge of larger and richer datasets. Gigascience 4:7.

49. Han B, Denlinger DL (2009) Length variation in a specific region of the period gene correlates with differences in pupal diapause incidence in the flesh fly, *Sarcophaga bullata*. J Insect Physiol 55:415–418.

50. Panda S, Hogenesch JB, Kay SA (2002) Circadian rhythms from flies to human. Nature 417:329–335.

51. Rutila JE, Edery I, Hall JC, Rosbash M (1992) The analysis of new short-period circadian rhythm mutants suggests features of *D. melanogaster* period gene function. J Neurogenet 8:101–113.

52. Li Y, Guo F, Shen J, Rosbash M (2014) PDF and cAMP enhance PER stability in Drosophila clock neurons. Proc Natl Acad Sci USA 111:E1284–E1290.

53. Michael TP, et al. (2003) Enhanced fitness conferred by naturally occurring variation in the circadian clock. Science 302:1049–1053.

54. Zheng X, Sehgal A (2008) Probing the relative importance of molecular oscillations in the circadian clock. Genetics 178:1147–1155.

55. Renn SC, Park JH, Rosbash M, Hall JC, Taghert PH (1999) A *pdf* neuropeptide gene mutation and ablation of PDF neurons each cause severe abnormalities of behavioral circadian rhythms in *Drosophila*. Cell 99:791–802.

56. Závodská R, et al. (2012) Is the sex communication of two pyralid moths, *Plodia interpunctella* and *Ephestia kuehniella*, under circadian clock regulation? J Biol Rhythms 27:206–216.

57. Xu G, et al. (2016) Identification and expression profiles of neuropeptides and their G protein-coupled receptors in the rice stem borer *Chilo suppressalis*. Sci Rep 6:28976.

58. Yoshii T, et al. (2009) The neuropeptide pigment-dispersing factor adjusts period and phase of *Drosophila*’s clock. J Neurosci 29:2597–2610.

59. Im SH, Li W, Taghert PH (2011) PDFR and CRY signaling converge in a subset of clock neurons to modulate the amplitude and phase of circadian behavior in *Drosophila*. PLoS ONE 6:e18974.

60. Schlichting M, et al. (2016) A neural network underlying circadian entrainment and photoperiodic adjustment of sleep and activity in *Drosophila*. J Neurosci 36:9084–9096.

61. Sawa M, Nusinow DA, Kay SA, Imaizumi T (2007) FKF1 and GIGANTEA complex formation is required for day-length measurement in *Arabidopsis*. Science 318:261–265.

62. Ikeno T, Numata H, Goto SG (2011) Circadian clock genes period and cycle regulate photoperiodic diapause in the bean bug *Riptortus pedestris* males. J Insect Physiol 57:935–938.

63. Meuti ME, Stone M, Ikeno T, Denlinger DL (2015) Functional circadian clock genes are essential for the overwintering diapause of the Northern house mosquito, *Culex pipiens*. J Exp Biol 218:412–422.

64. Mukai A, Goto SG (2016) The clock gene period is essential for the photoperiodic response in the jewel wasp *Nasonia vitripennis* (Hymenoptera: Pteromalidae). Appl Entomol Zool 51:185–194.

65. Tauber E, et al. (2007) Natural selection favors a newly derived timeless allele in *Drosophila* melanogaster. Science 316:1895–1898.

66. Schmidt PS, et al. (2008) An amino acid polymorphism in the couch potato gene forms the basis for climatic adaptation in *Drosophila melanogaster*. Proc Natl Acad Sci USA 105:16207–16211.

67. Bradshaw WE, Holzapfel CM, Mathias D (2006) Circadian rhythmicity and photoperiodism in the pitcher-plant mosquito: can the seasonal timer evolve independently of the circadian clock? Am Nat 167:601–605.

68. Paolucci S, Salis L, Vermeulen CJ, Beukeboom LW, Zande L (2016) QTL analysis of the photoperiodic response and clinal distribution of period alleles in *Nasonia vitripennis*. Mol Ecol 25:4805–4817.

69. Pruisscher P, Nylin S, Gotthard K, Wheat CW (2018) Genetic variation underlying local adaptation of diapause induction along a cline in a butterfly. Mol Ecol 27:3613–3626.

70. Bunning (1936) Endogenous daily rhythms as the basis of photoperiodism. Ber Deut Bot Ges 54:590–607.

71. Bradshaw WE, Holzapfel CM (2010) What season is it anyway? Circadian tracking vs. photoperiodic anticipation in insects. J Biol Rhythms 25:155–165.

72. Emerson KJ, Bradshaw WE, Holzapfel CM (2009) Complications of complexity: integrating environmental, genetic and hormonal control of insect diapause. Trends Genet 25:217–225.

73. Pegoraro M, Gesto JS, Kyriacou CP, Tauber E (2014) Role for circadian clock genes in seasonal timing: testing the Bünning hypothesis. PLoS Genet 10:e1004603.

74. Lankinen, P (1986) Geographical variation in circadian eclosion rhythm and photoperiodic adult diapause in *Drosophila littoralis*. J Comp Physiol A, 159:123–142.

75. Salmela MJ, McMinn RL, Guadagno CR, Ewers BE, Weinig C (2018) Circadian rhythms and reproductive phenology covary in a natural plant population. J Biol Rhythms 33:245–254.

76. Beck SD (1989) Factors influencing the intensity of larval diapause in *Ostrinia nubilalis*. J Insect Physiol 35:75–79.

77. Cloutier EJ, Beck SD, McLeod D, Silhacek DL (1962) Neural transplants and insect diapause. Nature 195:1222.

78. Gilbert LI, Bollenbacher WE, Granger NA (1980) Insect endocrinology: regulation of endocrine glands, hormone titer, and hormone metabolism. Annu Rev Physiol 42:493–510.

79. Gelman DB, et al. (1992) Prothoracicotropic hormone levels in brains of the European corn borer, *Ostrinia nubilalis*: diapause vs the non-diapause state. J Insect Physiol 38:383–395.

80. Wang Q, Mohamed AA, Takeda M (2013) Serotonin receptor B may lock the gate of PTTH release/synthesis in the Chinese silk moth, *Antheraea pernyi*; a diapause initiation/maintenance mechanism? PLoS ONE 8:e79381.

81. Mohamed AA, et al. (2014) N-acetyltransferase (*nat*) is a critical conjunct of photoperiodism between the circadian system and endocrine axis in *Antheraea pernyi*. PLoS ONE 9:e92680.

82. Seluzicki A, et al. (2014) Dual PDF signaling pathways reset clocks via TIMELESS and acutely excite target neurons to control circadian behavior. PLoS Biol 12:e1001810.

83. Levy RC, Kozak GM, Dopman EB (2018) Non-pleiotropic coupling of daily and seasonal temporal isolation in the European corn borer. Genes 9:180.

84. Iga M, Nakaoka T, Suzuki Y, Kataoka H (2014) Pigment Dispersing Factor regulates ecdysone biosynthesis via *Bombyx* neuropeptide G protein coupled receptor-B2 in the prothoracic glands of *Bombyx mori*. PLoS ONE 9:e103239.

85. Ikeno T, Numata H, Goto SG, Shiga S (2014) Involvement of the brain region containing pigment-dispersing factor-immunoreactive neurons in the photoperiodic response of the bean bug, *Riptortus pedestris*. J Exp Biol 217:453–462.

86. Shiga S, Numata H (2009) Roles of PER immunoreactive neurons in circadian rhythms and photoperiodism in the blow fly, *Protophormia terraenovae*. J Exp Biol 212:867–877.

87. Willis CG, et al. (2010) Favorable climate change response explains non-native species success in Thoreau’s woods. PLoS ONE 5:e8878.

88. Sánchez-Bayo F, Wyckhuys KA (2019) Worldwide decline of the entomofauna: A review of its drivers. Biol Conserv 232:8–27.

89. Breed GA, Stichter S, Crone EE (2013) Climate-driven changes in northeastern US butterfly communities. Nat Clim Chang 3:142–145.

90. Walsh J, Wuebbles D, Hayhoe K (2014) Ch. 2: Our Changing Climate. Climate Change Impacts in the United States: The Third National Climate Assessment. US Global Change Research Program. Melillo JM, Richmond T, Yohe G, eds.

91. Williams CM, Henry HA, Sinclair BJ (2015) Cold truths: how winter drives responses of terrestrial organisms to climate change. Biol Rev 90:214–235.

92. Diffenbaugh NS, Krupke CH, White MA, Alexander CE (2008) Global warming presents new challenges for maize pest management. Environ Res Lett 3:044007.

93. Svobodová E, et al. (2014) Determination of areas with the most significant shift in persistence of pests in Europe under climate change. Pest Manag Sci 70:708–715.

94. Coates BS, et al. (2011) The application and performance of single nucleotide polymorphism markers for population genetic analyses of Lepidoptera. Front Genet 2: doi:10.3389/fgene.2011.00038.

95. Broman KW, Wu H, Sen S, Churchill GA (2003) R/qtl: QTL mapping in experimental crosses. Bioinformatics 19:889–890.

96. Broman KW, Sen S (2009) A Guide to QTL Mapping with R/qtl (Springer).

97. R Core Team (2018). R: A language and environment for statistical computing. R Foundation for Statistical Computing, Vienna, Austria. URL https://www.R-project.org/.

98. Endelman JB, Plomion C (2014) LPmerge: an R package for merging genetic maps by linear programming. Bioinformatics 30:1623–1624.

99. Coates BS, et al. (2013) Frequency of hybridization between *Ostrinia nubilalis* E-and Z-pheromone races in regions of sympatry within the United States. Ecol Evol 3:2459–2470.

100. Coates BS, et al. (in review) Influence of host plant, geography and pheromone strain on genomic differentiation in sympatric populations of Ostrinia nubilalis. Mol Ecol.

101. Calvin DD, Song PZ (1994) Variability in postdiapause development periods of geographically separate *Ostrinia nubilalis* (Lepidoptera: Pyralidae) populations in Pennsylvania. Environ Entomol 23:431–436.

102. uz Zaman MF (2008) A comparison of univoltine and multivoltine European corn borer (*Ostrinia nubilalis* Hübner): Life history characters, Bt toxin susceptibility, parasitoid impact, and population pattern. PhD thesis: Penn State University.

103. Waterhouse RM, et al. (2017) BUSCO applications from quality assessments to gene prediction and phylogenomics. Mol Biol Evol 35:543–548.

104. Smit AFA, Hubley R & Green P. RepeatMasker Open-4.0. 2013-2015 <http://www.repeatmasker.org

105. Ben Langmead, Salzberg SL (2012) Fast gapped-read alignment with Bowtie 2. Nature Methods 9:357–359.

106. Li H, et al. (2009) The sequence alignment/map format and SAMtools. Bioinformatics 25:2078–2079.

107. Kofler R, et al. (2011) PoPoolation: A Toolbox for Population Genetic Analysis of Next Generation Sequencing Data from Pooled Individuals. PLoS ONE 6:e15925.

108. Kofler R, Pandey RV, Schlotterer C (2011) PoPoolation2: identifying differentiation between populations using sequencing of pooled DNA samples (Pool-Seq). Bioinformatics 27(:3435–3436.

109. Jeffreys H (1961) Theory of Probability (Clarendon Press, Oxford). 3rd Ed.

110. Vitalis R, Gautier M, Dawson KJ, Beaumont MA (2014) Detecting and measuring selection from gene frequency data. Genetics 196:799–817.

111. McKenna A, et al. (2010) The Genome Analysis Toolkit: a MapReduce framework for analyzing next-generation DNA sequencing data. Genome Res 20:1297–1303.

112. DePristo MA, et al. (2011) A framework for variation discovery and genotyping using next-generation DNA sequencing data. Nature Genet 43:491.

113. Van der Auwera GA, et al. (2013) From FastQ data to high-confidence variant calls: the genome analysis toolkit best practices pipeline. Curr Protoc Bioinformatics:11–10.

114. Rausch T, et al. (2012) DELLY: structural variant discovery by integrated paired-end and split-read analysis. Bioinformatics 28:i333–i339.

115. Browning BL, Zhou Y, Browning SR (2018) A One-Penny Imputed Genome from Next-Generation Reference Panels. Am J Hum Genet 103:338–348.

116. Danecek, Petr, et al (2011) The variant call format and VCFtools. Bioinformatics 27: 2156–2158.

117. Dowle M, Srinivasan A (2018). data.table: Extension of “data.frame”. R package version 1.11.8. https://CRAN.R-project.org/package=data.table

118. Wickham H (2011) The Split-Apply-Combine Strategy for Data Analysis. J Stat Softw 40:1–29.

119. Canty A, Ripley B (2017) boot: Bootstrap R (S-Plus) Functions. R package version 1.3-20.

120. Wickham, H (2017) ggplot2: Elegant Graphics for Data Analysis. Springer-Verlag New York, 2016.

121. Turner, S (2017) qqman: Q-Q and Manhattan Plots for GWAS Data. R package version 0.1.4. https://CRAN.R-project.org/package=qqman

122. Strimmer K (2008) fdrtool: a versatile R package for estimating local and tail area-based false discovery rates. Bioinformatics 24:1461–1462.

123. Levine JD, Funes P, Dowse HB, Hall JC (2002) Signal analysis of behavioral and molecular cycles. BMC Neurosci 3:1.

124. Ko HW, et al. (2010) A hierarchical phosphorylation cascade that regulates the timing of PERIOD nuclear entry reveals novel roles for proline-directed kinases and GSK-3β/SGG in circadian clocks. J Neurosci 30:12664–12675.

125. Zhan S, Merlin C, Boore JL, Reppert SM (2011) The monarch butterfly genome yields insights into long-distance migration. Cell 147:1171–1185.

