## Supplemental Material for "Genomic basis of circannual rhythm in the european corn borer moth"

**Table of contents**

| | | Page |
|---|---|---|
| **Supplemental Methods and Results** | | 2 |
| **Figure S1** | | |
| | The overall effect of reduced generation time on the critical rate of environmental change | 5 |
| **Figure S2** | | 6 |
| | A latitudinal cline in diapause termination timing in North America | |
| **Figure S3** | | 7 |
| | Variation in PDD timing and relation to voltinism | |
| **Figure S4** | | |
| | Sympatric bivoltine (2 generation) and univoltine (1 generation) populations of ECB moth in Farmington, NY | 8 |
| **Figure S5.** | | |
| | Plot of PDD time based on the interaction of alleles at *per* and *Ldh*. | 9 |
| **Figure S6.** | | |
| | Mean $F_{ST}$ within and between PDD types | 10 |
| **Figure S7** | | |
| | $eBP_{is}$ using Z covariance structure for the autosomes including all positions | 11 |
| **Figure S8.** | | |
| | Results using covariance structure for estimated from entire genome | 12 |
| **Figure S9.** | | |
| | Genome-wide CMH BZ-UZ pool-seq all chromosomes | 13 |
| **Figure S10.** | | |
| | Maximum linkage disequilibrium of genes in QTL1 and QTL2 and all gene pairs | 14 |
| **Figure S11.** | | |
| | Heatmap of linkage disequilibrium among genes in QTL1 and QTL2 | 15 |
| **Table S1.** | | |
| | QTL interaction ANOVA table | 16 |
| **Table S2** | | |
| | ECB Genome quality information | 17 |
| **Table S3.** | | |
| | Populations sampled for genomic studies | 18 |
| **Table S4.** | | |
| | Z chromosome scaffolds | 19 |
| **Table S5.** | | |
| | Genes in Figure 3 | 20 |
| **Table S6.** | | |
| | Population covariance from BayPASS | 21 |
| **Table S7.** | | |
| | CMH joint outliers | 22 |
| **Table S8.** | | |
| | Linkage disequilibrium outlier pairs | 23 |
| **Table S9.** | | |
| | Significant association *per* plink case/control analysis. | 24 |
| **Table S10.** | | |
| | Significant association *Pdfr* plink case/control analysis. | 25 |
| **Table S11.** | | |
| | Data plotted in Figure S3. | 29 |

### SUPPLEMENTAL METHODS AND RESULTS

*Genomic location of scaffolds*

Chromosomal positions of genome scaffolds were determined by aligning genotyping-by-sequencing data from pedigree families FQ4, FQ5, and BC4E to the genome scaffolds and constructing linkage groups from identified SNPs in rQTL (1, 2) . Additional Z scaffold data was obtained from blastn searches of markers from previously published linkage maps and pedigree families against the genome scaffolds (2-6). Genes that ran across multiple scaffolds were used to estimate the orientation of each Z scaffold, if this was lacking, we used gene order in the *Bombyx mori* genome (7). Once all scaffolds had been aligned and oriented, approximate Megabase (Mb) location along the chromosome was calculated (20.94 Mb of scaffolds; estimated to be ~99% of the Z chromosome using *Bombyx*). The autosomal map was based on the FQ4 linkage groups (constructed in rQTL using 60 males, maximum LOD = 6, minimum recombination frequency = 0.35). Chromosomes were identified from linkage groups by blasting against BAC sequences used as probes from a fluorescent in-situ hybridization study in ECB that identified all 31 chromosomes relative to their *Bombyx* homologs (6). We further verified chromosomal identity by blasting the sequences against the *Bombyx* genome. *Bombyx* has 3 fused chromosomes (N = 28) relative to ECB (which has the ancestral Lepidopteran karyotype of N=31); *Bombyx* chromosomes 11, 23, and 24 are fusions (6, 8). Other than these fusions, synteny is well conserved even between *Bombyx* and the distantly related fritillary butterfly, *Melitaea cinxia* (N=31; (8)). Because many scaffolds lacked any linkage information in ECB, we used the chromosomal location information for homologous sequences in *Bombyx* for non-fused chromosomes. For fused chromosomes, we used location information from homologous sequences in *Melitaea* rather than *Bombyx* (*Bombyx* chromosome 11 is a fusion of 31 and 12 in *Melitaea*; 24 is a fusion of 29 and 27; 23 is a fusion of 14 and 30). Scaffold orientation was not estimated for autosomal scaffolds. Scaffolds with no assigned chromosomal position remained in our analyses (assigned to "chromosome 32").

*BayPASS with alternate covariance structures*

BayPASS 2.1 (9) was used to identify associated SNPs while controlling for population demography using an updated version of the BayEnv approach (10) to estimate a covariance matrix (omega) that represents an approximation of the unknown demographic history. BayPASS results were robust to the choice of a Z-chromosome or genome-wide covariance matrix (Table S6). *Per* and *Pdfr* had the highest *eBPis* when we specified the Z covariance matrix for autosomal and unscaffolded loci (Figure S6) and *per* and *Pdfr* contained the only 3 loci that showed BF > 10 dB, *eBPis* > 0.53 (99.9999% quantile of pseudo-observed data from 1 million simulated loci) when we used the covariance matrix estimated from all loci (Z and autosomal; Figure S8).

*CMH tests*

We conducted Cochran-Mantel-Haenszel (CMH) tests for consistent differences in allele frequency in SNPs among independent fast-slow populations pairs (11). P-values from all genomic loci and all possible independent pairwise comparisons were tested separately (6 total 2 x 2 comparisons) for multiple testing using the false discovery rate using the fdrtools package in R. Only SNPs with non-significant values for Woolf heterogeneity test were considered outliers (12) . The Z chromosome contained the most strongly significant SNPs for the CMH test (minimum $q < 10^{-17}$, Figure S9) and the highest number of joint outlier SNPs in fast-slow population comparisons (Chr Z = 37, Chr 3 = 1, Chr 4 = 2, Chr10=1, Chr13=1, Chr17 = 2). Of these joint outliers, 14 were intergenic, the remaining 30 were predicted to be located in 12 different genes; 11 of which were on the Z chromosome (Table S7). Annotated CMH joint outlier genes located under Q1 included period (*per*) and GTPase activating protein (*GAPsec*). Annotated outlier genes located under Q2 included SNF4/AMP-activated protein kinase gamma subunit (SNF4$\gamma$), pigment-dispersing factor receptor (*Pdfr*), terribly reduced optic lobes (*trol*), and coiled-coil domain containing protein (*CG32809*) (Table S7).

*Gene Annotation*

Predictions of protein coding genes were derived from RNA sequencing (RNA-seq) reads mapped to *O. nubilalis* genome assembly using Bowtie2-Tophat2-cufflinks pipeline (13). RNA-seq data were obtained from a colony of bivoltine individuals derived from collection in Ames, Iowa (larval midgut: (14); embryos, pupae, and whole male and female moths: Coates *unpublished data*) and the univoltine BV colony (larval heads: (15); adult female heads: (16); whole pupae: (Wadsworth & Dopman *unpublished*). Gene models were generated by alignment of *de novo* assembled transcripts to the genome scaffolds using GMAP-GSNAP (17, 18). GMAP-GSNAP was trained using all available Lepidopteran model species in Lepbase (release 2; accessed 2016) prior to building gene models (19). A condensed set of genes was generated by clustering transcripts produced from a single locus, and accomplished using CD-HIT-EST (95% similarity cutoff; (20)). This resulted in 27,830 predicted peptide coding regions (Table S3). We also constructed transcriptomes *de novo* in Trinity (following methods in Levy et al. 2018 (16)) to identify all splice variants as well as exons that might be present in scaffold gaps and therefore not represented in the genome. *De novo* transcripts were then blasted against the genome. Regions present on ends transcripts but not containing open reading frames were annotated as 5' or 3' UTR based on their positions. Start and stop codons were the first codon of their kind in a transcript's open reading frame (ORF). Protein domains were annotated using NCBI's conserved domain database (21) and Prosite annotations from PredictProtein (https://www.predictprotein.org, accessed 4/2/2018). Changes in splice sites were predicted using *Drosophila* predictions from BDGP (http://www.fruitfly.org/seq_tools/splice.html; (22)). Locations within proteins of interest are listed by the amino acid number in the homologous *B. mori* protein (BM-AA) as determined by alignment in clustal omega (23). For amino acid comparisons among insect species, protein sequences were downloaded from uniprot, lepbase (release 4, accessed 2018) (19), and Genbank. ECB sequences were aligned using MUSCLE and Jalview was used for visualization (24, 25).

Using homology with *per* in the silkworm *Bombyx mori*, we found that exons 5-6 corresponded to the PAS domains (TIM binding sites), exon 14 contains the DOUBLETIME (DBT) phosphorylation sites, and exons 18-20 contained the CLK-CYC inhibitory domain (CCID) in the C-terminal of *per* (26). Two nuclear localization sequences (NLS) were detected using PredictProtein (https://www.predictprotein.org, accessed 4/2/2018); one near the beginning of the protein and one near the end. Overall, 13 protein kinase C, 34 casein kinase 2 (CK2), 2 cAMP/cGMP dependent protein kinase, and 2 tyrosine kinase phosphorylation sites were predicted across the protein. Like other Lepidopteran sequences, but unlike *Drosophila*, *per* in ECB did not contain T-G repeats upstream of the DBT sites. RNA-seq alignment indicated several splice variants of *per*. Exon 7 sometimes lacked the 5 terminal amino acids (GTQKK); this region likely contains a protein kinase C phosphorylation site and matches a splice variant reported in *Bombyx mori* (27). Exon 21 and exon 22 could each be spliced out. Exon 26 could be missing 2 terminal amino acids (RV). Only the RV variant differed in its presence among PDD types. We never detected the RV-containing splice variant in slow PDD libraries, but it was present in fast PDD libraries (N = 31 of each type). This RV deletion was near the predicted terminal NLS sequence. A SNP in the 5'UTR intron was predicted to remove a predicted splice site in the slow variant; however, a splice variant was not detected in RNA-seq data. This SNP had the most significant association ($q < 1.34 \times 10^{-7}$) and it was the same variant identified in the pool-seq data (scaffold532:93691; Table S9).

*Circadian activity*
Circadian rhythm index (RI) did not significantly differ among fast and slow PDD groups ($F_{1, 56} = 1.35$, $p = 0.24$) nor between replicates ($F_{1, 56} = 0.014$, $p = 0.91$). After removal of arrhythmic males possessing RI values < 0.1 (N = 7 slow PDD males, N = 6 fast PDD males), estimated activity period ($\tau$) significantly differed among fast and slow PDD groups ($F_{1, 43} = 13.48$, $p < 0.001$), but not between replicates ($F_{1, 43} = 0.01$, $p = 0.92$) and therefore replicates were combined in the final analysis.

### SUPPLEMENTAL REFERENCES

1. Coates BS, Siegfried BD (2015) Linkage of an ABCC transporter to a single QTL that controls *Ostrinia nubilalis* larval resistance to the Bacillus thuringiensis Cry1Fa toxin. *Insect Biochem Mol Biol* 63:86–96.
2. Kozak GM, et al. (2017) A combination of sexual and ecological divergence contributes to rearrangement spread during initial stages of speciation. *Mol Ecol* 26:2331–2347.
3. Kroemer JA, et al. (2011) A rearrangement of the Z chromosome topology influences the sex-linked gene display in the European corn borer, *Ostrinia nubilalis. Mol Genet Genomics* 286:37–56.
4. Levy RC, Kozak GM, Wadsworth CB, Coates BS, Dopman EB (2015) Explaining the sawtooth: latitudinal periodicity in a circadian gene correlates with shifts in generation number. *J Evol Biol* 28:40–53.
5. Koutroumpa FA, Groot AT, Dekker T, Heckel DG (2016) Genetic mapping of male pheromone response in the European corn borer identifies candidate genes regulating neurogenesis. *Proc Natl Acad Sci USA*:201610515.
6. Yasukochi Y, et al. (2016) A FISH-based chromosome map for the European corn borer yields insights into ancient chromosomal fusions in the silkworm. *Heredity* 116:75.
7. Consortium ISG (2008) The genome of a lepidopteran model insect, the silkworm *Bombyx mori. Insect Biochem Mol Biol* 38:1036–1045.
8. Ahola V, et al. (2014) The Glanville fritillary genome retains an ancient karyotype and reveals selective chromosomal fusions in Lepidoptera. *Nature Comm* 5:1–9.
9. Gautier M (2015) Genome-wide scan for adaptive divergence and association with population-specific covariates. *Genetics* 201:1555–1579.
10. Günther T, Coop G (2013) Robust identification of local adaptation from allele frequencies. *Genetics*195:205-220.
11. Bastide H, et al. (2013) A genome-wide, fine-scale map of natural pigmentation variation in *Drosophila melanogaster. PLoS Genet* 9:e1003534.
12. Wiberg RAW, Gaggiotti OE, Morrissey MB, Ritchie MG (2017) Identifying consistent allele frequency differences in studies of stratified populations. *Methods Ecol Evol* 8(12):1899–1909.
13. Kim D, et al. (2013) TopHat2: accurate alignment of transcriptomes in the presence of insertions, deletions and gene fusions. *Genome Biol* 14:R36.
14. Vitalis R, Gautier M, Dawson KJ, Beaumont MA (2014) Detecting and measuring selection from gene frequency data. *Genetics* 196:799–817.
15. Wadsworth CB, Dopman EB (2015) Transcriptome profiling reveals mechanisms for the evolution of insect seasonality. *J Exp Biol* 218:3611–3622.
16. Levy RC, Kozak GM, Dopman EB (2018) Non-Pleiotropic Coupling of Daily and Seasonal Temporal Isolation in the European Corn Borer. *Genes* 9:180.
17. Wu TD, Watanabe CK (2005) GMAP: a genomic mapping and alignment program for mRNA and EST sequences. *Bioinformatics* 21:1859–1875.
18. Wu TD, Nacu S (2010) Fast and SNP-tolerant detection of complex variants and splicing in short reads. *Bioinformatics* 26:873–881.
19. Challis RJ, Kumar S, Dasmahapatra KKK, Jiggins CD, Blaxter M (2016) Lepbase: the Lepidopteran genome database. *bioRxiv*:056994.
20. Huang Y, Niu B, Gao Y, Fu L, Li W (2010) CD-HIT Suite: a web server for clustering and comparing biological sequences. *Bioinformatics* 26:680–682.
21. Marchler-Bauer A, et al. (2016) CDD/SPARCLE: functional classification of proteins via subfamily domain architectures. *Nucleic Acids Res* 45:D200–D203.
22. Reese MG, Eeckman FH, Kulp D, Haussler D (1997) Improved splice site detection in Genie. *J Comput Biol* 4:311–323.
23. Sievers F, Higgins DG (2014) Clustal Omega, accurate alignment of very large numbers of sequences. *Multiple Sequence Alignment Methods* (Springer), pp 105–116.
24. Edgar RC (2004) MUSCLE: multiple sequence alignment with high accuracy and high throughput. *Nucleic Acids Res* 32:1792–1797.
25. Waterhouse AM, Procter JB, Martin DM, Clamp M, Barton GJ (2009) Jalview Version 2—a multiple sequence alignment editor and analysis workbench. *Bioinformatics* 25:1189–1191.
26. Chang DC, Reppert SM (2003) A novel C-terminal domain of *Drosophila* PERIOD inhibits dCLOCK: CYCLE-mediated transcription. *Curr Biol* 13:758–762.
27. Takeda Y, et al. (2004) Structural analysis and identification of novel isoforms of the circadian clock gene period in the silk moth *Bombyx mori. Zool Sci* 21:903–915.
28. Chevin L-M, Lande R, Mace GM (2010) Adaptation, Plasticity, and Extinction in a Changing Environment: Towards a Predictive Theory. *PLoS Biol* 8:e1000357.
29. Showers WB (1979) Effect of diapause on the migration of the European corn borer into the southeastern United States. In: Movement of highly mobile insects: concepts and methodology in research; proceedings of a conference (Raleigh, NC, University Graphics, 1979).
30. Wadsworth CB, Woods WA Jr, Hahn DA, Dopman EB (2013) One phase of the dormancy developmental pathway is critical for the evolution of insect seasonality. *J Evol Biol* 26:2359–2368.
31. McLeod DGR (1976) Geographical variation of diapause termination in the European corn borer, *Ostrinia nubilalis* (Lepidoptera: Pyralidae), in southwestern Ontario. *Can Entomol* 108:1403-1408.
32. McLeod DGR, Ritchot C, Nagai T (1979). Occurrence of a two generation strain of the European corn borer, *Ostrinia nubilalis* (Lepidoptera: Pyralidae), in Quebec. *Can Entomol* 111:233-236.
33. Glover TJ, Robbins PS, Eckenrode CJ, Roelofs WL (1992) Genetic control of voltinism characteristics in European corn borer races assessed with a marker gene. *Arch Insect Biochem Physiol* 20:107–117.
34. Dopman EB, Perez L, Bogdanowicz SM, Harrison RG (2005) Consequences of reproductive barriers for genealogical discordance in the European corn borer. *Proc Natl Acad Sci USA* 102:14706–14711.

**Figure S1. The overall effect of reduced generation time on the critical rate of environmental change**. Data is shown for several values of voltinism. Depicted is the relative increase in allowable environmental change possible with each added generation. Values of other model parameters given in Figure 2 of Chevin et al (2010) (28).

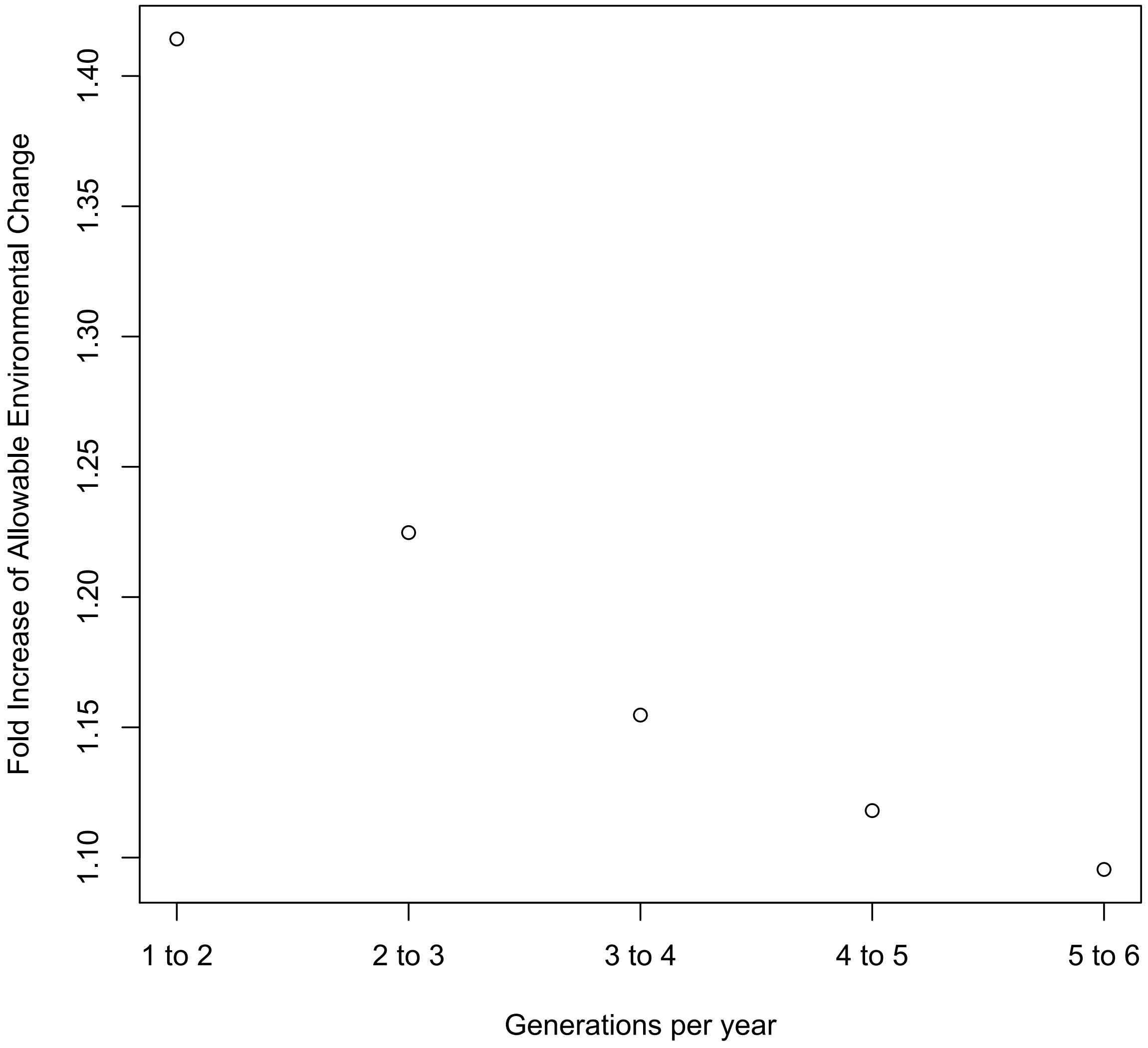

**Figure S2. A latitudinal cline in diapause termination timing in North America**. Mean number of days until 50% pupation of sample in European corn borer reared at 28ºC and 16 h light based on latitude. Data from Showers 1979 (29), measurements made 1947-50. Pearson correlation $r = 0.72$, P = 0.03. State abbreviations noted.

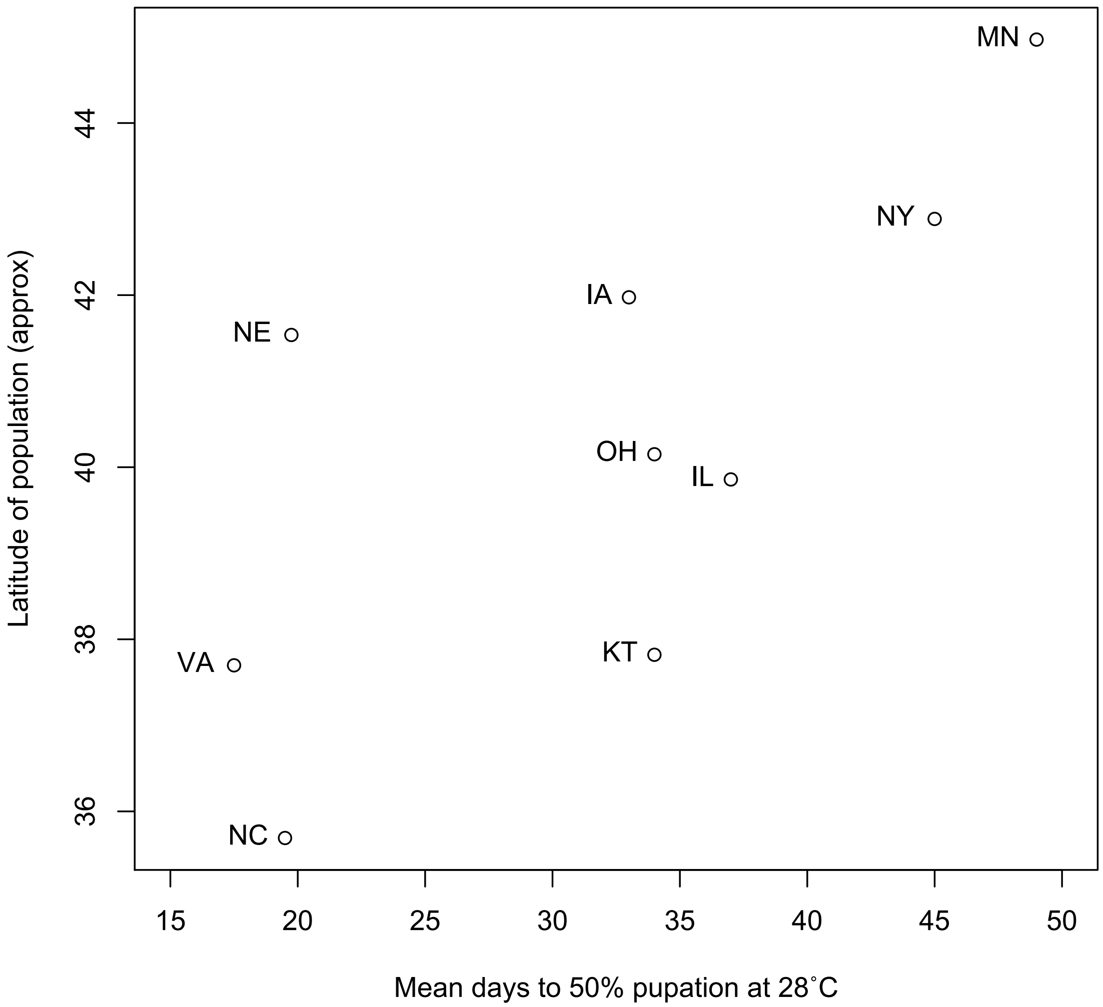

**Figure S3. Variation in PDD timing and relation to voltinism**. Mean number of days to pupation under 16 h light at 30ºC (filled symbols) and 26ºC (open symbols) for populations that are univoltine or bivoltine in the field. Bars indicate standard error. Data sources listed in Table S11.

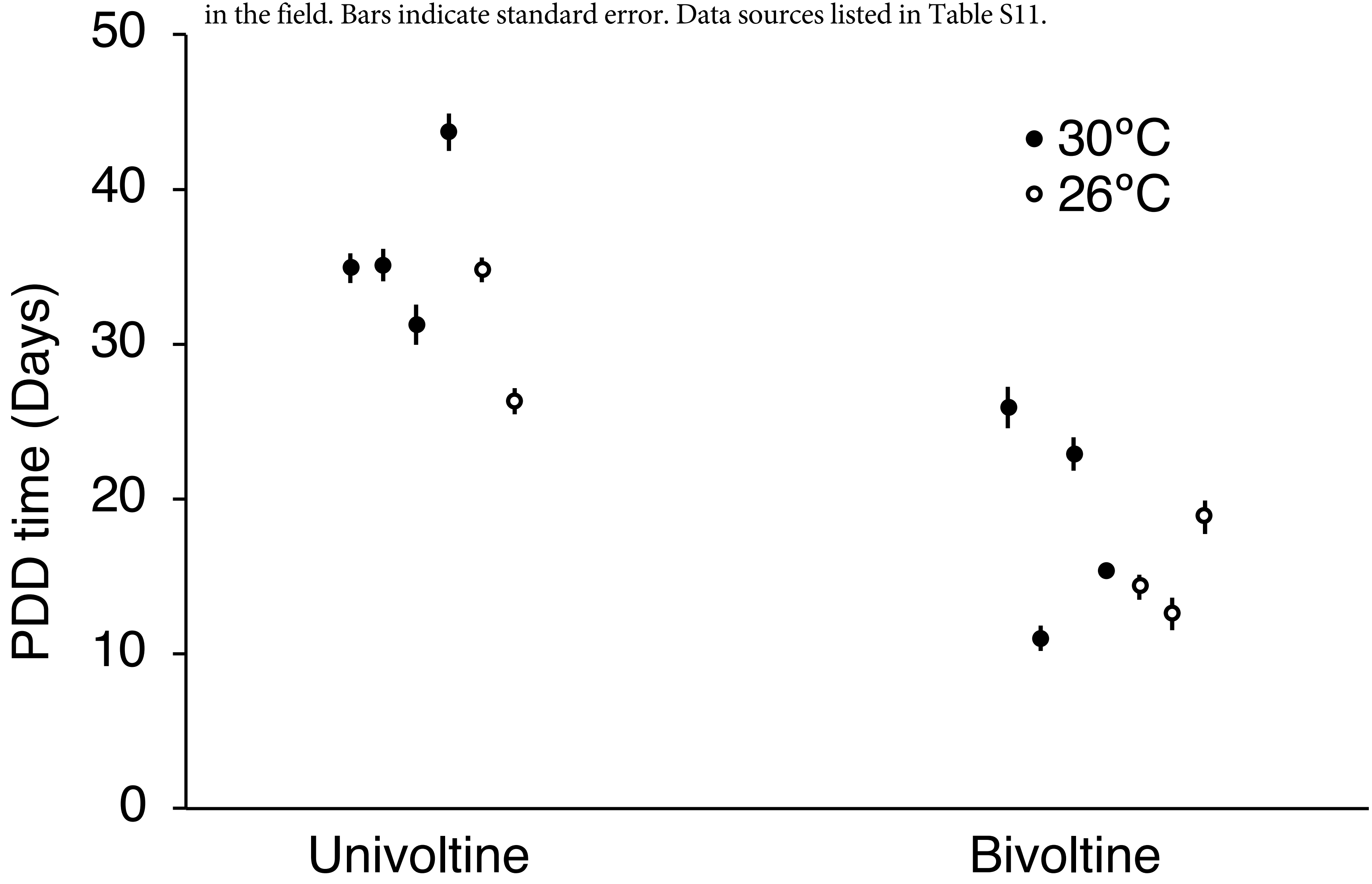

**Figure S4. Sympatric bivoltine (2 generation) and univoltine (1 generation) populations of ECB moth in Farmington, NY.** Data adapted from Wadsworth et al. 2013 (30).

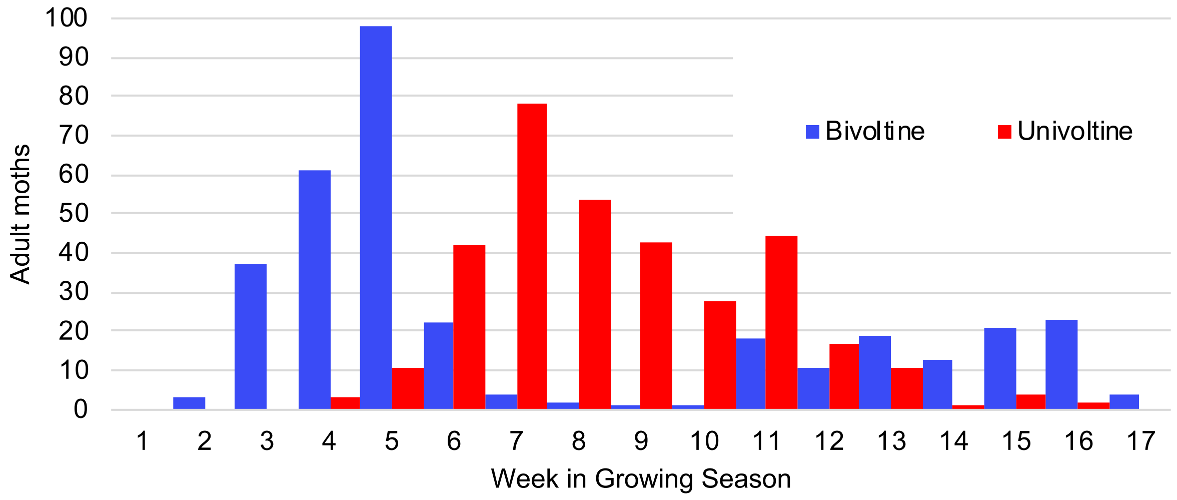

**Figure S5**. **Plot of PDD time based on the interaction of alleles at *per* and *Ldh***. Females with a long allele at *per* but a short allele at *Ldh* have longer PDD time than those with a long allele at *Ldh*.

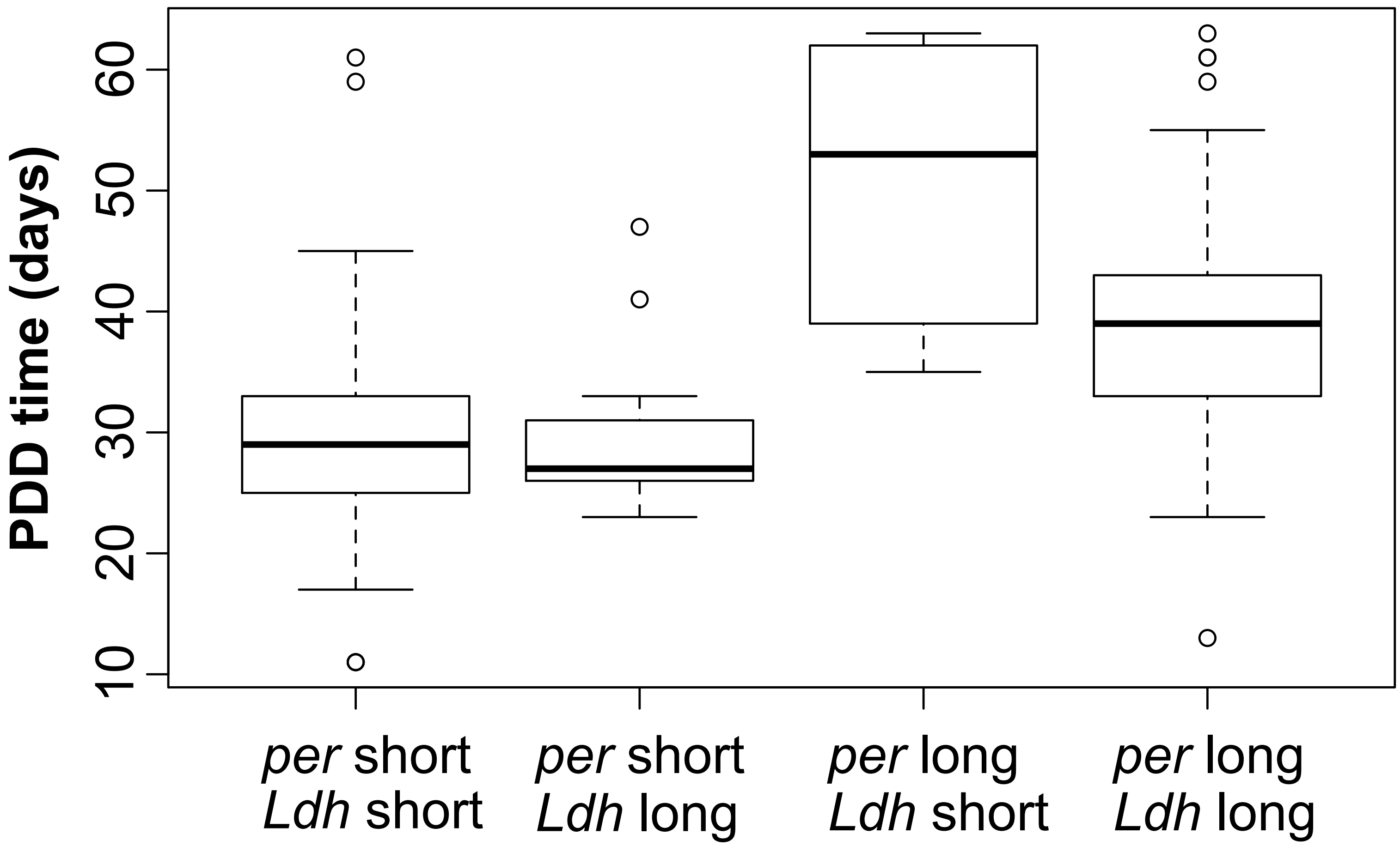

**Figure S6. Mean $F_{ST}$ within and between PDD types.** (a) Mean $F_{ST}$ between PDD type on the Z chromosome (position in Mb) for 1 kb windows within genes. Windows $F_{ST} \geq 0.5$ highlighted in red (occur in the 5'UTR of *per* and *Pdfr*). (b) Mean $F_{ST}$ within PDD type on the Z chromosome 1 kb windows within genes.

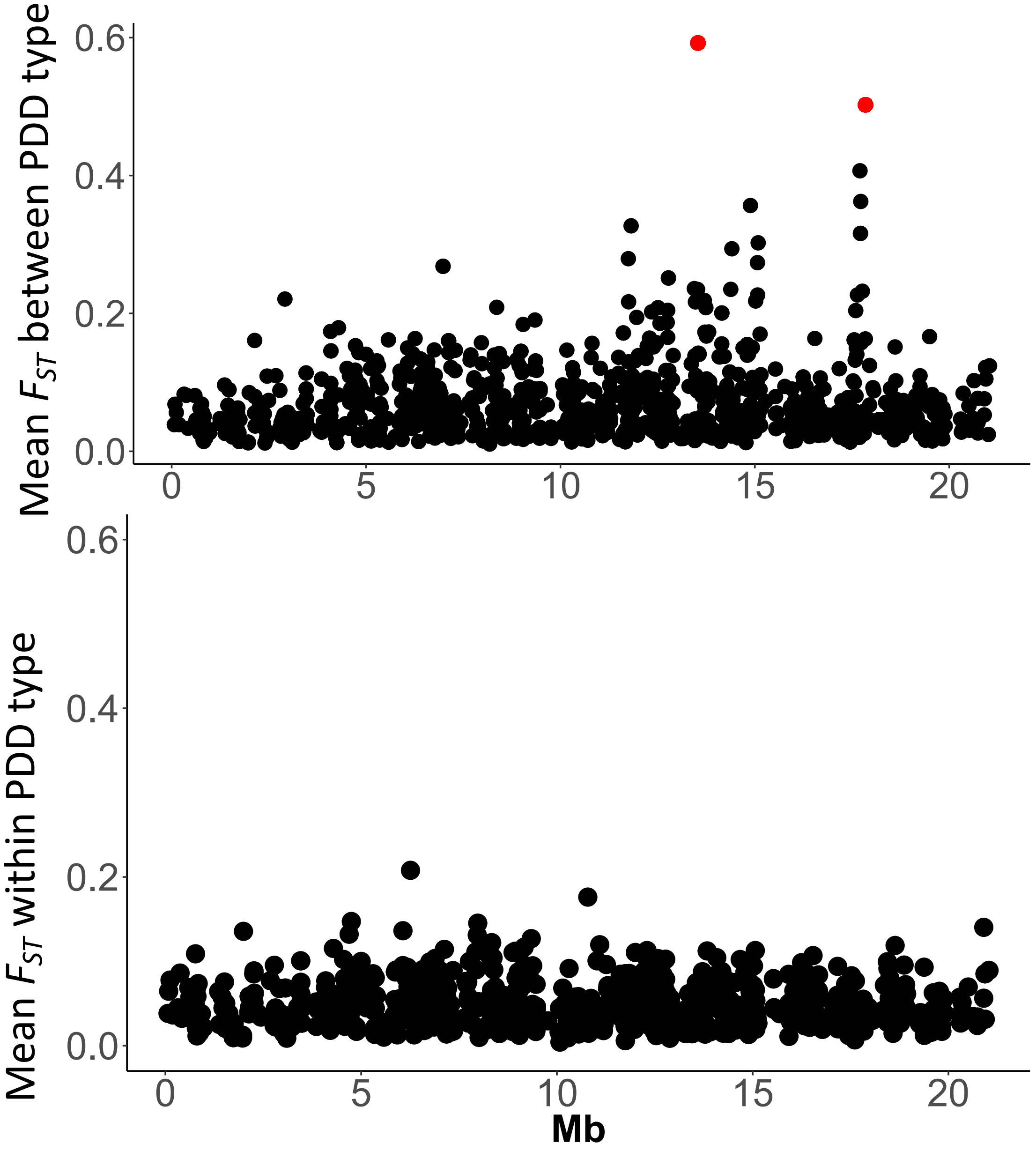

**Figure S7. *$eBP_{is}$* using Z covariance structure for the autosomes including all positions.** Intergenic regions included. No outliers with *$eBP_{is}$* > 2 detected outside of the Z chromosome.

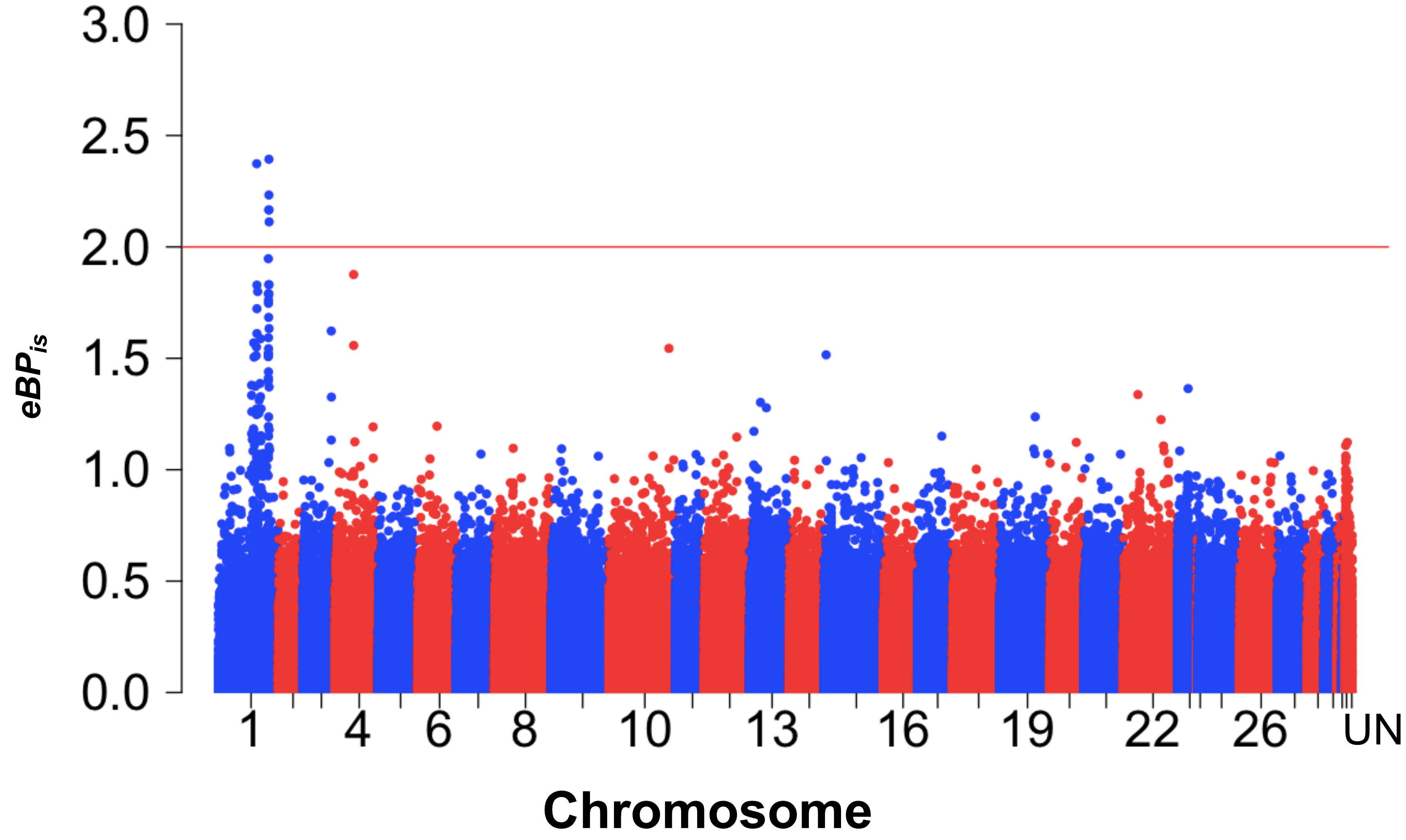

**Figure S8. Results using covariance structure for estimated from entire genome**. (a) eBPis and (b) Absolute value of beta from BayPASS analysis using omega estimated from entire genome (Z and Autosomes combined; all sites). Red line indicates the 99.999% quantile from a simulated pseudo-observed dataset (POD) of 1M SNPs. *Per* and *Pdfr* has the highest $eBP_{is}$ (> 0.75) and the maximum beta (> 0.0139), and were the only loci in the genome with beta in the 99.999% quantile and with BF > 10 *dB*.

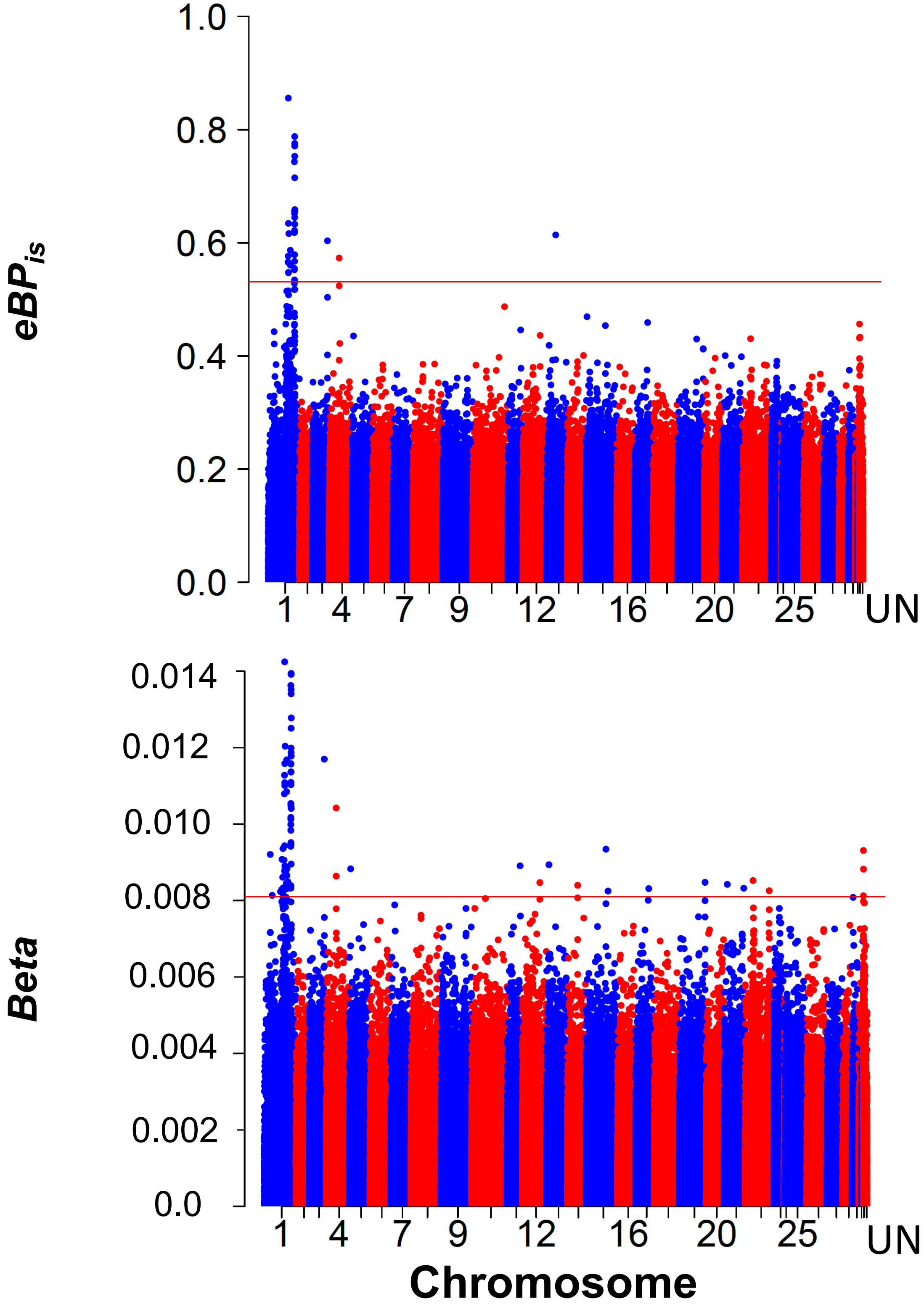

**Figure S9. Genome-wide CMH BZ-UZ Pool-Seq all chromosomes**. FDR corrected q-values for EA vs. BV and LA vs. GEN comparison shown. Chromosomes 1(Z)-31, with Unscaffolded at the end.

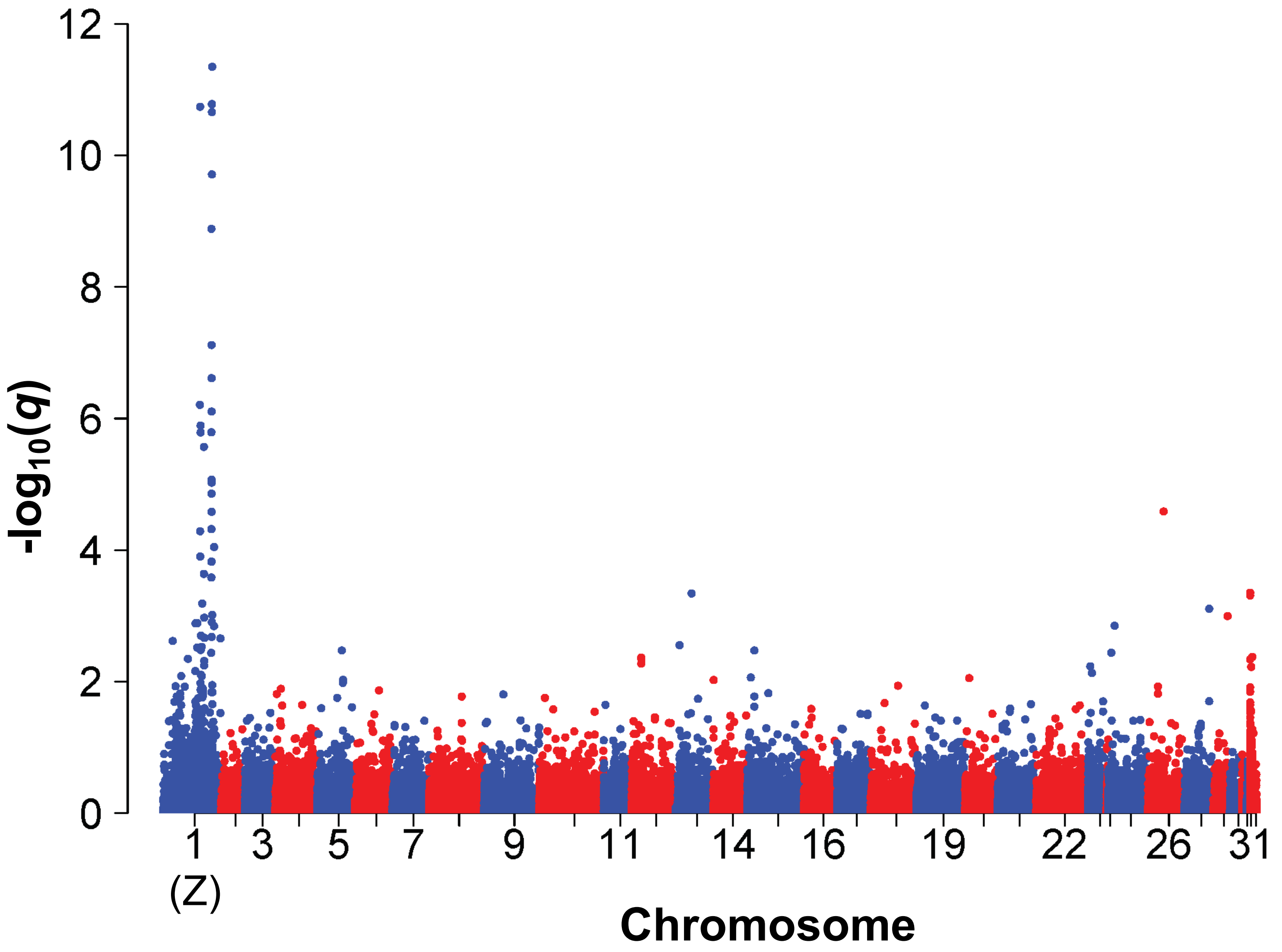

**Figure S10. Maximum linkage disequilibrium all gene pairs.** LD (measured by $r^2$) observed between gene pairs. Red line represents the upper limit of the 95% confidence interval for the 99.9% quantile for maximum LD as estimated by bootstrapping at 1 Mb intervals. LD outliers are above this line. red point indicates *per-Pdfr* (a) all gene pairs located within/between QTL1 and QTL2 with 2 Mb ≤ estimated distance ≤ 7 Mb. (b) All Z chromosome genes 2 Mb ≤ estimated distance ≤ 6 Mb. Open points indicate genes where only one gene is within a QTL region or neither gene is in a QTL. Blue points indicated gene pairs in QTL1 and QTL2. The only other gene pair with comparable LD to *per-Pdfr* is *dunce-SNF4γ (dunce* is not under the QTL, *SNF4γ* is in QTL2).

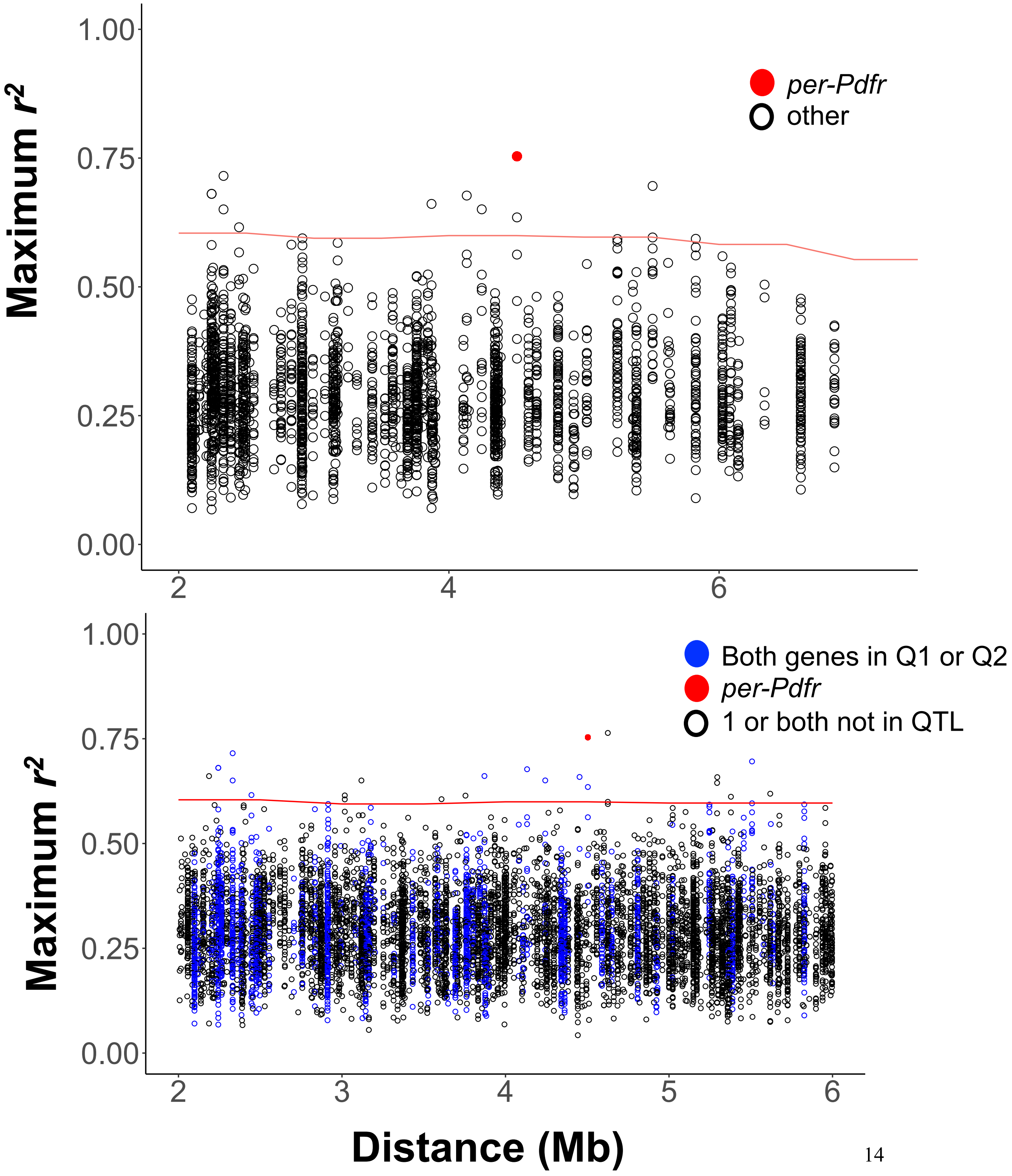

**Figure S11. Heatmap of linkage disequilibrium among genes in QTL1 and QTL2.** Heatmap of the maximum linkage disequilibrium (LD, measured by $r^2$) vs. mean distance between gene pairs in QTL1 and QTL2.

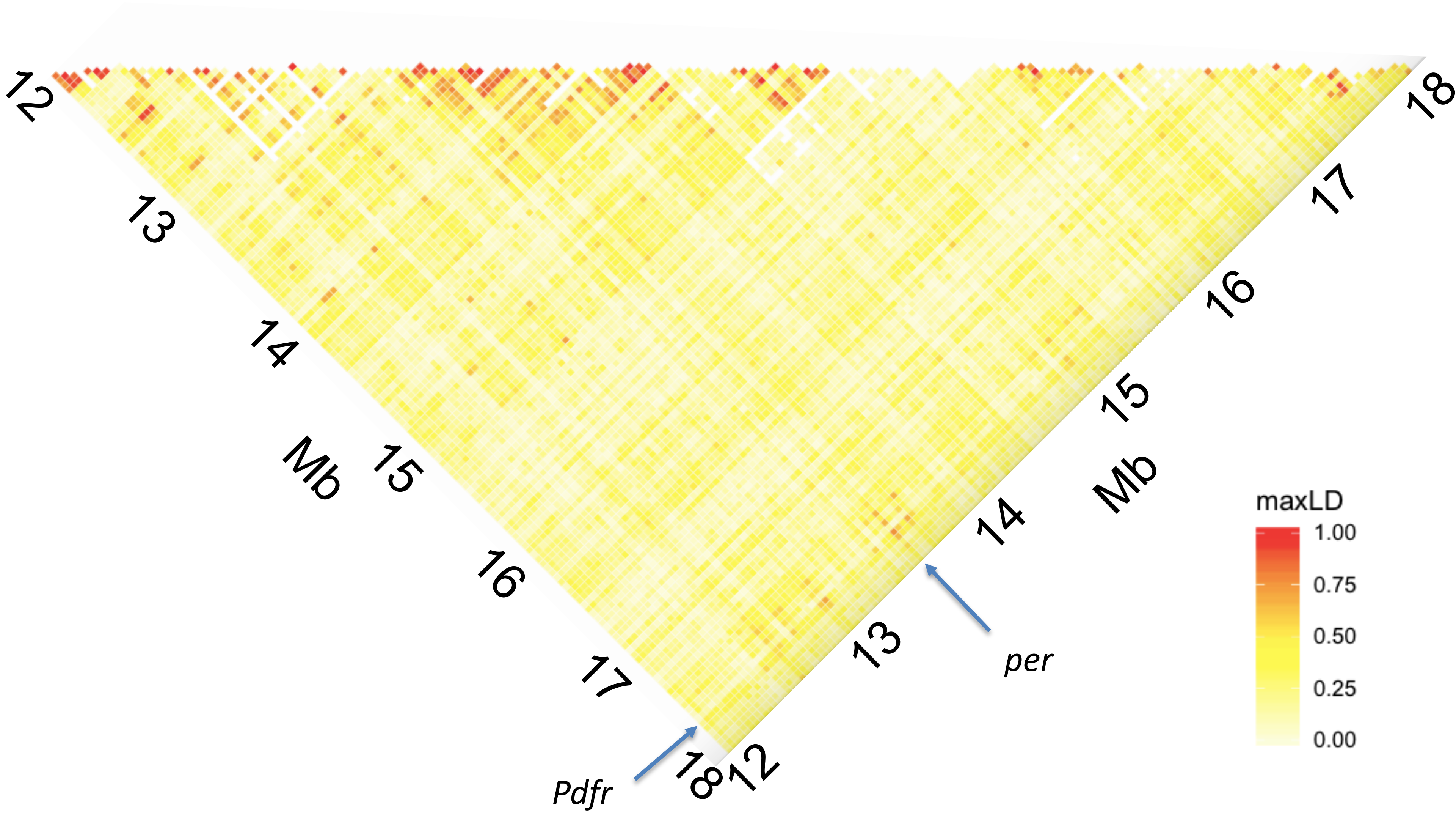

**Table S1. QTL interaction**

**A. QTL interaction EA x BV pedigree females (position in cM)**

y~Q1+Q2+Q1:Q2

| | df | SS | MS | LOD | %var | P |
|---|---|---|---|---|---|---|
| Model | 3 | 7401.44 | 2467.14655 | 21.60897 | 35.61715 | 0 |
| Error | 222 | 13379.11 | 60.26626 | | | |
| Total | 225 | 20780.55 | | | | |

| | df | Type III SS | LOD | %var | F value | P |
|---|---|---|---|---|---|---|
| 1@29.5 | 2 | 2759.2 | 9.202 | 13.278 | 22.892 | 9.14E-10 |
| 1@34.0 | 2 | 1095.1 | 3.861 | 5.27 | 9.086 | 0.000161 |
| 1@29.5:1@34.0 | 1 | 469.9 | 1.694 | 2.261 | 7.797 | 0.005689 |

**B. Marker genotype interaction EA x BV pedigree females**

| | DF | SS | F | P |
|---|---|---|---|---|
| (Intercept) | 1 | 99150 | 1643.9807 | 2.20E-16 |
| *per* | 1 | 2216 | 36.7422 | 5.87E-09 |
| *Ldh* | 1 | 21 | 0.347 | 0.55642 |
| *per:Ldh* | 1 | 528 | 8.7599 | 0.00342 |
| Residuals | 218 | 13148 | | |

**Table S2. ECB Genome quality information**

| BUSCO | Number | Percent |
|---|---|---|
| Total complete BUSCOs | 993 | 93.10% |
| Complete and single-copy BUSCOs | 983 | 92.20% |
| Complete and duplicated BUSCOs | 10 | 0.90% |
| Fragmented BUSCOs (F) | 35 | 3.30% |
| Missing BUSCOs (M) | 38 | 3.60% |
| **Total BUSCO groups searched** | **1066** | |
| Total scaffolds | 8,843 | |
| Total gene models | 27,830 | |
| Scaffold N50 | 392.5 kb | |
| Maximum Scaffold Size | 3.32 Mb | |
| Total sequenced genome | 454.7 Mb | |
| Total assigned chromosomes | 278 | 61% |

**Table S3. Populations sampled for genomic studies**

| Field populations | State | PDD time | Generations/year | Pool-seq | Indiv-seq | Collection Date | Abbreviation | Latitude, Longitude |
|---|---|---|---|---|---|---|---|---|
| East Aurora | NY | Short | Two | 34 | 11 | 2011 | EA | 42.7678° N, 78.6134° W |
| Landisville | PA | Short | Two | 39 | 14 | 2013 | LA | 40.0953° N, 76.4103° W |
| Bouckville-Madison | NY | Long | One | 20 | 4 | 2000 | BV | 42.8990° N, 75.5121° W |
| Penn Yan | NY | Long | One | 26 | 4 | 2008 | PY | 42.6609° N, 77.0539° W |
| Geneva | NY | Long | One | 25 | 10 | 2000<br>2004<br>2007 | GEN | 42.8680° N, 76.9856° W |

| Table S4. Z chromosome scaffolds | | | | | | | | | |
|---|---|---|---|---|---|---|---|---|---|
| **Scaffold.Full** | **size** | **LG** | **ECBChr** | **BombyxChr** | **McinChr** | **Direction** | **Start.Mb.est** | **Marker** | ***QTL*** |
| scaffold4435\|size4078 | 4078 | Z | 1 | 1 | 1 | F | 0.000 | | |
| scaffold751\|size275148 | 275148 | Z | 1 | 1 | 1 | F | 0.004 | | |
| scaffold35\|size722381 | 722381 | Z | 1 | 1 | 1 | F | 0.279 | *Ket* | |
| scaffold740\|size235087 | 235087 | Z | 1 | 1 | 1 | R | 1.165 | | |
| scaffold1546\|size33109 | 33109 | Z | 1 | 1 | 1 | F | 1.400 | | |
| scaffold247\|size627634 | 627634 | Z | 1 | 1 | 1 | F | 1.434 | *alpha* | |
| scaffold196\|size868841 | 868841 | Z | 1 | 1 | 1 | R | 2.061 | | |
| scaffold779\|size92251 | 92251 | Z | 1 | 1 | 1 | F | 2.930 | | |
| scaffold2358\|size14287 | 14287 | Z | 1 | 1 | 1 | F | 3.022 | | |
| scaffold600\|size417131 | 417131 | Z | 1 | 1 | 1 | F | 3.037 | | |
| scaffold216\|size461870 | 461870 | Z | 1 | 1 | 1 | R | 3.454 | | |
| scaffold390\|size298204 | 298204 | Z | 1 | 1 | 1 | F | 3.916 | | |
| scaffold364\|size585662 | 585662 | Z | 1 | 1 | 1 | F | 4.214 | | |
| scaffold1895\|size21730 | 21730 | Z | 1 | 1 | 1 | F | 4.799 | | |
| scaffold409\|size222884 | 222884 | Z | 1 | 1 | 1 | F | 4.821 | | |
| scaffold444\|size384115 | 384115 | Z | 1 | 1 | 1 | R | 5.044 | | |
| scaffold325\|size886211 | 886211 | Z | 1 | 1 | 1 | F | 5.428 | | |
| scaffold15\|size673834 | 673834 | Z | 1 | 1 | 1 | R | 6.314 | | |
| scaffold169\|size242750 | 242750 | Z | 1 | 1 | 1 | R | 6.988 | *Desert* | |
| scaffold348\|size845032 | 845032 | Z | 1 | 1 | 1 | F | 7.231 | *Cyp303a1* | |
| scaffold865\|size221853 | 221853 | Z | 1 | 1 | 1 | R | 8.076 | | |
| scaffold518\|size233254 | 233254 | Z | 1 | 1 | 1 | R | 8.298 | | |
| scaffold2225\|size15954 | 15954 | Z | 1 | 1 | 1 | F | 8.531 | | |
| scaffold546\|size598492 | 598492 | Z | 1 | 1 | 1 | F | 8.547 | *Tpi* | |
| scaffold645\|size139123 | 139123 | Z | 1 | 1 | 1 | F | 9.146 | | |
| scaffold1618\|size30317 | 30317 | Z | 1 | 1 | 1 | F | 9.285 | | |
| scaffold789\|size91266 | 91266 | Z | 1 | 1 | 1 | R | 9.315 | | |
| scaffold218\|size453784 | 453784 | Z | 1 | 1 | 1 | F | 9.406 | | |
| scaffold1660\|size63095 | 63095 | Z | 1 | 1 | 1 | F | 9.860 | | |
| scaffold1044\|size193178 | 193178 | Z | 1 | 1 | 1 | F | 9.923 | | |
| scaffold581\|size122005 | 122005 | Z | 1 | 1 | 1 | R | 10.116 | | |
| scaffold1237\|size50895 | 50895 | Z | 1 | 1 | 1 | F | 10.238 | | |
| scaffold1246\|size194460 | 194460 | Z | 1 | 1 | 1 | F | 10.289 | | |
| scaffold1756\|size66030 | 66030 | Z | 1 | 1 | 1 | F | 10.484 | | |
| scaffold1161\|size202793 | 202793 | Z | 1 | 1 | 1 | F | 10.550 | | |
| scaffold155\|size350779 | 350779 | Z | 1 | 1 | 1 | F | 10.752 | | |
| scaffold282\|size321509 | 321509 | Z | 1 | 1 | 1 | R | 11.103 | | |
| scaffold502\|size248588 | 248588 | Z | 1 | 1 | 1 | R | 11.425 | *710* | *QTL1* |
| scaffold804\|size458571 | 458571 | Z | 1 | 1 | 1 | F | 11.673 | | *QTL1* |
| scaffold3599\|size6641 | 6641 | Z | 1 | 1 | 1 | R | 12.132 | *AcylB* | *QTL1* |
| scaffold1040\|size105542 | 105542 | Z | 1 | 1 | 1 | R | 12.139 | | *QTL1* |
| scaffold4204\|size4752 | 4752 | Z | 1 | 1 | 1 | F | 12.244 | | *QTL1* |
| scaffold4243\|size4645 | 4645 | Z | 1 | 1 | 1 | F | 12.249 | | *QTL1* |
| scaffold275\|size1004024 | 1004024 | Z | 1 | 1 | 1 | R | 12.253 | *clock* | *QTL1* |
| scaffold532\|size371787 | 371787 | Z | 1 | 1 | 1 | R | 13.258 | *per* | *QTL1* |
| scaffold459\|size265770 | 265770 | Z | 1 | 1 | 1 | F | 13.629 | | *QTL1* |
| scaffold1889\|size21771 | 21771 | Z | 1 | 1 | 1 | F | 13.895 | | *QTL1* |
| scaffold221\|size590889 | 590889 | Z | 1 | 1 | 1 | R | 13.917 | *13318* | *QTL1* |
| scaffold931\|size77297 | 77297 | Z | 1 | 1 | 1 | F | 14.508 | | |
| scaffold140\|size987430 | 987430 | Z | 1 | 1 | 1 | R | 14.585 | *12295* | *QTL2* |
| scaffold1038\|size208510 | 208510 | Z | 1 | 1 | 1 | F | 15.572 | | *QTL2* |
| scaffold598\|size233855 | 233855 | Z | 1 | 1 | 1 | F | 15.781 | | *QTL2* |
| scaffold48\|size965936 | 965936 | Z | 1 | 1 | 1 | F | 16.015 | | *QTL2* |
| scaffold1486\|size80768 | 80768 | Z | 1 | 1 | 1 | F | 16.981 | | *QTL2* |
| scaffold193\|size439059 | 439059 | Z | 1 | 1 | 1 | F | 17.062 | *Ldh* | *QTL2* |
| scaffold137\|size260670 | 260670 | Z | 1 | 1 | 1 | R | 17.501 | | *QTL2* |
| scaffold87\|size517236 | 517236 | Z | 1 | 1 | 1 | F | 17.761 | | *QTL2* |
| scaffold50\|size611115v02 | 357676 | Z | 1 | 1 | 1 | R | 18.278 | *12238* | |
| scaffold662\|size413436 | 413436 | Z | 1 | 1 | 1 | R | 18.636 | | |
| scaffold728\|size179971v02 | 161281 | Z | 1 | 1 | 1 | F | 19.050 | | |
| scaffold229\|size374464 | 374464 | Z | 1 | 1 | 1 | R | 19.211 | *12360* | |
| scaffold85\|size326585 | 326585 | Z | 1 | 1 | 1 | F | 19.585 | | |
| scaffold882\|size106910 | 106910 | Z | 1 | 1 | 1 | F | 19.912 | | |
| scaffold330\|size554422 | 554422 | Z | 1 | 1 | 1 | F | 20.019 | *Sh* | |
| scaffold673\|size283851 | 283851 | Z | 1 | 1 | 1 | R | 20.573 | *EIPB* | |
| scaffold812\|size88987 | 88987 | Z | 1 | 1 | 1 | R | 20.857 | *c00710* | |
| scaffold1046\|size116232 | 116232 | Z | 1 | 1 | 1 | F | 20.946 | | |

**Table S5. Genes in Figure 3**

| Abbreviation | Description | Scaffold |
|---|---|---|
| OR1 | Olfactory receptor 5 | scaffold275 |
| magu | SPARC | scaffold275 |
| DUBAI | Ubiquitin carboxyl | scaffold275 |
| un1 | unknown protein | scaffold532 |
| grnd | Grindelwald | scaffold532 |
| per | period | scaffold532 |
| yCOP | Coatomer | scaffold532 |
| CG6752 | RING SPRY | scaffold459 |
| un3 | unknown protein | scaffold459 |
| sca | scaborous | scaffold459<br>scaffold8883<br>scaffold221 |
| oseg1 | Putative intraflagellar transport 122 (wd-40 domain containing) | scaffold221 |
| *GAPsec* | GTPase activating protein, SECIS-dependent read-through | scaffold221 |
| *whd* | withered; Carnitine O-palmitoyltransferase 1 | scaffold221 |
| Ldh | lactate dehydrogenase-L | scaffold193 |
| trio | Triple functional domain protein | scaffold193 |
| blot | Sodium- and chloride-dependent neutral and basic amino acid transporter B(0+)-like protein | scaffold193<br>scaffold137 |
| csw | corkscrew | scaffold137 |
| CG32809 | Coiled-coil domain-containing protein | scaffold137 |
| meigo | Solute carrier family 35 member B1 homolog | scaffold137<br>scaffold87 |
| Pdfr | Class B secretin-like G-protein coupled receptor | scaffold87 |
| trol | terribly reduced optic lobes; heperan sulfate | scaffold87 |
| CG10467 | aldose-1 epimerase | scaffold50 |
| kon | kontiki; chondroitin sulfate proteoglycan 4 | scaffold50 |

**Table S6. Population covariance from BayPASS**

| **Zchr** | EA | GEN | LA | PY | BV |
|---|---|---|---|---|---|
| EA | 0.009759 | 0.006276 | -0.010536 | 0.007115 | 0.006824 |
| Gen | 0.006276 | 0.019102 | -0.019033 | 0.014849 | 0.014758 |
| LA | -0.010536 | -0.019033 | 0.040306 | -0.019901 | -0.019919 |
| PY | 0.007115 | 0.014849 | -0.019901 | 0.017866 | 0.014944 |
| BV | 0.006824 | 0.014758 | -0.019919 | 0.014944 | 0.017713 |

| **Auto** | EA | GEN | LA | PY | BV |
|---|---|---|---|---|---|
| EA | 0.001049 | -0.000142 | -0.000914 | 0.000112 | -0.000118 |
| Gen | -0.000142 | 0.001397 | 0.001461 | 0.000132 | 0.000111 |
| LA | -0.000914 | 0.001461 | 0.015081 | -0.000702 | -0.000389 |
| PY | 0.000112 | 0.000132 | -0.000702 | 0.000943 | 0.000153 |
| BV | -0.000118 | 0.000111 | -0.000389 | 0.000153 | 0.000916 |

| **All=Auto+Z** | EA | GEN | LA | PY | BV |
|---|---|---|---|---|---|
| EA | 0.0011 | -0.000181 | -0.000723 | 0.000043 | -0.000155 |
| Gen | -0.000181 | 0.001726 | 0.001648 | 0.000343 | 0.000386 |
| LA | -0.000723 | 0.001648 | 0.01661 | -0.000634 | -0.000083 |
| PY | 0.000043 | 0.000343 | -0.000634 | 0.001195 | 0.000381 |
| BV | -0.000155 | 0.000386 | -0.000083 | 0.000381 | 0.001215 |

**Table S7. CMH joint outliers**

| Chr | QTL | Mb | Scaffold | Start | End | Outliers | Gene Annotation | D. mel annotation |
|---|---|---|---|---|---|---|---|---|
| 1 | Q1 | 13.48 | scaffold532\|size371787 | 178601 | 179802 | 1 | Function unknown | un2 |
| 1 | Q1 | 13.55 | scaffold532\|size371787 | 75000 | 149301 | 2 | Period | *per* |
| 1 | Q1 | 13.67 | scaffold459\|size265770 | 43909 | 149893 | 2 | Function unknown | un3 |
| 1 | Q1 | 14.08 | scaffold221\|size590889 | 207992 | 229237 | 1 | GTPase activating protein, SECIS-dependent read-through | *GAPsec* |
| 1 | Q1 | 14.28 | scaffold221\|size590889 | 394081 | 423512 | 1 | withered; Carnitine O-palmitoyltransferase 1 | *whd* |
| 1 | Q2 | 14.88 | scaffold140\|size987430 | 438021 | 559854 | 2 | SNF4/AMP-activated protein kinase gamma subunit | *SNF4gamma* |
| 1 | Q2 | 15.01 | scaffold140\|size987430 | 675900 | 687782 | 2 | Autophagy-related protein 9A-like protein | *Atg9* |
| 1 | Q2 | 17.55 | scaffold137\|size260670 | 11882 | 207815 | 12 | Coiled-coil domain-containing protein | *CG32809* |
| 1 | Q2 | 17.76 | scaffold87\|size517236 | 1 | 6406 | 1 | Solute carrier family 35 member B1 | *meigo* |
| 1 | Q2 | 17.80 | scaffold87\|size517236 | 43074 | 83200 | 3 | *Pdfr* | *Pdfr* |
| 1 | Q2 | 17.94 | scaffold87\|size517236 | 173994 | 347134 | 2 | *trol* | *trol* |
| 13 | NA | 1.11 | scaffold447\|size529996 | 410849 | 455786 | 1 | Flocculation protein FLO11-like protein | *CG16786* |

**Table S8. Linkage disequilibrium outlier pairs**

**Among QTL**

| QTL1 | QTL2 | max LD ($r^2$) | median LD | mean Dist (Mb) | 99.90% | lower 95% CI | upper 95% CI |
|---|---|---|---|---|---|---|---|
| *Per* | *Pdfr* | 0.753 | 0.102 | 4.504 | 0.563 | 0.532 | 0.600 |
| *magu* | *AmpK/SNF4gamm* | 0.715 | 0.129 | 2.332 | 0.582 | 0.542 | 0.604 |
| *magu* | *Pdfr* | 0.696 | 0.268 | 5.508 | 0.585 | 0.564 | 0.597 |
| *CG6752* | *Pdfr* | 0.677 | 0.064 | 4.132 | 0.563 | 0.532 | 0.600 |
| *unknown1* | *CG32809* | 0.661 | 0.041 | 3.871 | 0.559 | 0.536 | 0.594 |
| *Per* | *meigo* | 0.659 | 0.032 | 4.453 | 0.563 | 0.532 | 0.600 |
| *Per* | *CG32809* | 0.651 | 0.049 | 4.243 | 0.563 | 0.532 | 0.600 |
| *Clock* | *Prx6005* | 0.650 | 0.013 | 2.332 | 0.582 | 0.542 | 0.604 |
| *Per* | *trol* | 0.635 | 0.016 | 4.504 | 0.563 | 0.532 | 0.600 |

**Within QTL**

| QTL1 | QTL1 | | | | | | |
|---|---|---|---|---|---|---|---|
| CG10984 | *whd* | 0.681 | 0.036 | 2.244 | 0.582 | 0.542 | 0.604 |
| *Surf1* | *whd* | 0.681 | 0.070 | 2.244 | 0.582 | 0.542 | 0.604 |
| *unknown2* | *Atg9* | 0.616 | 0.033 | 2.446 | 0.582 | 0.542 | 0.604 |

| Table S9. Significant association *per* plink case/control analysis | | | | | | | | | |
|---|---|---|---|---|---|---|---|---|---|
| Gene | SNP | Type | Major Allele | Minor Alllele | P | fdr | LogQ | Mb | Location |
| *per* | scaffold532\|size371787:75166 | SNP | T | A | 1.85E-06 | 0.00766683 | 2.115 | 13.554 | near Ebox (75057-80) |
| *per* | scaffold532\|size371787:76540 | SNP | T | G | 4.41E-07 | 0.00257742 | 2.589 | 13.553 | between Ebox an UTR |
| *per* | scaffold532\|size371787:78628 | SNP | C | G | 4.00E-10 | 1.01E-05 | 4.996 | 13.551 | 5UTR; creates new Ebox CACGTG |
| *per* | scaffold532\|size371787:78716 | SNP | T | G | 8.77E-09 | 0.00011734 | 3.931 | 13.551 | 5UTR |
| *per* | scaffold532\|size371787:78739 | SNP | T | G | 2.89E-09 | 4.91E-05 | 4.309 | 13.551 | 5UTR |
| *per* | scaffold532\|size371787:85637 | INDEL | AGTAG | A | 8.47E-08 | 0.00070328 | 3.153 | 13.544 | 5UTR |
| *per* | scaffold532\|size371787:85723 | INDEL | T | TA | 4.00E-10 | 1.01E-05 | 4.996 | 13.544 | 5UTR |
| *per* | scaffold532\|size371787:85733 | SNP | C | A | 3.66E-07 | 0.00221886 | 2.654 | 13.544 | 5UTR |
| *per* | scaffold532\|size371787:85896 | SNP | T | A | 8.47E-08 | 0.00070328 | 3.153 | 13.543 | 5UTR |
| *per* | scaffold532\|size371787:87095 | INDEL | TCCC | T | 1.14E-06 | 0.00532452 | 2.274 | 13.542 | 5UTR |
| *per* | scaffold532\|size371787:87524 | SNP | T | C | 9.13E-09 | 0.0001211 | 3.917 | 13.542 | 5UTR |
| *per* | scaffold532\|size371787:90815 | SNP | A | T | 2.41E-06 | 0.00932215 | 2.030 | 13.538 | 5UTR |
| *per* | scaffold532\|size371787:90881 | SNP | A | C | 1.48E-08 | 0.00017551 | 3.756 | 13.538 | 5UTR |
| *per* | scaffold532\|size371787:90925 | SNP | GT | G | 1.43E-08 | 0.00017099 | 3.767 | 13.538 | 5UTR |
| *per* | scaffold532\|size371787:93691 | SNP | G | T | 2.09E-12 | 9.39E-08 | 7.027 | 13.536 | 5UTR |
| *per* | scaffold532\|size371787:100347 | SNP | T | A | 7.27E-08 | 0.00062347 | 3.205 | 13.529 | 5UTR |
| *per* | scaffold532\|size371787:100348 | INDEL | ACGCA | GCGCA | 1.06E-06 | 0.00504083 | 2.297 | 13.529 | 5UTR |
| *per* | scaffold532\|size371787:100534 | SNP | C | T | 8.58E-07 | 0.00425681 | 2.371 | 13.529 | 5UTR |
| *per* | scaffold532\|size371787:100736 | SNP | A | G | 2.42E-06 | 0.0093414 | 2.030 | 13.529 | 5UTR |
| *per* | scaffold532\|size371787:102168 | SNP | A | T | 2.33E-06 | 0.00911106 | 2.040 | 13.527 | Intron |
| *per* | scaffold532\|size371787:102446 | INDEL | AGTAT | A | 9.75E-08 | 0.00078352 | 3.106 | 13.527 | Intron |
| *per* | scaffold532\|size371787:105541 | INDEL | G | GTTTAC | 2.06E-06 | 0.00830769 | 2.081 | 13.524 | Intron |
| *per* | scaffold532\|size371787:112015 | SNP | A | G | 8.47E-08 | 0.00070328 | 3.153 | 13.517 | Intron |
| *per* | scaffold532\|size371787:116704 | INDEL | CAT | C | 4.96E-09 | 7.46E-05 | 4.127 | 13.513 | Intron |
| *per* | scaffold532\|size371787:117528 | INDEL | 7.84 kb deletion | T | 2.16E-08 | 0.0002396 | 3.621 | 13.512 | Intron |
| *per* | scaffold532\|size371787:120819 | SNP | T | C | 6.35E-07 | 0.00340875 | 2.467 | 13.508 | Intron |
| *per* | scaffold532\|size371787:121279 | SNP | A | G | 2.46E-08 | 0.00026493 | 3.577 | 13.508 | Intron |
| *per* | scaffold532\|size371787:121294 | SNP | A | G | 1.11E-07 | 0.00087107 | 3.060 | 13.508 | Intron |
| *per* | scaffold532\|size371787:121300 | INDEL | TTA | T | 2.46E-08 | 0.00026493 | 3.577 | 13.508 | Intron |
| *per* | scaffold532\|size371787:121313 | SNP | A | C | 5.28E-09 | 7.82E-05 | 4.107 | 13.508 | Intron |
| *per* | scaffold532\|size371787:121329 | SNP | A | G | 6.69E-08 | 0.00058368 | 3.234 | 13.508 | Intron |
| *per* | scaffold532\|size371787:121334 | SNP | T | G | 1.18E-07 | 0.00091291 | 3.040 | 13.508 | Intron |
| *per* | scaffold532\|size371787:126158 | SNP | T | C | 7.56E-08 | 0.00064326 | 3.192 | 13.503 | Intron |
| *per* | scaffold532\|size371787:126317 | SNP | T | G | 1.25E-07 | 0.00095426 | 3.020 | 13.503 | Intron |
| *per* | scaffold532\|size371787:128172 | SNP | C | A | 7.53E-09 | 0.00010391 | 3.983 | 13.501 | Intron |
| *per* | scaffold532\|size371787:131364 | INDEL | TA | T | 4.00E-10 | 1.01E-05 | 4.996 | 13.498 | Intron |
| *per* | scaffold532\|size371787:131365 | INDEL | TC | T | 1.28E-08 | 0.00015702 | 3.804 | 13.498 | Intron |
| *per* | scaffold532\|size371787:131463 | SNP | G | T | 2.32E-09 | 4.12E-05 | 4.385 | 13.498 | Intron |
| *per* | scaffold532\|size371787:131844 | SNP | C | G | 5.57E-09 | 8.17E-05 | 4.088 | 13.497 | Intron |
| *per* | scaffold532\|size371787:132085 | SNP | T | G | 3.58E-10 | 9.23E-06 | 5.035 | 13.497 | Intron |
| *per* | scaffold532\|size371787:133557 | INDEL | A | AACAGAGG | 2.27E-06 | 0.0089388 | 2.049 | 13.496 | Intron |
| *per* | scaffold532\|size371787:133565 | INDEL | T | TGGG | 2.27E-06 | 0.0089388 | 2.049 | 13.496 | Intron |
| *per* | scaffold532\|size371787:133580 | SNP | A | G | 2.27E-06 | 0.0089388 | 2.049 | 13.496 | Intron |
| *per* | scaffold532\|size371787:133600 | INDEL | A | AGT | 1.16E-06 | 0.0053824 | 2.269 | 13.496 | Intron |
| *per* | scaffold532\|size371787:133829 | SNP | T | C | 6.33E-07 | 0.00340168 | 2.468 | 13.495 | Syn |
| *per* | scaffold532\|size371787:133844 | SNP | A | C | 4.39E-07 | 0.00256856 | 2.590 | 13.495 | Syn |
| *per* | scaffold532\|size371787:133907 | SNP | A | C | 4.12E-08 | 0.00039529 | 3.403 | 13.495 | Syn |
| *per* | scaffold532\|size371787:133971 | SNP | G | A | 1.49E-07 | 0.00110393 | 2.957 | 13.495 | Intron |
| *per* | scaffold532\|size371787:133989 | SNP | C | T | 1.40E-07 | 0.00104571 | 2.981 | 13.495 | Intron |
| *per* | scaffold532\|size371787:134048 | SNP | G | T | 1.14E-06 | 0.00533903 | 2.273 | 13.495 | Intron |
| *per* | scaffold532\|size371787:134055 | SNP | A | G | 1.03E-06 | 0.00492416 | 2.308 | 13.495 | Intron |
| *per* | scaffold532\|size371787:134527 | SNP | C | T | 1.76E-07 | 0.00126117 | 2.899 | 13.495 | Intron |
| *per* | scaffold532\|size371787:134880 | SNP | A | G | 3.31E-10 | 8.67E-06 | 5.062 | 13.494 | Intron |
| *per* | scaffold532\|size371787:134885 | SNP | A | T | 3.31E-10 | 8.67E-06 | 5.062 | 13.494 | Intron |
| *per* | scaffold532\|size371787:135799 | SNP | A | G | 8.80E-08 | 0.00072441 | 3.140 | 13.493 | Intron |
| *per* | scaffold532\|size371787:135951 | SNP | T | A | 4.21E-10 | 1.05E-05 | 4.979 | 13.493 | Syn |
| *per* | scaffold532\|size371787:135984 | SNP | A | G | 9.12E-09 | 0.00012099 | 3.917 | 13.493 | Syn |
| *per* | scaffold532\|size371787:136259 | INDEL | C | CTTTTATT | 7.79E-07 | 0.00396637 | 2.402 | 13.493 | Intron |
| *per* | scaffold532\|size371787:136345 | SNP | T | C | 2.56E-07 | 0.0016898 | 2.772 | 13.493 | Intron |
| *per* | scaffold532\|size371787:136554 | INDEL | T | TCTACCGTG | 2.16E-08 | 0.0002396 | 3.621 | 13.493 | Intron |
| *per* | scaffold532\|size371787:137090 | SNP | G | A | 7.24E-08 | 0.00062146 | 3.207 | 13.492 | Intron |
| *per* | scaffold532\|size371787:137689 | SNP | C | T | 4.05E-08 | 0.00038991 | 3.409 | 13.492 | Intron |
| *per* | scaffold532\|size371787:138138 | INDEL | ATTTAT | ATTAT | 2.45E-06 | 0.00943882 | 2.025 | 13.491 | Intron |
| *per* | scaffold532\|size371787:138308 | INDEL | AT | A | 9.64E-08 | 0.00077691 | 3.110 | 13.491 | Intron |
| *per* | scaffold532\|size371787:138319 | SNP | T | A | 9.13E-09 | 0.0001211 | 3.917 | 13.491 | Intron |
| *per* | scaffold532\|size371787:140359 | SNP | A | G | 1.33E-06 | 0.00598927 | 2.223 | 13.489 | Syn |
| *per* | scaffold532\|size371787:140360 | SNP | C | A | 1.33E-06 | 0.00598927 | 2.223 | 13.489 | Nonsyn |
| *per* | scaffold532\|size371787:140369 | SNP | C | T | 1.33E-06 | 0.00598927 | 2.223 | 13.489 | Nonsyn |
| *per* | scaffold532\|size371787:140498 | SNP | A | C | 1.03E-07 | 0.00081733 | 3.088 | 13.489 | Nonsyn |
| *per* | scaffold532\|size371787:140791 | SNP | G | A | 1.03E-07 | 0.00081733 | 3.088 | 13.489 | Intron |
| *per* | scaffold532\|size371787:142016 | INDEL | T | TTTG | 8.47E-08 | 0.00070328 | 3.153 | 13.487 | Intron |
| *per* | scaffold532\|size371787:142027 | SNP | T | G | 8.47E-08 | 0.00070328 | 3.153 | 13.487 | Intron |
| *per* | scaffold532\|size371787:142062 | SNP | A | G | 6.32E-07 | 0.00339579 | 2.469 | 13.487 | Intron |
| *per* | scaffold532\|size371787:142445 | INDEL | T | TAG | 1.87E-06 | 0.00772993 | 2.112 | 13.487 | Intron |
| *per* | scaffold532\|size371787:142448 | INDEL | GTC | G | 3.66E-07 | 0.00221886 | 2.654 | 13.487 | Intron |
| *per* | scaffold532\|size371787:142456 | SNP | T | C | 3.66E-07 | 0.00221886 | 2.654 | 13.487 | Intron |
| *per* | scaffold532\|size371787:142677 | SNP | T | C | 2.67E-08 | 0.0002833 | 3.548 | 13.487 | Syn |
| *per* | scaffold532\|size371787:143855 | SNP | A | G | 2.20E-08 | 0.00024347 | 3.614 | 13.485 | 3UTR |

**Table S10. Significant association *Pdfr* plink case/control analysis**

| Gene | SNP | Type | Major Allele | Minor Alllele | P | fdr | LogQ | Mb | Location |
|---|---|---|---|---|---|---|---|---|---|
| *Pdfr* | scaffold87\|size517236:45716 | SNP | C | T | 3.82E-12 | 1.62E-07 | 6.790 | 17.807 | 3UTR |
| *Pdfr* | scaffold87\|size517236:45997 | INDEL | A | ATTTT | 1.52E-17 | 5.52E-12 | 11.258 | 17.807 | 3UTR |
| *Pdfr* | scaffold87\|size517236:46103 | SNP | C | T | 6.03E-15 | 6.03E-10 | 9.220 | 17.807 | 3UTR |
| *Pdfr* | scaffold87\|size517236:46147 | INDEL | AT | A | 2.66E-06 | 0.00999353 | 2.000 | 17.807 | 3UTR |
| *Pdfr* | scaffold87\|size517236:46148 | SNP | T | G | 2.66E-06 | 0.00999353 | 2.000 | 17.807 | 3UTR |
| *Pdfr* | scaffold87\|size517236:46150 | SNP | C | G | 2.66E-06 | 0.00999353 | 2.000 | 17.807 | 3UTR |
| *Pdfr* | scaffold87\|size517236:46152 | INDEL | TACAATTTGTATCTC | T | 2.66E-06 | 0.00999353 | 2.000 | 17.807 | 3UTR |
| *Pdfr* | scaffold87\|size517236:46176 | INDEL | CAAATA | C | 5.87E-15 | 5.91E-10 | 9.228 | 17.807 | 3UTR |
| *Pdfr* | scaffold87\|size517236:46197 | SNP | T | G | 5.87E-15 | 5.91E-10 | 9.228 | 17.807 | 3UTR |
| *Pdfr* | scaffold87\|size517236:46459 | SNP | A | C | 9.56E-13 | 4.59E-08 | 7.338 | 17.808 | Intron |
| *Pdfr* | scaffold87\|size517236:46471 | SNP | C | T | 9.56E-13 | 4.59E-08 | 7.338 | 17.808 | Intron |
| *Pdfr* | scaffold87\|size517236:46486 | SNP | T | C | 6.94E-12 | 2.80E-07 | 6.553 | 17.808 | Intron |
| *Pdfr* | scaffold87\|size517236:46491 | SNP | A | G | 1.99E-13 | 1.11E-08 | 7.955 | 17.808 | Intron |
| *Pdfr* | scaffold87\|size517236:46513 | INDEL | T | TA | 9.30E-15 | 8.58E-10 | 9.067 | 17.808 | Intron |
| *Pdfr* | scaffold87\|size517236:46543 | SNP | T | C | 9.30E-15 | 8.58E-10 | 9.067 | 17.808 | Intron |
| *Pdfr* | scaffold87\|size517236:46564 | SNP | T | C | 2.98E-14 | 2.27E-09 | 8.644 | 17.808 | Intron |
| *Pdfr* | scaffold87\|size517236:46575 | INDEL | G | GC | 2.16E-13 | 1.19E-08 | 7.924 | 17.808 | Intron |
| *Pdfr* | scaffold87\|size517236:46671 | SNP | C | A | 9.56E-13 | 4.59E-08 | 7.338 | 17.808 | Intron |
| *Pdfr* | scaffold87\|size517236:46753 | INDEL | ACG | A | 2.39E-08 | 0.00025947 | 3.586 | 17.808 | Intron |
| *Pdfr* | scaffold87\|size517236:46754 | SNP | T | A | 2.39E-08 | 0.00025947 | 3.586 | 17.808 | Intron |
| *Pdfr* | scaffold87\|size517236:46818 | INDEL | A | AACCTTATGCTGTTTTTTTAAATTGTTTTGACTTGAAGTC | 8.37E-07 | 0.00417614 | 2.379 | 17.808 | Intron |
| *Pdfr* | scaffold87\|size517236:47009 | SNP | G | T | 6.32E-07 | 0.00339579 | 2.469 | 17.808 | Intron |
| *Pdfr* | scaffold87\|size517236:47031 | SNP | G | A | 7.56E-08 | 0.00064326 | 3.192 | 17.808 | Intron |
| *Pdfr* | scaffold87\|size517236:47044 | SNP | G | C | 9.54E-09 | 0.00012528 | 3.902 | 17.808 | Intron |
| *Pdfr* | scaffold87\|size517236:47051 | SNP | A | G | 5.95E-09 | 8.59E-05 | 4.066 | 17.808 | Intron |
| *Pdfr* | scaffold87\|size517236:47070 | SNP | T | C | 2.16E-08 | 0.0002396 | 3.621 | 17.808 | Syn |
| *Pdfr* | scaffold87\|size517236:47142 | SNP | C | G | 4.39E-07 | 0.00256856 | 2.590 | 17.808 | Syn |
| *Pdfr* | scaffold87\|size517236:47391 | SNP | A | T | 2.83E-13 | 1.53E-08 | 7.815 | 17.809 | Intron |
| *Pdfr* | scaffold87\|size517236:48815 | SNP | A | G | 1.97E-15 | 2.52E-10 | 9.599 | 17.810 | Intron |
| *Pdfr* | scaffold87\|size517236:53709 | SNP | A | T | 7.16E-16 | 1.08E-10 | 9.967 | 17.815 | Intron |
| *Pdfr* | scaffold87\|size517236:56411 | INDEL | 825 bp deletion | T | 1.01E-13 | 6.47E-09 | 8.189 | 17.818 | Intron |
| *Pdfr* | scaffold87\|size517236:57259 | INDEL | A | ATTAGTTATCTTCTGATATTTTTCTTCTTCT | 4.71E-11 | 1.59E-06 | 5.799 | 17.819 | Intron |
| *Pdfr* | scaffold87\|size517236:57397 | SNP | A | G | 1.99E-15 | 2.54E-10 | 9.595 | 17.819 | Intron |
| *Pdfr* | scaffold87\|size517236:57525 | INDEL | GTACGGCAATACTATC | G | 1.50E-15 | 2.02E-10 | 9.695 | 17.819 | Intron |
| *Pdfr* | scaffold87\|size517236:57656 | SNP | T | G | 6.01E-15 | 6.02E-10 | 9.220 | 17.819 | Intron |
| *Pdfr* | scaffold87\|size517236:57953 | SNP | A | G | 5.92E-14 | 4.08E-09 | 8.389 | 17.819 | Intron |
| *Pdfr* | scaffold87\|size517236:58210 | INDEL | TGCAAAAA | T | 1.76E-10 | 5.13E-06 | 5.290 | 17.819 | Intron |
| *Pdfr* | scaffold87\|size517236:58212 | INDEL | TTAAA | T | 5.62E-10 | 1.32E-05 | 4.879 | 17.819 | Intron |
| *Pdfr* | scaffold87\|size517236:58605 | SNP | T | G | 4.29E-12 | 1.79E-07 | 6.747 | 17.820 | Intron |
| *Pdfr* | scaffold87\|size517236:58815 | INDEL | TT | AT | 1.53E-12 | 7.05E-08 | 7.152 | 17.820 | Intron |
| *Pdfr* | scaffold87\|size517236:58826 | INDEL | GA | GAA | 1.83E-08 | 0.00020992 | 3.678 | 17.820 | Intron |
| *Pdfr* | scaffold87\|size517236:59443 | SNP | C | T | 1.43E-12 | 6.64E-08 | 7.178 | 17.821 | Intron |
| *Pdfr* | scaffold87\|size517236:59455 | SNP | C | G | 8.06E-10 | 1.78E-05 | 4.750 | 17.821 | Intron |
| *Pdfr* | scaffold87\|size517236:59643 | INDEL | GTT | GT | 7.54E-07 | 0.00387261 | 2.412 | 17.821 | Intron |
| *Pdfr* | scaffold87\|size517236:59960 | SNP | C | G | 1.40E-13 | 8.45E-09 | 8.073 | 17.821 | Intron |
| *Pdfr* | scaffold87\|size517236:60203 | INDEL | CA | C | 1.32E-14 | 1.17E-09 | 8.932 | 17.821 | Intron |
| *Pdfr* | scaffold87\|size517236:60907 | SNP | C | G | 5.82E-14 | 4.02E-09 | 8.396 | 17.822 | Intron |
| *Pdfr* | scaffold87\|size517236:61083 | SNP | A | T | 1.40E-13 | 8.45E-09 | 8.073 | 17.822 | Intron |
| *Pdfr* | scaffold87\|size517236:61267 | SNP | A | T | 7.33E-15 | 6.96E-10 | 9.157 | 17.823 | Intron |
| *Pdfr* | scaffold87\|size517236:61967 | SNP | T | A | 3.53E-14 | 2.57E-09 | 8.590 | 17.823 | Intron |
| *Pdfr* | scaffold87\|size517236:62217 | SNP | T | A | 7.33E-15 | 6.96E-10 | 9.157 | 17.823 | Intron |
| *Pdfr* | scaffold87\|size517236:62561 | SNP | C | T | 3.68E-12 | 1.57E-07 | 6.804 | 17.824 | Intron |
| *Pdfr* | scaffold87\|size517236:62776 | INDEL | T | 500 bp insertion | 3.79E-13 | 2.01E-08 | 7.697 | 17.824 | Intron |
| *Pdfr* | scaffold87\|size517236:63392 | SNP | G | T | 4.39E-10 | 1.08E-05 | 4.967 | 17.825 | Intron |
| *Pdfr* | scaffold87\|size517236:63393 | SNP | T | A | 4.39E-10 | 1.08E-05 | 4.967 | 17.825 | Intron |
| *Pdfr* | scaffold87\|size517236:65111 | SNP | A | AT | 9.57E-16 | 1.38E-10 | 9.860 | 17.826 | Intron |
| *Pdfr* | scaffold87\|size517236:65913 | SNP | A | T | 2.64E-13 | 1.43E-08 | 7.845 | 17.827 | Intron |
| *Pdfr* | scaffold87\|size517236:66396 | SNP | G | A | 1.40E-13 | 8.45E-09 | 8.073 | 17.828 | NonSyn |
| *Pdfr* | scaffold87\|size517236:66493 | SNP | A | G | 1.74E-15 | 2.28E-10 | 9.642 | 17.828 | Intron |
| *Pdfr* | scaffold87\|size517236:66516 | INDEL | CTTA | C | 7.12E-16 | 1.07E-10 | 9.971 | 17.828 | Intron |
| *Pdfr* | scaffold87\|size517236:66529 | INDEL | CAAGAGAATA | C | 2.93E-16 | 5.67E-11 | 10.246 | 17.828 | Intron |
| *Pdfr* | scaffold87\|size517236:66541 | SNP | C | T | 1.74E-15 | 2.28E-10 | 9.642 | 17.828 | Intron |
| *Pdfr* | scaffold87\|size517236:66562 | SNP | A | G | 4.74E-17 | 1.34E-11 | 10.873 | 17.828 | Intron |
| *Pdfr* | scaffold87\|size517236:66567 | SNP | G | A | 1.63E-18 | 1.31E-12 | 11.883 | 17.828 | Intron |
| *Pdfr* | scaffold87\|size517236:66571 | INDEL | CT | C | 4.74E-17 | 1.34E-11 | 10.873 | 17.828 | Intron |
| *Pdfr* | scaffold87\|size517236:66572 | SNP | A | G | 4.74E-17 | 1.34E-11 | 10.873 | 17.828 | Intron |
| *Pdfr* | scaffold87\|size517236:66603 | SNP | T | C | 1.80E-13 | 1.03E-08 | 7.987 | 17.828 | Intron |
| *Pdfr* | scaffold87\|size517236:66612 | SNP | T | A | 9.71E-13 | 4.66E-08 | 7.332 | 17.828 | Intron |
| *Pdfr* | scaffold87\|size517236:66635 | SNP | C | G | 9.71E-13 | 4.66E-08 | 7.332 | 17.828 | Intron |
| *Pdfr* | scaffold87\|size517236:66646 | INDEL | AACT | A | 1.94E-12 | 8.77E-08 | 7.057 | 17.828 | Intron |
| *Pdfr* | scaffold87\|size517236:66672 | SNP | C | A | 9.71E-13 | 4.66E-08 | 7.332 | 17.828 | Intron |
| *Pdfr* | scaffold87\|size517236:66673 | SNP | T | TA | 6.87E-11 | 2.24E-06 | 5.650 | 17.828 | Intron |
| *Pdfr* | scaffold87\|size517236:66677 | SNP | G | A | 9.71E-13 | 4.66E-08 | 7.332 | 17.828 | Intron |
| *Pdfr* | scaffold87\|size517236:66688 | SNP | T | A | 9.71E-13 | 4.66E-08 | 7.332 | 17.828 | Intron |
| *Pdfr* | scaffold87\|size517236:66689 | SNP | T | A | 9.93E-12 | 3.88E-07 | 6.411 | 17.828 | Intron |
| *Pdfr* | scaffold87\|size517236:66690 | SNP | T | C | 9.71E-13 | 4.66E-08 | 7.332 | 17.828 | Intron |
| *Pdfr* | scaffold87\|size517236:66695 | INDEL | AATAATTTAAAATGAAATAT | A | 9.71E-13 | 4.66E-08 | 7.332 | 17.828 | Intron |
| *Pdfr* | scaffold87\|size517236:66696 | INDEL | CT | C | 9.71E-13 | 4.66E-08 | 7.332 | 17.828 | Intron |

| Gene | SNP | Type | Major Allele | Minor Alllele | P | fdr | LogQ | Mb | Location |
|---|---|---|---|---|---|---|---|---|---|
| *Pdfr* | scaffold87\|size517236:66709 | SNP | C | T | 1.80E-13 | 1.03E-08 | 7.987 | 17.828 | Intron |
| *Pdfr* | scaffold87\|size517236:66745 | INDEL | GT | G | 7.12E-16 | 1.07E-10 | 9.971 | 17.828 | Intron |
| *Pdfr* | scaffold87\|size517236:66762 | SNP | T | G | 7.12E-16 | 1.07E-10 | 9.971 | 17.828 | Intron |
| *Pdfr* | scaffold87\|size517236:66819 | SNP | A | G | 1.98E-14 | 1.64E-09 | 8.785 | 17.828 | Intron |
| *Pdfr* | scaffold87\|size517236:66845 | SNP | G | A | 6.03E-16 | 9.63E-11 | 10.016 | 17.828 | Intron |
| *Pdfr* | scaffold87\|size517236:66866 | SNP | C | G | 1.98E-14 | 1.64E-09 | 8.785 | 17.828 | Intron |
| *Pdfr* | scaffold87\|size517236:66868 | INDEL | G | GT | 1.98E-14 | 1.64E-09 | 8.785 | 17.828 | Intron |
| *Pdfr* | scaffold87\|size517236:66871 | SNP | A | T | 1.98E-14 | 1.64E-09 | 8.785 | 17.828 | Intron |
| *Pdfr* | scaffold87\|size517236:66878 | SNP | C | A | 1.98E-14 | 1.64E-09 | 8.785 | 17.828 | Intron |
| *Pdfr* | scaffold87\|size517236:66894 | SNP | G | C | 4.39E-14 | 3.12E-09 | 8.506 | 17.828 | Intron |
| *Pdfr* | scaffold87\|size517236:66914 | SNP | T | A | 1.74E-15 | 2.28E-10 | 9.642 | 17.828 | Intron |
| *Pdfr* | scaffold87\|size517236:66928 | SNP | A | T | 1.74E-15 | 2.28E-10 | 9.642 | 17.828 | Intron |
| *Pdfr* | scaffold87\|size517236:66931 | SNP | T | G | 1.74E-15 | 2.28E-10 | 9.642 | 17.828 | Intron |
| *Pdfr* | scaffold87\|size517236:66960 | SNP | G | A | 3.53E-14 | 2.57E-09 | 8.590 | 17.828 | Intron |
| *Pdfr* | scaffold87\|size517236:66989 | SNP | T | A | 9.71E-13 | 4.66E-08 | 7.332 | 17.828 | Intron |
| *Pdfr* | scaffold87\|size517236:66999 | SNP | C | T | 1.04E-13 | 6.63E-09 | 8.178 | 17.828 | Intron |
| *Pdfr* | scaffold87\|size517236:67007 | SNP | C | G | 9.57E-12 | 3.75E-07 | 6.426 | 17.828 | Intron |
| *Pdfr* | scaffold87\|size517236:77518 | INDEL | T | TTAA | 2.04E-11 | 7.53E-07 | 6.123 | 17.839 | 5UTR |
| *Pdfr* | scaffold87\|size517236:77555 | INDEL | T | TTAGG | 4.88E-13 | 2.53E-08 | 7.597 | 17.839 | 5UTR |
| *Pdfr* | scaffold87\|size517236:77593 | SNP | G | T | 1.01E-13 | 6.47E-09 | 8.189 | 17.839 | 5UTR |
| *Pdfr* | scaffold87\|size517236:77595 | SNP | T | A | 1.01E-13 | 6.47E-09 | 8.189 | 17.839 | 5UTR |
| *Pdfr* | scaffold87\|size517236:77639 | SNP | C | A | 1.01E-13 | 6.47E-09 | 8.189 | 17.839 | 5UTR |
| *Pdfr* | scaffold87\|size517236:77642 | SNP | A | T | 1.74E-14 | 1.47E-09 | 8.833 | 17.839 | 5UTR |
| *Pdfr* | scaffold87\|size517236:77672 | SNP | A | G | 9.96E-16 | 1.42E-10 | 9.848 | 17.839 | 5UTR |
| *Pdfr* | scaffold87\|size517236:77680 | SNP | A | T | 6.03E-15 | 6.03E-10 | 9.220 | 17.839 | 5UTR |
| *Pdfr* | scaffold87\|size517236:77689 | SNP | G | A | 6.03E-15 | 6.03E-10 | 9.220 | 17.839 | 5UTR |
| *Pdfr* | scaffold87\|size517236:77708 | SNP | G | A | 3.07E-14 | 2.32E-09 | 8.635 | 17.839 | 5UTR |
| *Pdfr* | scaffold87\|size517236:77719 | SNP | A | G | 2.29E-15 | 2.83E-10 | 9.548 | 17.839 | 5UTR |
| *Pdfr* | scaffold87\|size517236:77800 | SNP | A | G | 1.66E-13 | 9.68E-09 | 8.014 | 17.839 | 5UTR |
| *Pdfr* | scaffold87\|size517236:77847 | SNP | G | A | 5.16E-15 | 5.35E-10 | 9.272 | 17.839 | 5UTR |
| *Pdfr* | scaffold87\|size517236:77854 | INDEL | TGTA | CGTA | 4.95E-14 | 3.48E-09 | 8.458 | 17.839 | 5UTR |
| *Pdfr* | scaffold87\|size517236:77866 | INDEL | TAA | T | 5.16E-15 | 5.35E-10 | 9.272 | 17.839 | 5UTR |
| *Pdfr* | scaffold87\|size517236:77875 | SNP | A | T | 3.07E-14 | 2.32E-09 | 8.635 | 17.839 | 5UTR |
| *Pdfr* | scaffold87\|size517236:77890 | INDEL | C | CA | 5.16E-15 | 5.35E-10 | 9.272 | 17.839 | 5UTR |
| *Pdfr* | scaffold87\|size517236:77918 | INDEL | TAC | TAAC | 3.64E-15 | 4.05E-10 | 9.393 | 17.839 | 5UTR |
| *Pdfr* | scaffold87\|size517236:77920 | SNP | T | A | 3.64E-15 | 4.05E-10 | 9.393 | 17.839 | 5UTR |
| *Pdfr* | scaffold87\|size517236:78030 | INDEL | G | GTC | 1.43E-16 | 3.21E-11 | 10.493 | 17.839 | 5UTR |
| *Pdfr* | scaffold87\|size517236:78126 | SNP | T | C | 3.64E-15 | 4.05E-10 | 9.393 | 17.839 | 5UTR |
| *Pdfr* | scaffold87\|size517236:78146 | SNP | G | C | 7.11E-13 | 3.55E-08 | 7.450 | 17.839 | 5UTR |
| *Pdfr* | scaffold87\|size517236:78168 | SNP | A | G | 2.25E-14 | 1.82E-09 | 8.740 | 17.839 | 5UTR |
| *Pdfr* | scaffold87\|size517236:78194 | SNP | A | T | 1.99E-15 | 2.54E-10 | 9.595 | 17.839 | 5UTR; near putative Ebox CACGTG |
| *Pdfr* | scaffold87\|size517236:78197 | INDEL | T | TA | 1.43E-16 | 3.21E-11 | 10.493 | 17.839 | 5UTR; near putative Ebox CACGTG |
| *Pdfr* | scaffold87\|size517236:78201 | SNP | C | A | 1.43E-16 | 3.21E-11 | 10.493 | 17.839 | 5UTR; near putative Ebox CACGTG |
| *Pdfr* | scaffold87\|size517236:78215 | SNP | G | T | 1.43E-16 | 3.21E-11 | 10.493 | 17.839 | 5UTR |
| *Pdfr* | scaffold87\|size517236:78272 | INDEL | A | ACTAATACTGTG | 3.64E-15 | 4.05E-10 | 9.393 | 17.840 | 5UTR |
| *Pdfr* | scaffold87\|size517236:78480 | SNP | A | T | 2.29E-15 | 2.83E-10 | 9.548 | 17.840 | 5UTR |
| *Pdfr* | scaffold87\|size517236:78510 | SNP | A | C | 6.03E-15 | 6.03E-10 | 9.220 | 17.840 | 5UTR |
| *Pdfr* | scaffold87\|size517236:78529 | SNP | G | A | 2.29E-15 | 2.83E-10 | 9.548 | 17.840 | 5UTR |
| *Pdfr* | scaffold87\|size517236:78559 | SNP | A | T | 3.84E-14 | 2.78E-09 | 8.556 | 17.840 | 5UTR |
| *Pdfr* | scaffold87\|size517236:78572 | INDEL | ATAATAATATTAT | A | 1.77E-14 | 1.50E-09 | 8.824 | 17.840 | 5UTR |
| *Pdfr* | scaffold87\|size517236:78639 | SNP | C | T | 2.58E-16 | 5.11E-11 | 10.292 | 17.840 | 5UTR |
| *Pdfr* | scaffold87\|size517236:78705 | SNP | A | T | 1.32E-14 | 1.17E-09 | 8.932 | 17.840 | 5UTR |
| *Pdfr* | scaffold87\|size517236:78750 | SNP | G | T | 2.66E-11 | 9.58E-07 | 6.019 | 17.840 | 5UTR |
| *Pdfr* | scaffold87\|size517236:78762 | SNP | T | C | 2.66E-11 | 9.58E-07 | 6.019 | 17.840 | 5UTR |
| *Pdfr* | scaffold87\|size517236:79153 | INDEL | CACTG | C | 2.40E-10 | 6.66E-06 | 5.177 | 17.840 | 5UTR |
| *Pdfr* | scaffold87\|size517236:79218 | INDEL | GTACCTTA | G | 1.02E-08 | 0.00013196 | 3.880 | 17.840 | 5UTR |
| *Pdfr* | scaffold87\|size517236:79393 | INDEL | A | AAAGATGATTACCTAAGACACAA ATTATTAATTAAT | 1.77E-09 | 3.32E-05 | 4.479 | 17.841 | 5UTR |
| *Pdfr* | scaffold87\|size517236:79457 | SNP | GT | G | 1.14E-13 | 7.18E-09 | 8.144 | 17.841 | 5UTR |
| *Pdfr* | scaffold87\|size517236:79470 | INDEL | T | TTTC | 1.14E-13 | 7.18E-09 | 8.144 | 17.841 | 5UTR |
| *Pdfr* | scaffold87\|size517236:79476 | INDEL | A | AAGGTACCCGTGCTGCCCGGGCT TCCCC | 1.14E-13 | 7.18E-09 | 8.144 | 17.841 | 5UTR |
| *Pdfr* | scaffold87\|size517236:79674 | SNP | TT | AT | 4.74E-17 | 1.34E-11 | 10.873 | 17.841 | 5UTR |
| *Pdfr* | scaffold87\|size517236:79759 | SNP | T | G | 3.72E-15 | 4.12E-10 | 9.385 | 17.841 | 5UTR |
| *Pdfr* | scaffold87\|size517236:79774 | SNP | A | G | 5.92E-14 | 4.08E-09 | 8.389 | 17.841 | 5UTR |
| *Pdfr* | scaffold87\|size517236:79955 | INDEL | AAGAGT | AAGT | 7.12E-16 | 1.07E-10 | 9.971 | 17.841 | 5UTR |
| *Pdfr* | scaffold87\|size517236:80066 | SNP | G | A | 7.33E-15 | 6.96E-10 | 9.157 | 17.841 | 5UTR |
| *Pdfr* | scaffold87\|size517236:80226 | INDEL | AT | A | 1.40E-13 | 8.45E-09 | 8.073 | 17.841 | 5UTR |
| *Pdfr* | scaffold87\|size517236:80256 | SNP | G | T | 4.74E-17 | 1.34E-11 | 10.873 | 17.842 | 5UTR |
| *Pdfr* | scaffold87\|size517236:80277 | SNP | T | A | 4.74E-17 | 1.34E-11 | 10.873 | 17.842 | 5UTR |
| *Pdfr* | scaffold87\|size517236:80439 | INDEL | 1460 bp deletion | G | 7.33E-15 | 6.96E-10 | 9.157 | 17.842 | 5UTR |
| *Pdfr* | scaffold87\|size517236:81909 | SNP | A | T | 2.52E-14 | 2.00E-09 | 8.699 | 17.843 | 5UTR |
| *Pdfr* | scaffold87\|size517236:82193 | INDEL | ATTTTG | A | 1.50E-15 | 2.02E-10 | 9.695 | 17.843 | 5UTR |
| *Pdfr* | scaffold87\|size517236:82196 | INDEL | G | GGC | 1.50E-15 | 2.02E-10 | 9.695 | 17.843 | 5UTR |
| *Pdfr* | scaffold87\|size517236:82388 | SNP | T | C | 3.53E-14 | 2.57E-09 | 8.590 | 17.844 | 5UTR |
| *Pdfr* | scaffold87\|size517236:82400 | SNP | T | G | 2.58E-16 | 5.11E-11 | 10.292 | 17.844 | 5UTR |
| *Pdfr* | scaffold87\|size517236:82406 | INDEL | GTACC | G | 1.12E-17 | 4.73E-12 | 11.325 | 17.844 | 5UTR |
| *Pdfr* | scaffold87\|size517236:82464 | SNP | C | T | 7.33E-15 | 6.96E-10 | 9.157 | 17.844 | 5UTR |
| *Pdfr* | scaffold87\|size517236:82693 | SNP | G | T | 7.33E-15 | 6.96E-10 | 9.157 | 17.844 | 5UTR |
| *Pdfr* | scaffold87\|size517236:82731 | SNP | C | A | 7.12E-16 | 1.07E-10 | 9.971 | 17.844 | 5UTR |
| *Pdfr* | scaffold87\|size517236:82806 | SNP | A | G | 7.33E-15 | 6.96E-10 | 9.157 | 17.844 | 5UTR |

| Gene | SNP | Type | Major Allele | Minor Alllele | P | fdr | LogQ | Mb | Location |
|---|---|---|---|---|---|---|---|---|---|
| *Pdfr* | scaffold87\|size517236:82833 | SNP | C | A | 7.33E-15 | 6.96E-10 | 9.157 | 17.844 | 5UTR |
| *Pdfr* | scaffold87\|size517236:82839 | SNP | A | C | 7.33E-15 | 6.96E-10 | 9.157 | 17.844 | 5UTR |
| *Pdfr* | scaffold87\|size517236:82890 | SNP | G | A | 1.79E-14 | 1.51E-09 | 8.821 | 17.844 | 5UTR |
| *Pdfr* | scaffold87\|size517236:82992 | INDEL | C | CAT | 6.94E-12 | 2.80E-07 | 6.553 | 17.844 | 5UTR |
| *Pdfr* | scaffold87\|size517236:83041 | INDEL | CTG | C | 1.27E-06 | 0.00579491 | 2.237 | 17.844 | 5UTR |
| *Pdfr* | scaffold87\|size517236:83063 | SNP | G | T | 3.93E-11 | 1.35E-06 | 5.870 | 17.844 | 5UTR |
| *Pdfr* | scaffold87\|size517236:83087 | SNP | G | A | 8.21E-14 | 5.43E-09 | 8.265 | 17.844 | 5UTR |
| *Pdfr* | scaffold87\|size517236:83137 | SNP | T | A | 7.33E-15 | 6.96E-10 | 9.157 | 17.844 | 5UTR |
| *Pdfr* | scaffold87\|size517236:83209 | INDEL | CTG | C | 3.64E-15 | 4.05E-10 | 9.393 | 17.844 | 5UTR |
| *Pdfr* | scaffold87\|size517236:83223 | SNP | A | G | 3.64E-15 | 4.05E-10 | 9.393 | 17.844 | Enhancer |
| *Pdfr* | scaffold87\|size517236:83225 | SNP | T | C | 3.64E-15 | 4.05E-10 | 9.393 | 17.844 | Enhancer |
| *Pdfr* | scaffold87\|size517236:83242 | SNP | G | A | 3.64E-15 | 4.05E-10 | 9.393 | 17.844 | Enhancer |
| *Pdfr* | scaffold87\|size517236:83268 | SNP | G | A | 7.16E-16 | 1.08E-10 | 9.967 | 17.845 | Enhancer |
| *Pdfr* | scaffold87\|size517236:83272 | SNP | T | C | 2.58E-16 | 5.11E-11 | 10.292 | 17.845 | Enhancer |
| *Pdfr* | scaffold87\|size517236:83285 | SNP | A | T | 1.50E-15 | 2.02E-10 | 9.695 | 17.845 | Enhancer |
| *Pdfr* | scaffold87\|size517236:83286 | SNP | G | T | 1.50E-15 | 2.02E-10 | 9.695 | 17.845 | Enhancer |
| *Pdfr* | scaffold87\|size517236:83223 | SNP | A | G | 3.64E-15 | 4.05E-10 | 9.393 | 17.844 | Enhancer |
| *Pdfr* | scaffold87\|size517236:83225 | SNP | T | C | 3.64E-15 | 4.05E-10 | 9.393 | 17.844 | Enhancer |
| *Pdfr* | scaffold87\|size517236:83242 | SNP | G | A | 3.64E-15 | 4.05E-10 | 9.393 | 17.844 | Enhancer |
| *Pdfr* | scaffold87\|size517236:83268 | SNP | G | A | 7.16E-16 | 1.08E-10 | 9.967 | 17.845 | Enhancer |
| *Pdfr* | scaffold87\|size517236:83272 | SNP | T | C | 2.58E-16 | 5.11E-11 | 10.292 | 17.845 | Enhancer |
| *Pdfr* | scaffold87\|size517236:83285 | SNP | A | T | 1.50E-15 | 2.02E-10 | 9.695 | 17.845 | Enhancer |
| *Pdfr* | scaffold87\|size517236:83286 | SNP | G | T | 1.50E-15 | 2.02E-10 | 9.695 | 17.845 | Enhancer |
| *Pdfr* | scaffold87\|size517236:83288 | SNP | G | A | 1.50E-15 | 2.02E-10 | 9.695 | 17.845 | Enhancer |
| *Pdfr* | scaffold87\|size517236:83289 | INDEL | C | CTA | 1.98E-14 | 1.64E-09 | 8.785 | 17.845 | Enhancer |
| *Pdfr* | scaffold87\|size517236:83293 | INDEL | G | GC | 1.98E-14 | 1.64E-09 | 8.785 | 17.845 | Enhancer |
| *Pdfr* | scaffold87\|size517236:83321 | SNP | A | G | 1.98E-14 | 1.64E-09 | 8.785 | 17.845 | Enhancer |
| *Pdfr* | scaffold87\|size517236:83335 | SNP | G | A | 1.98E-14 | 1.64E-09 | 8.785 | 17.845 | Enhancer |
| *Pdfr* | scaffold87\|size517236:83361 | SNP | G | A | 3.53E-14 | 2.57E-09 | 8.590 | 17.845 | Enhancer |
| *Pdfr* | scaffold87\|size517236:83407 | INDEL | G | GTAAATTT | 4.21E-16 | 7.49E-11 | 10.126 | 17.845 | Enhancer |
| *Pdfr* | scaffold87\|size517236:83496 | SNP | G | A | 8.86E-16 | 1.29E-10 | 9.889 | 17.845 | Enhancer |
| *Pdfr* | scaffold87\|size517236:83514 | SNP | G | A | 2.83E-13 | 1.53E-08 | 7.815 | 17.845 | Enhancer |
| *Pdfr* | scaffold87\|size517236:83557 | SNP | G | C | 3.07E-14 | 2.32E-09 | 8.635 | 17.845 | Enhancer |
| *Pdfr* | scaffold87\|size517236:83618 | SNP | T | C | 5.92E-14 | 4.08E-09 | 8.389 | 17.845 | Enhancer |
| *Pdfr* | scaffold87\|size517236:83639 | SNP | A | C | 5.92E-14 | 4.08E-09 | 8.389 | 17.845 | Enhancer |
| *Pdfr* | scaffold87\|size517236:83653 | SNP | T | C | 5.92E-14 | 4.08E-09 | 8.389 | 17.845 | Enhancer |
| *Pdfr* | scaffold87\|size517236:83655 | SNP | A | C | 5.92E-14 | 4.08E-09 | 8.389 | 17.845 | Enhancer |
| *Pdfr* | scaffold87\|size517236:83731 | INDEL | A | ATAAT | 2.83E-13 | 1.53E-08 | 7.815 | 17.845 | Enhancer |
| *Pdfr* | scaffold87\|size517236:83782 | INDEL | ACCTGTACCACGGT | A | 1.47E-12 | 6.81E-08 | 7.167 | 17.845 | Enhancer |
| *Pdfr* | scaffold87\|size517236:83796 | SNP | C | A | 3.57E-17 | 1.09E-11 | 10.963 | 17.845 | Enhancer |
| *Pdfr* | scaffold87\|size517236:83798 | SNP | C | T | 1.16E-14 | 1.04E-09 | 8.983 | 17.845 | Enhancer |
| *Pdfr* | scaffold87\|size517236:83834 | INDEL | AC | A | 2.47E-17 | 8.27E-12 | 11.082 | 17.845 | Enhancer |
| *Pdfr* | scaffold87\|size517236:83875 | INDEL | G | GT | 7.12E-16 | 1.07E-10 | 9.971 | 17.845 | Enhancer |
| *Pdfr* | scaffold87\|size517236:83903 | SNP | T | C | 7.12E-16 | 1.07E-10 | 9.971 | 17.845 | Enhancer |
| *Pdfr* | scaffold87\|size517236:83918 | SNP | A | G | 7.12E-16 | 1.07E-10 | 9.971 | 17.845 | Enhancer |
| *Pdfr* | scaffold87\|size517236:83931 | INDEL | G | GCTTAACAGC | 7.12E-16 | 1.07E-10 | 9.971 | 17.845 | Enhancer |
| *Pdfr* | scaffold87\|size517236:83953 | INDEL | T | TACGC | 7.13E-17 | 1.85E-11 | 10.733 | 17.845 | Enhancer |
| *Pdfr* | scaffold87\|size517236:84001 | SNP | G | A | 7.12E-16 | 1.07E-10 | 9.971 | 17.845 | Enhancer |
| *Pdfr* | scaffold87\|size517236:84010 | INDEL | AT | A | 4.00E-18 | 2.59E-12 | 11.587 | 17.845 | Enhancer |
| *Pdfr* | scaffold87\|size517236:84012 | INDEL | AT | A | 1.63E-18 | 1.31E-12 | 11.883 | 17.845 | Enhancer |
| *Pdfr* | scaffold87\|size517236:84013 | INDEL | CG | C | 7.12E-16 | 1.07E-10 | 9.971 | 17.845 | Enhancer |
| *Pdfr* | scaffold87\|size517236:84017 | SNP | T | A | 7.12E-16 | 1.07E-10 | 9.971 | 17.845 | Enhancer |
| *Pdfr* | scaffold87\|size517236:84054 | INDEL | GT | AT | 7.12E-16 | 1.07E-10 | 9.971 | 17.845 | Enhancer |
| *Pdfr* | scaffold87\|size517236:84055 | SNP | A | T | 7.12E-16 | 1.07E-10 | 9.971 | 17.845 | Enhancer |
| *Pdfr* | scaffold87\|size517236:84059 | SNP | C | A | 7.12E-16 | 1.07E-10 | 9.971 | 17.845 | Enhancer |
| *Pdfr* | scaffold87\|size517236:84083 | SNP | A | G | 7.12E-16 | 1.07E-10 | 9.971 | 17.845 | Enhancer |
| *Pdfr* | scaffold87\|size517236:84088 | SNP | T | G | 6.03E-15 | 6.03E-10 | 9.220 | 17.845 | Enhancer |
| *Pdfr* | scaffold87\|size517236:84099 | SNP | A | G | 6.03E-15 | 6.03E-10 | 9.220 | 17.845 | Enhancer |
| *Pdfr* | scaffold87\|size517236:84114 | SNP | A | T | 6.03E-15 | 6.03E-10 | 9.220 | 17.845 | Enhancer |
| *Pdfr* | scaffold87\|size517236:84156 | SNP | T | C | 5.92E-14 | 4.08E-09 | 8.389 | 17.845 | Enhancer |
| *Pdfr* | scaffold87\|size517236:84237 | INDEL | A | ACATCAT | 9.93E-12 | 3.88E-07 | 6.411 | 17.845 | Enhancer |
| *Pdfr* | scaffold87\|size517236:84320 | SNP | G | A | 1.80E-13 | 1.03E-08 | 7.987 | 17.846 | Enhancer |
| *Pdfr* | scaffold87\|size517236:84327 | SNP | G | A | 1.80E-13 | 1.03E-08 | 7.987 | 17.846 | Enhancer |
| *Pdfr* | scaffold87\|size517236:84370 | SNP | A | G | 7.33E-15 | 6.96E-10 | 9.157 | 17.846 | Enhancer |
| *Pdfr* | scaffold87\|size517236:84404 | SNP | A | G | 7.33E-15 | 6.96E-10 | 9.157 | 17.846 | Enhancer |
| *Pdfr* | scaffold87\|size517236:84446 | INDEL | T | TA | 4.74E-17 | 1.34E-11 | 10.873 | 17.846 | Enhancer |
| *Pdfr* | scaffold87\|size517236:84479 | SNP | G | A | 7.33E-15 | 6.96E-10 | 9.157 | 17.846 | Enhancer |
| *Pdfr* | scaffold87\|size517236:84506 | SNP | C | A | 7.33E-15 | 6.96E-10 | 9.157 | 17.846 | Enhancer |
| *Pdfr* | scaffold87\|size517236:84538 | SNP | G | A | 3.53E-14 | 2.57E-09 | 8.590 | 17.846 | Enhancer |
| *Pdfr* | scaffold87\|size517236:84567 | SNP | G | A | 9.96E-16 | 1.42E-10 | 9.848 | 17.846 | Enhancer |
| *Pdfr* | scaffold87\|size517236:84691 | SNP | T | G | 3.64E-15 | 4.05E-10 | 9.393 | 17.846 | Enhancer |
| *Pdfr* | scaffold87\|size517236:84726 | SNP | G | C | 3.64E-15 | 4.05E-10 | 9.393 | 17.846 | Enhancer |
| *Pdfr* | scaffold87\|size517236:84795 | SNP | G | C | 7.33E-15 | 6.96E-10 | 9.157 | 17.846 | Enhancer |
| *Pdfr* | scaffold87\|size517236:84808 | SNP | C | A | 3.53E-14 | 2.57E-09 | 8.590 | 17.846 | Enhancer |
| *Pdfr* | scaffold87\|size517236:84812 | SNP | C | T | 3.53E-14 | 2.57E-09 | 8.590 | 17.846 | Enhancer |
| *Pdfr* | scaffold87\|size517236:84936 | INDEL | TATA | T | 2.83E-13 | 1.53E-08 | 7.815 | 17.846 | Enhancer |
| *Pdfr* | scaffold87\|size517236:84949 | INDEL | GGAGCTTAATATTGATAGAAATA | G | 1.42E-14 | 1.24E-09 | 8.907 | 17.846 | Enhancer |
| *Pdfr* | scaffold87\|size517236:85255 | SNP | G | A | 1.07E-09 | 2.22E-05 | 4.654 | 17.847 | Enhancer |
| *Pdfr* | scaffold87\|size517236:85262 | SNP | T | C | 9.73E-08 | 0.00078195 | 3.107 | 17.847 | Enhancer |

| **Gene** | **SNP** | **Type** | **Major Allele** | **Minor Alllele** | **P** | **fdr** | **LogQ** | **Mb** | **Location** |
|---|---|---|---|---|---|---|---|---|---|
| *Pdfr* | scaffold87\|size517236:85363 | SNP | T | C | 6.94E-12 | 2.80E-07 | 6.553 | 17.847 | Enhancer |
| *Pdfr* | scaffold87\|size517236:85365 | INDEL | A | ATAGGTACC | 2.98E-14 | 2.27E-09 | 8.644 | 17.847 | Enhancer |
| *Pdfr* | scaffold87\|size517236:85370 | INDEL | * | T | 1.19E-13 | 7.43E-09 | 8.129 | 17.847 | Enhancer |
| *Pdfr* | scaffold87\|size517236:85417 | SNP | T | C | 4.74E-17 | 1.34E-11 | 10.873 | 17.847 | Enhancer |
| *Pdfr* | scaffold87\|size517236:86427 | SNP | G | T | 4.21E-16 | 7.49E-11 | 10.126 | 17.848 | Enhancer |
| *Pdfr* | scaffold87\|size517236:86641 | INDEL | TTATGC | T | 7.12E-16 | 1.07E-10 | 9.971 | 17.848 | Enhancer |
| *Pdfr* | scaffold87\|size517236:87232 | INDEL | AATAATT | A | 4.00E-18 | 2.59E-12 | 11.587 | 17.848 | Enhancer |
| *Pdfr* | scaffold87\|size517236:87271 | SNP | T | A | 7.12E-16 | 1.07E-10 | 9.971 | 17.849 | Enhancer |
| *Pdfr* | scaffold87\|size517236:87516 | SNP | T | C | 4.74E-17 | 1.34E-11 | 10.873 | 17.849 | Enhancer |
| *Pdfr* | scaffold87\|size517236:87684 | INDEL | 745 bp deletion | T | 4.74E-17 | 1.34E-11 | 10.873 | 17.849 | Enhancer |
| *Pdfr* | scaffold87\|size517236:88637 | SNP | T | C | 1.32E-14 | 1.17E-09 | 8.932 | 17.850 | Enhancer |
| *Pdfr* | scaffold87\|size517236:88794 | SNP | G | A | 4.21E-16 | 7.49E-11 | 10.126 | 17.850 | Enhancer |
| *Pdfr* | scaffold87\|size517236:88818 | INDEL | G | GTACC | 4.21E-16 | 7.49E-11 | 10.126 | 17.850 | Enhancer |
| *Pdfr* | scaffold87\|size517236:88838 | SNP | C | A | 2.29E-15 | 2.83E-10 | 9.548 | 17.850 | Enhancer |
| *Pdfr* | scaffold87\|size517236:88864 | SNP | C | T | 5.18E-14 | 3.62E-09 | 8.441 | 17.850 | Enhancer |
| *Pdfr* | scaffold87\|size517236:89791 | INDEL | G | GA | 4.21E-16 | 7.49E-11 | 10.126 | 17.851 | Enhancer |
| *Pdfr* | scaffold87\|size517236:89861 | SNP | A | T | 1.32E-14 | 1.17E-09 | 8.932 | 17.851 | Enhancer |
| *Pdfr* | scaffold87\|size517236:89863 | INDEL | AGGTACC | A | 3.25E-12 | 1.40E-07 | 6.854 | 17.851 | Enhancer |
| *Pdfr* | scaffold87\|size517236:89865 | SNP | A | T | 2.29E-15 | 2.83E-10 | 9.548 | 17.851 | Enhancer |
| *Pdfr* | scaffold87\|size517236:89866 | SNP | A | T | 1.32E-14 | 1.17E-09 | 8.932 | 17.851 | Enhancer |
| *Pdfr* | scaffold87\|size517236:89869 | SNP | A | G | 2.29E-15 | 2.83E-10 | 9.548 | 17.851 | Enhancer |
| *Pdfr* | scaffold87\|size517236:89883 | INDEL | GTT | G | 1.50E-15 | 2.02E-10 | 9.695 | 17.851 | Enhancer |
| *Pdfr* | scaffold87\|size517236:89906 | SNP | G | A | 2.58E-16 | 5.11E-11 | 10.292 | 17.851 | Enhancer |
| *Pdfr* | scaffold87\|size517236:89908 | SNP | C | T | 2.58E-16 | 5.11E-11 | 10.292 | 17.851 | Enhancer |
| *Pdfr* | scaffold87\|size517236:89941 | INDEL | T | TA | 2.58E-16 | 5.11E-11 | 10.292 | 17.851 | Enhancer |
| *Pdfr* | scaffold87\|size517236:89943 | INDEL | T | TTTTTA | 2.58E-16 | 5.11E-11 | 10.292 | 17.851 | Enhancer |
| *Pdfr* | scaffold87\|size517236:89951 | INDEL | C | CCTCT | 2.58E-16 | 5.11E-11 | 10.292 | 17.851 | Enhancer |
| *Pdfr* | scaffold87\|size517236:89957 | INDEL | C | CG | 2.58E-16 | 5.11E-11 | 10.292 | 17.851 | Enhancer |
| *Pdfr* | scaffold87\|size517236:89987 | SNP | A | C | 4.21E-16 | 7.49E-11 | 10.126 | 17.851 | Enhancer |
| *Pdfr* | scaffold87\|size517236:90028 | SNP | A | C | 7.33E-15 | 6.96E-10 | 9.157 | 17.851 | Enhancer |
| *Pdfr* | scaffold87\|size517236:90174 | SNP | A | AT | 9.43E-18 | 4.29E-12 | 11.368 | 17.851 | Enhancer |
| *Pdfr* | scaffold87\|size517236:90179 | SNP | TCA | T | 7.12E-16 | 1.07E-10 | 9.971 | 17.851 | Enhancer |
| *Pdfr* | scaffold87\|size517236:90224 | INDEL | TTTCTTC | T | 5.47E-16 | 9.02E-11 | 10.045 | 17.851 | Enhancer |
| *Pdfr* | scaffold87\|size517236:90224 | INVERSION | Inverted 400 kb | Collinear | 6.25E-14 | 4.28E-09 | 8.369 | 17.852 | Enhancer |
| *Pdfr* | scaffold87\|size517236:90356 | INDEL | A | AGTGCGGTTTTCACCTGGTCAGTCCAGCGCATGGGCGACCTGCCGCGCGCTCTT | 2.17E-13 | 1.19E-08 | 7.924 | 17.852 | Enhancer |
| *Pdfr* | scaffold87\|size517236:90420 | INDEL | C | CCTG | 8.21E-14 | 5.43E-09 | 8.265 | 17.852 | Enhancer |
| *Pdfr* | scaffold87\|size517236:90458 | SNP | G | T | 3.92E-16 | 7.11E-11 | 10.148 | 17.852 | Enhancer |
| *Pdfr* | scaffold87\|size517236:90473 | SNP | A | T | 2.98E-14 | 2.27E-09 | 8.644 | 17.852 | Enhancer |
| *Pdfr* | scaffold87\|size517236:90524 | SNP | T | C | 2.16E-13 | 1.19E-08 | 7.924 | 17.852 | Enhancer |
| *Pdfr* | scaffold87\|size517236:90539 | SNP | C | G | 3.66E-15 | 4.07E-10 | 9.390 | 17.852 | Enhancer |
| *Pdfr* | scaffold87\|size517236:90578 | SNP | T | G | 2.62E-12 | 1.15E-07 | 6.939 | 17.852 | Enhancer |
| *Pdfr* | scaffold87\|size517236:90625 | SNP | A | G | 1.63E-18 | 1.31E-12 | 11.883 | 17.852 | Enhancer |
| *Pdfr* | scaffold87\|size517236:90851 | SNP | C | T | 3.92E-16 | 7.11E-11 | 10.148 | 17.852 | Enhancer |
| *Pdfr* | scaffold87\|size517236:90867 | SNP | T | C | 5.47E-16 | 9.02E-11 | 10.045 | 17.852 | Enhancer |
| *Pdfr* | scaffold87\|size517236:90891 | SNP | T | C | 5.86E-17 | 1.59E-11 | 10.799 | 17.852 | Enhancer |
| *Pdfr* | scaffold87\|size517236:90900 | SNP | T | C | 5.47E-16 | 9.02E-11 | 10.045 | 17.852 | Enhancer |
| *Pdfr* | scaffold87\|size517236:90925 | SNP | T | A | 3.66E-15 | 4.07E-10 | 9.390 | 17.852 | Enhancer |
| *Pdfr* | scaffold87\|size517236:90930 | SNP | T | A | 3.66E-15 | 4.07E-10 | 9.390 | 17.852 | Enhancer |
| *Pdfr* | scaffold87\|size517236:90931 | INDEL | CCATA | C | 3.66E-15 | 4.07E-10 | 9.390 | 17.852 | Enhancer |
| *Pdfr* | scaffold87\|size517236:90951 | SNP | T | A | 2.11E-13 | 1.17E-08 | 7.932 | 17.852 | Enhancer |
| *Pdfr* | scaffold87\|size517236:90959 | SNP | G | A | 2.11E-13 | 1.17E-08 | 7.932 | 17.852 | Enhancer |
| *Pdfr* | scaffold87\|size517236:90963 | INDEL | A | ATCTT | 2.11E-13 | 1.17E-08 | 7.932 | 17.852 | Enhancer |
| *Pdfr* | scaffold87\|size517236:90971 | INDEL | T | TAATTTTAATTA | 2.11E-13 | 1.17E-08 | 7.932 | 17.852 | Enhancer |
| *Pdfr* | scaffold87\|size517236:90983 | INDEL | C | CGAGTATCTTA | 2.11E-13 | 1.17E-08 | 7.932 | 17.852 | Enhancer |
| *Pdfr* | scaffold87\|size517236:90995 | SNP | A | T | 2.11E-13 | 1.17E-08 | 7.932 | 17.852 | Enhancer |
| *Pdfr* | scaffold87\|size517236:91001 | SNP | G | A | 2.11E-13 | 1.17E-08 | 7.932 | 17.852 | Enhancer |
| *Pdfr* | scaffold87\|size517236:91004 | SNP | A | G | 2.11E-13 | 1.17E-08 | 7.932 | 17.852 | Enhancer |
| *Pdfr* | scaffold87\|size517236:91017 | SNP | C | A | 1.18E-08 | 0.00014823 | 3.829 | 17.852 | Enhancer |
| *Pdfr* | scaffold87\|size517236:91037 | INDEL | A | ATAATAAT | 2.48E-11 | 9.00E-07 | 6.046 | 17.852 | Enhancer |
| *Pdfr* | scaffold87\|size517236:91064 | INDEL | G | GC | 1.51E-07 | 0.00111119 | 2.954 | 17.852 | Enhancer |
| *Pdfr* | scaffold87\|size517236:91066 | INDEL | TGTGAGTGTGAATG | T | 1.51E-07 | 0.00111119 | 2.954 | 17.852 | Enhancer |
| *Pdfr* | scaffold87\|size517236:91703 | INDEL | <DEL> | T | 7.82E-13 | 3.85E-08 | 7.415 | 17.853 | Enhancer |
| *Pdfr* | scaffold87\|size517236:91707 | SNP | C | T | 2.17E-13 | 1.19E-08 | 7.924 | 17.853 | Enhancer |
| *Pdfr* | scaffold87\|size517236:91715 | SNP | C | A | 2.98E-14 | 2.27E-09 | 8.644 | 17.853 | Enhancer |
| *Pdfr* | scaffold87\|size517236:91719 | INDEL | A | ATTAT | 2.98E-14 | 2.27E-09 | 8.644 | 17.853 | Enhancer |
| *Pdfr* | scaffold87\|size517236:91730 | INDEL | T | TGA | 2.98E-14 | 2.27E-09 | 8.644 | 17.853 | Enhancer |
| *Pdfr* | scaffold87\|size517236:91738 | INDEL | T | TCTAATTACCTTCAATTAATATTTTTA | 3.23E-14 | 2.41E-09 | 8.618 | 17.853 | Enhancer |
| *Pdfr* | scaffold87\|size517236:91758 | INDEL | <DEL> | A | 1.33E-13 | 8.11E-09 | 8.091 | 17.853 | Enhancer |
| *Pdfr* | scaffold87\|size517236:91771 | INDEL | T | TAA | 9.26E-15 | 8.55E-10 | 9.068 | 17.853 | Enhancer |
| *Pdfr* | scaffold87\|size517236:91775 | SNP | T | G | 9.26E-15 | 8.55E-10 | 9.068 | 17.853 | Enhancer |
| *Pdfr* | scaffold87\|size517236:91777 | SNP | T | A | 9.26E-15 | 8.55E-10 | 9.068 | 17.853 | Enhancer |
| *Pdfr* | scaffold87\|size517236:91818 | INDEL | CGA | GGA | 2.59E-16 | 5.13E-11 | 10.290 | 17.853 | Enhancer |
| *Pdfr* | scaffold87\|size517236:91820 | INDEL | GTTT | ATTT | 1.47E-12 | 6.81E-08 | 7.167 | 17.853 | Enhancer |

**Table S11. Data plotted in Figure S3**

| Citation | Location | Province/State | Type | PDD (days) | SE | Temp (ºC) | Sample Size | Origin |
|---|---|---|---|---|---|---|---|---|
| McLeod 1976 (31) | London | ON | Uni | 35.1 | 1.05 | 30 | NA | Field |
| McLeod 1976 (31) | Guelph | ON | Uni | 34.9 | 0.96 | 30 | NA | Field |
| McLeod 1976 (31) | Harrow | ON | Bi | 22.9 | 1.08 | 30 | NA | Field |
| McLeod 1976 (31) | Wallacetown | ON | Bi | 25.9 | 1.34 | 30 | NA | Field |
| McLeod 1979 (32) | St. Remi | QC | Uni | 31.25 | 1.3 | 30 | 40 | Lab colony |
| McLeod 1979 (32) | St. Remi | QC | Bi | 11 | 0.82 | 30 | 10 | Field |
| Glover et al. 1992 (33) | Bouckville | NY | Uni | 43.68 | 1.21 | 30 | 126 | ~1 year in lab |
| Glover et al. 1992 (33) | Geneva | NY | Bi | 15.3 | 0.45 | 30 | 77 | ~1 year in lab |
| Dopman 2005 (34) | Bouckville | NY | Uni | 26.31 | 0.84 | 26 | 133 | Field |
| Dopman 2005 (34) | Eden | NY | Bi | 12.57 | 1.05 | 26 | 63 | Field |
| Kozak & Dopman unpub | Hurley | NY | Bi | 14.30 | 0.81 | 26 | 44 | Field |
| Wadsworth & Dopman unpub | Dover | MA | Bi | 18.81 | 1.08 | 26 | 54 | F1 generation in lab |
| Hamilton & Harrison unpub | Union Springs | NY | Uni | 37.49 | 0.80 | 26 | 108 | Field |
